# Coding-sequence evolution does not explain divergence in petal anthocyanin pigmentation between *Mimulus luteus* var. *luteus* and *M. l. variegatus*

**DOI:** 10.1101/2022.12.11.519905

**Authors:** Walker E. Orr, Ji Yang Kim, Tejas Raj, Ellen K. Hom, Ashley E. Person, Anne Vonada, John A. Stratton, Arielle M. Cooley

**Affiliations:** Whitman College Biology Department, Walla Walla WA, 99362, U.S.A; Whitman College Computer Science Department, Walla Walla WA, 99362, U.S.A

**Keywords:** anthocyanin regulation, floral color patterning, gene expression, *cis*-regulatory evolution, R2R3 MYBs, *Mimulus*, petal lobe pigmentation, post-transcriptional mRNA editing, A- to-I editing, transient transformation, digital image analysis

## Abstract

Phenotypic transitions in related taxa often share a common genetic basis, which suggests that there are constraints that shape the process of evolution at the genetic level. For example, noncoding changes in a gene might be favored relative to coding changes due to being less constrained by pleiotropic effects. Here we evaluate the importance of coding-sequence changes to the recent evolution of a novel anthocyanin pigmentation trait in the monkeyflower genus *Mimulus*. The magenta-flowered *Mimulus luteus* var. *variegatus* recently gained petal lobe anthocyanin pigmentation via a single-locus Mendelian difference from its sister taxon, the yellow-flowered *M. l. luteus*. Previous work showed that the differentially expressed transcription factor gene *MYB5a/NEGAN* is the single causal gene. However, it was not clear whether *MYB5a* coding-sequence evolution (in addition to the observed patterns of differential expression) might also have contributed to increased anthocyanin production in *M. l. variegatus*. Quantitative image analysis of tobacco leaves, transfected with highly expressed *MYB5a* coding sequence from each taxon, revealed robust anthocyanin production driven by both alleles compared to a negative control. Counter to expectations, significantly higher anthocyanin production was driven by the coding sequence from the low-anthocyanin taxon *M. l. luteus*. Together with previously-published expression studies, this supports the hypothesis that petal pigment in *M. l. variegatus* was not gained by protein-coding changes, but instead via non-coding *cis*-regulatory evolution. Finally, while constructing the transgenes needed for this experiment, we unexpectedly discovered two sites in *MYB5a* that appear to be post-transcriptionally edited – a phenomenon that has been rarely reported, and even less often explored, for nuclear-encoded plant mRNAs.

## Introduction

To what extent are the molecular mechanisms of evolutionary diversification predictable? Biologists have long been interested in understanding constraints on the evolutionary process, which can increase our ability to predict the molecular mechanisms that underlie a specific trait.

For example, coding and noncoding mutations have been hypothesized to differ in their contributions to evolutionary change (Hoekstra and Coyne, 2007; Stern and Orgogozo, 2008). Noncoding “*cis*-regulatory” regions integrate upstream signals to determine the conditions under which a protein will be expressed. These regions tend to be modular, so mutations can easily alter the gene expression driven by one *cis*-regulatory region without altering expression patterns driven by neighboring regions (Wray, 2007). By contrast, a change in a gene’s coding sequence tends to be more pleiotropic since it will likely alter the gene product’s function in all contexts under which that gene is expressed. Stern and Orgogozo (2008) proposed a model predicting that the contribution of coding changes to phenotypic transitions will decline with evolutionary distance between the diverged taxa, due to lower average fitness values for coding changes compared to noncoding changes. The pattern has been documented empirically by Wittkopp et al. (2008) in *Drosophila*.

In plants, transitions in anthocyanin pigmentation are especially well suited to investigating the mechanisms of molecular evolution (Davies et al., 2012; Sobel and Streisfeld, 2013; LaFountain and Yuan, 2021; Li et al., 2022). Anthocyanins are a class of flavonoids that are responsible for purple, pink, and red colors in diverse angiosperm tissues, including leaves, seed coats, and flowers (Durbin et al., 2003). In the model plant *Arabidopsis thaliana* and many other species, the anthocyanin pathway consists of “early” genes *CHS, CHI,* and *F3’H*, followed by the “late” genes *DFR, ANS,* and *UF3GT* (Dubos et al., 2010). The “late” genes are coordinately activated by MYB-bHLH-WD40 (MBW) transcription factor complexes (Grotewold, 2006). Much of the target specificity of an MBW complex is conferred by the MYB partner in the complex: different tissues make use of different *MYB* genes to stimulate anthocyanin production, but may use the same *bHLH* gene (Quattrocchio et al., 2006). *MYB* genes thus serve as “input-output” integrators that regulate gene batteries under highly specific contexts. Consistent with their position in gene networks and the hypothesized importance of “input-output” genes in repeated evolution events, *MYB* genes demonstrate remarkable reuse in floral pigment transitions across a variety of species (Streisfeld and Rausher, 2011; Yuan et al., 2013).

The independent evolution of petal lobe anthocyanin (PLA) pigmentation in three lineages in the *luteus* group of the monkeyflower genus *Mimulus* (synonym *Erythranthe*; see Barker (2012) and Lowry et al. (2019) for a debate within the monkeyflower research community over which genus name to use) provides an opportunity to test the relative importance of coding and noncoding changes at different evolutionary time scales. *Mimulus* lends itself to studies in plant evo-devo, thanks to a diversity of species with short generation times, high fecundity, amenability to greenhouse cultivation, and a large range of environmental adaptation (Wu et al., 2008; Yuan 2018). Sequenced genomes have been published for several species including *M. guttatus* (Puzey et al., 2017) and *M. l. luteus* (Edger et al., 2017).

The *luteus* group of *Mimulus* has an ancestral phenotype of yellow, carotenoid-pigmented flowers with red, anthocyanin-pigmented spots in the nectar guide region (Fig. 1). The overall color of the anthocyanin-pigmented petal tissue ranges from orange in *M. cupreus*, to red in *M. l. luteus*, to magenta in *M. l. variegatus* and *M. naiandinus*, depending on the relative intensity of carotenoid and anthocyanin pigmentation (Fig. 1).

**Fig. 1.**
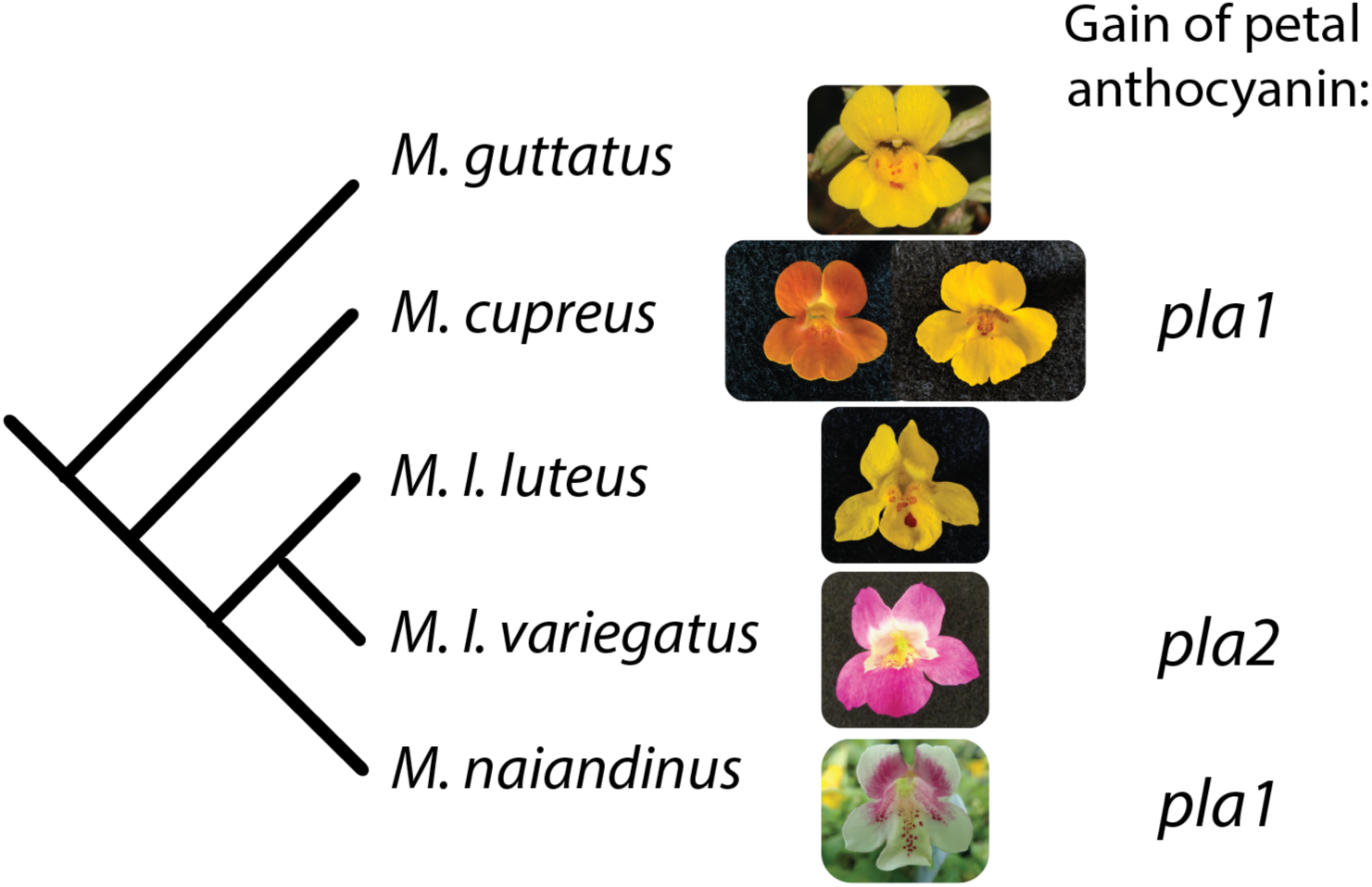
Petal lobe anthocyanin has been gained repeatedly in the *luteus* group of *Mimulus*. Closely related species outside the *luteus* group, like *M. guttatus*, are typically yellow-flowered, with red anthocyanin pigmentation restricted to the nectar guide region of the corolla. *Mimulus cupreus* and *M. naiandinus* have each gained petal lobe anthocyanin via a single-locus change at genomic region *pla1*, while the magenta- petaled *M. luteus* var. *variegatus* gained petal lobe anthocyanin via a change at *pla2*. A rare yellow-flowered morph of *M. cupreus*, found in a single population in Chile, has lost petal lobe anthocyanin via a change at *pla1*. Figure modified from Zheng et al. (2021).

In three members of the *luteus* group—the magenta-flowered *M. l. variegatus* and *M. naiandinus,* and the orange-flowered *M. cupreus*—anthocyanin pigment has expanded into the petal lobes. The gain of PLA is a derived, single-locus Mendelian trait in all three taxa. In *M. cupreus* and *M. naiandinus*, this is controlled by the *pla1* locus, which contains candidate anthocyanin-activating transcription factor genes *MYB2* and *MYB3a* (Cooley et al., 2011). A rare yellow-flowered morph of *M. cupreus* does not bear petal lobe anthocyanins (PLA). Since it is found in a single population in Chile, intermingled with orange morphs of *M. cupreus* (Cooley et al., 2008), it likely represents a secondary loss of the PLA trait, and also segregates as a single-locus trait mapping to *pla1*. In *M. l. variegatus*, in contrast, the gain of PLA is conferred by an unlinked second locus named *pla2*. The *pla2* locus contains candidate gene *MYB5a/NEGAN*. All of the candidate genes at *pla1* and *pla2* belong to subgroup 6 of the R2R3 MYB gene family (Cooley et al., 2011), which regulate the ”late” anthocyanin biosynthetic genes via participation in the MBW complex (reviewed in Lloyd et al. (2017); see Stracke et al. (2001) for a description of the subgroup 6 genes).

While transitions in floral pigment traits have been extensively studied in plants, pigment losses have been investigated more often than pigment gains (Rausher, 2008). Studying the repeated gain of PLA in the *luteus* group presents an opportunity to rectify this. PLA transitions in the group also enable a test of the Stern and Orgogozo (2008) prediction that protein-coding changes are relatively more important within populations, while noncoding changes become increasingly abundant as evolutionary divergence increases. Consistent with this prediction, the rare yellow morph of *M. cupreus* - a within-population polymorphism linked to *pla1* (Cooley and Willis 2009; Cooley et al. 2011) - appears to be associated with a deletion or other major mutation in exon 3 or 4 of *MYB2* (Supplemental Figures S1-S4). Based on the Stern and Orgogozo (2008) model, we expect that the fixed gains of PLA in *M. l. variegatus*, orange *M. cupreus*, and *M. naiandinus*, are more likely to be caused by *cis*-regulatory evolution.

Of the three taxa that recently gained PLA, *M. l. variegatus* is the most thoroughly characterized. In *M. l. variegatus,* a combination of genetic mapping, *MYB5a* RNAi and overexpression, and transcriptomic studies of wild-type versus *MYB5a* RNAi lines, shows that *MYB5a* is both necessary and sufficient for the gain of PLA (Cooley et al. 2011; Zheng et al., 2021), and points to one particular splice variant of the gene as participating in anthocyanin activation.

Unlike most *MYB* genes, which have a conserved three-exon structure (Dubos et al., 2010), *MYB5a* in *M. l. luteus* and *M. l. variegatus* has four exons. The exon 1-2-4 splice variant, but not the 1-2-3 variant, was found to be abundant specifically in anthocyanin-pigmented petal tissue: the petal lobes of *M. l. variegatus* and the nectar guide regions of both taxa (Zheng et al. 2021). Knockdown of the 1-2-4 splice variant resulted in the loss of petal pigmentation from *M. l. variegatus* (Zheng et al., 2021). Finally, the exon 1-2-4 splice variant of *M. l. variegatus* - but not the 1-2-3 splice variant - contains the subgroup 6 motif found in all known R2R3 MYB anthocyanin activators (Stracke et al., 2001). Thus, the exon 1-2-4 splice variant of *MYB5a* appears to be responsible for the gain of petal lobe anthocyanins in *M. l. variegatus*. While its pattern of spatially specific expression suggests a mechanism of *cis*-regulatory evolution, it is unknown whether divergence in the protein-coding sequence of *MYB5a* additionally contributed to the evolution of increased anthocyanin production in *M. l. variegatus*.

In previous studies (Zheng et al., 2021), we relied on the stable transformation procedure published by Yuan et. al. (2013) to test hypotheses about *MYB* gene function. While the method is capable of producing stably transformed offspring, it has the disadvantage of having a low transformation efficiency (about one seed per thousand in *M. l. variegatus*) and a large time cost of about five months between infiltration of plants and flowering of transformant offspring. Transient transformation is an attractive alternative for more rapid tests of gene function, particularly for genes - such as pigment activators - that are expected to produce an easily visible phenotype. In transient expression, the transgene is delivered to plant cells and transcribed by the plant’s transcriptional machinery without necessarily being incorporated into the plant’s genome, and transient transformation is regularly used in the tobacco genus *Nicotiana* (Kapila et al., 1997; Schöb et al., 1997; Yang et al. 2000; Sparkes 2006). *Nicotiana* is relatively closely related to *Mimulus*, as the two genera belong to the sister orders of Solanales and Lamiales, respectively, and *Nicotiana* is a highly tractable system for transgenic experimentation. It is routinely used for heterologous genetic experiments *in planta*, including tests of flower color genes from both rosids and asterids (Montefiori et al., 2015; Tian et al., 2017). Ding and Yuan (2016) adapted methods from *Nicotiana* for use in *Mimulus lewisii*. In our hands, however, transient transformation caused substantial leaf tissue death in both *M. lewisii* and *M. l. luteus*. We therefore returned to *Nicotiana*, using *N. tabacum* as the host for transient tests of *MYB5a* gene function.

If the gain of petal lobe anthocyanin (PLA) in the magenta-flowered *M. l. variegatus* is caused solely by a *cis*-regulatory-driven spatial expansion of *MYB5a* function, then we predict that the coding sequences of *MYB5a* from *M. l. variegatus* and the yellow-flowered *M. l. luteus* will be equally capable of stimulating anthocyanin production. We tested this hypothesis by transiently expressing each taxon’s *MYB5a* exon 1-2-4 sequence in leaves of *N. tabacum*. We used a custom image analysis pipeline to rapidly generate quantitative estimates of pigment production in the transformed leaves.

Somewhat surprisingly, the *luteus* allele of *MYB5a* drove significantly stronger anthocyanin pigmentation than did the *variegatus* allele. This indicates that *M. l. variegatus* did not evolve its greater anthocyanin pigmentation via coding-sequence changes at *MYB5a*, and points to *cis*- regulatory evolution at the causal *MYB5a* gene as a more likely driver of the gain of petal lobe anthocyanin pigmentation in *M. l. variegatus*.

## Methods

### Plant materials and growth conditions

*Nicotiana tabacum* cv. Petit Havana SR1 seeds were obtained from Lehle Seeds (Round Rock, TX, USA). *Mimulus lewisii*, line LF10HT1 (two generations inbred) seeds were a gift from the Yao-Wu Yuan lab at the University of Connecticut (Storrs, CT, USA). *Mimulus luteus* var. *luteus* and *M. l. variegatus* were originally collected in Chile from the El Yeso and Río Cipreses populations, respectively (Cooley et al. 2008), and self-fertilized repeatedly with single-seed descent to generate highly inbred lines. In this work, we utilized the 12-generations inbred line

### *M. l. luteus* EY7 and the 11-generations inbred line *M. l. variegatus* RC6

Seeds were surface-planted on wet soil and grown at Whitman College (Walla Walla, WA, USA) in a greenhouse with 16-hour day lengths and temperatures ranging from 15°C to 30°C. Plants were misted daily and fertilized three times per week with Open Sesame flowering fertilizer (Fox Farm, Samoa, CA, USA).

### Determining the genomically encoded sequence of MYB5a from M. l. variegatus

While investigating the *MYB5a* protein-coding regions of *M. l. luteus* and *M. l. variegatus*, using the cloning and sequencing methods described in Zheng et al. (2021), we discovered an unexpected new sequence in the fourth exon of *M. l. variegatus Myb5a*. In two different sequencing reactions, originating from two distinct cDNA syntheses from a single mRNA extraction from young bud *M. l. variegatus* tissue, a “GG” variant was found in which adenines at positions 582 and 684 of the exon 1-2-4 splice variant were replaced with guanines (Figure 2). A third sequencing reaction from the same mRNA extraction produced the expected “AA” sequence that was previously reported in Zheng et al. (2021). Because the same two variants were observed, twice each, in the same two samples, we discounted the likelihood of four random sequencing errors. Instead, we hypothesized that the “GG” variant might be encoded in the genome of *M. l. variegatus*, perhaps representing an alternate allele of *MYB5a* or a closely related gene duplicate, or that it might be the result of post-transcriptional editing.

**Figure 2.**
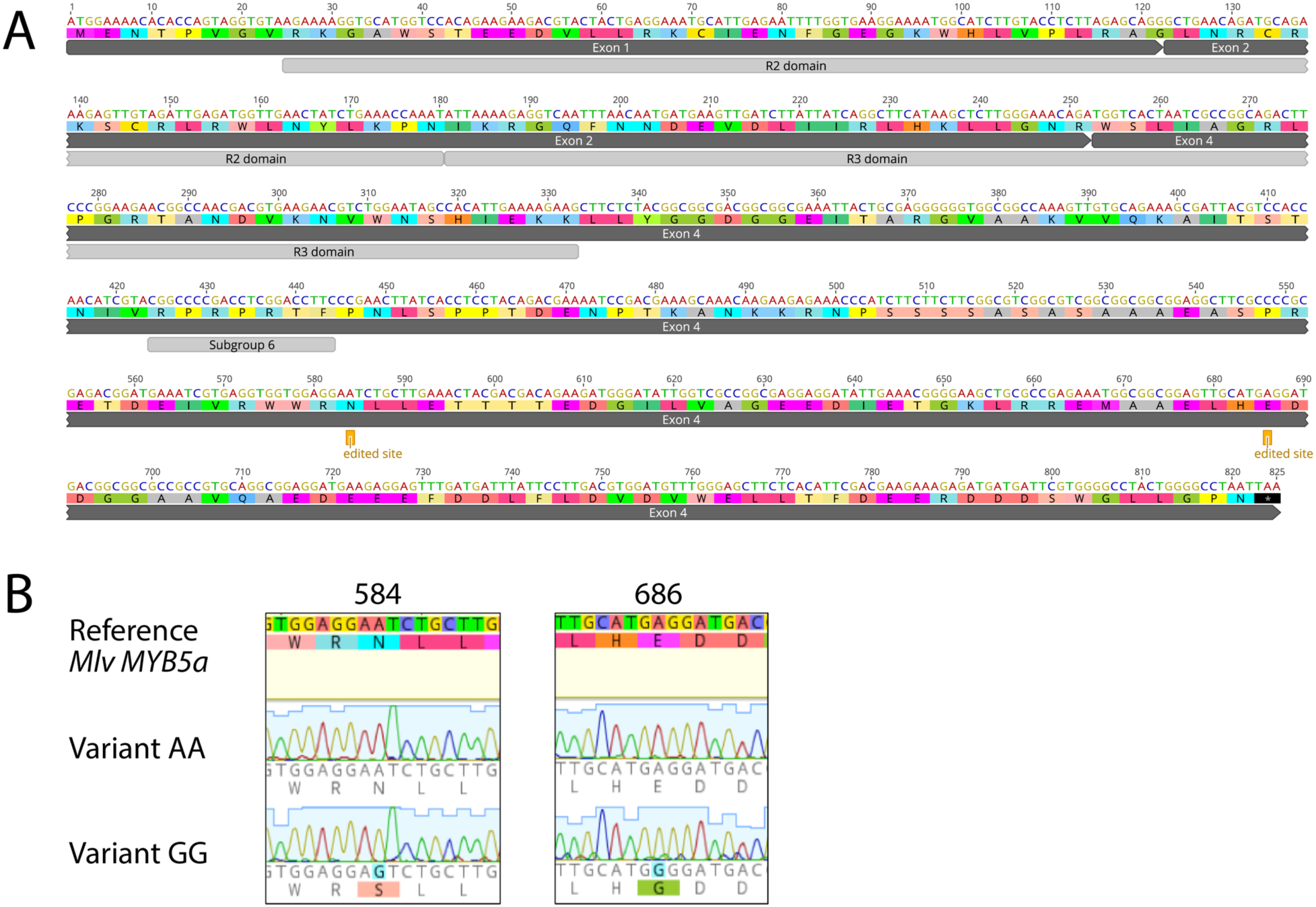
Putative A-to-I editing sites in the exon 1-2-4 splice variant of *M. l. variegatus MYB5a*. A. Dark gray bars show the exon structure from start codon to stop codon. Light gray bars show the DNA- binding R2 and R3 domains common to all members of the R2R3 MYB gene family (Stracke et al. 2001). “Subgroup 6” is a sequence motif that is conserved across all R2R3 MYB genes that encode activators of anthocyanin biosynthesis (Stracke et al. 2001). The two putative A-to-I editing sites are each marked as “edited site”. B. Chromatograms from *MYB5a* Variant AA and Variant GG. The two polymorphic sites are both located in the fourth exon of *MYB5*, 584 and 686 nucleotides downstream of the translation start site. Nucleotide and amino acid differences are highlighted. Sequences were obtained using Sanger sequencing and were visualized using Geneious R10.

To determine whether the “GG” variant represented the genomic sequence of an unknown, related *MYB* gene, we cloned and sequenced a fragment of genomic *MYB5a* from an F1 hybrid of *M. l. luteus* x *M. l. variegatus* using primers Myb5_64F and Myb5_57R (Supplemental Table S1), which encompass the first of the two sites in question; these primers were selected because they reliably amplified *MYB5a* from both *M. l. luteus* and *M. l. variegatus*. We reasoned that, if the “GG” variant were from a paralogous *MYB* gene, then we should be able to recover both variants from the F1 hybrid. The “AA” variant would originate from the *M. l. variegatus MYB5a* and the “GG” variant would originate from the other, unknown gene; we would also expect to recover the *M. l. luteus* allele of *MYB5a*. If the “GG” variant instead represented residual heterozygosity in *M. l. variegatus*, or post-transcriptional editing of the mRNA, then any single F1 hybrid would contain only one of the two *M. l. variegatus* variants, along with the *M. l. luteus* allele of *MYB5a*.

**Table 1.**
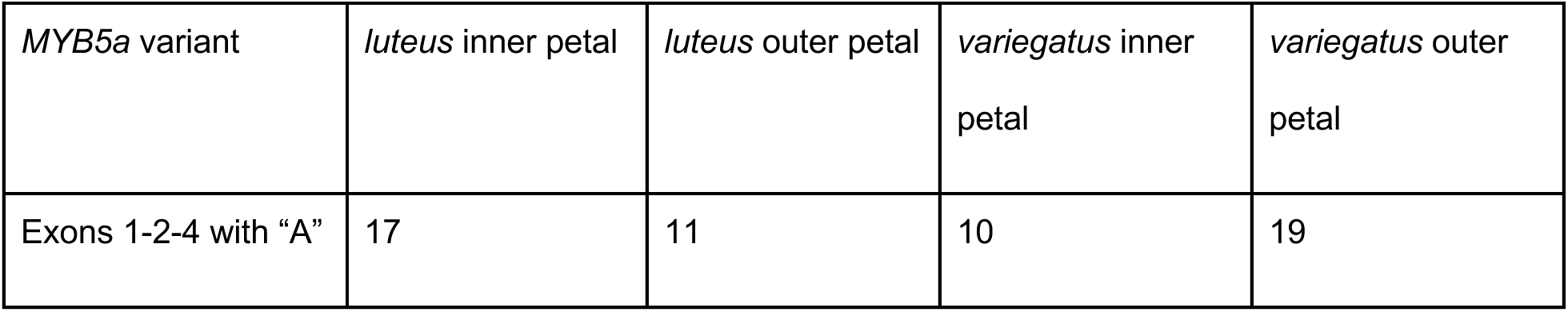
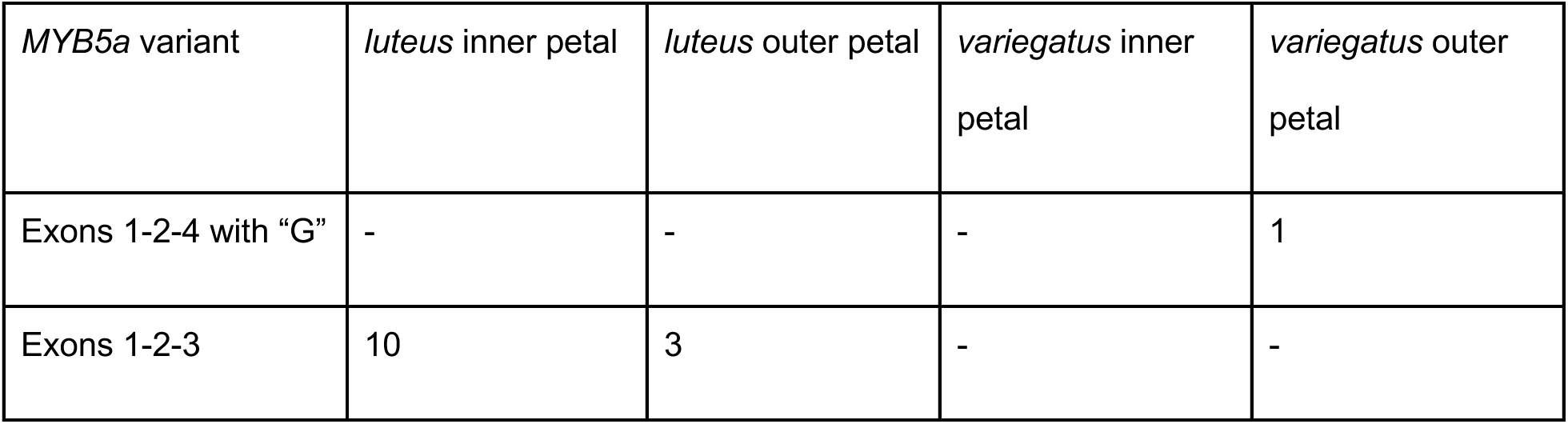
Sequencing MYB5a cDNA from M. l. luteus and M. l. variegatus developing flower bud tissue identifies splice variants in M. l. luteus, and sequence variants in M. l. variegatus. “Inner petal” corresponds to the nectar-guide-spotted throat region of the flower; “outer petal” corresponds to the petal lobes.

| *MYB5a* variant | *luteus* inner petal | *luteus* outer petal | *variegatus* inner petal | *variegatus* outer petal |
| --- | --- | --- | --- | --- |
| Exons 1-2-4 with “A” | 17 | 11 | 10 | 19 |
| Exons 1-2-4 with "G" | - | - | - | 1 |
| Exons 1-2-3 | 10 | 3 | - | - |

To determine whether the “GG” variant represented genomically encoded, residual heterozygosity in our highly inbred *M. l. variegatus*, we similarly cloned and sequenced genomic *MYB5a* from the *M. l. variegatus* inbred line. If the “GG” variant were the result of post- transcriptional modification, it should be absent from all genomic DNA samples (both the F1 hybrid and the *M. l. variegatus*).

Finally, to determine whether the “GG” variant could be repeatably isolated from cDNA, we performed new mRNA extractions, and cloned and sequenced *MYB5a* cDNA, from two types of *M. l. variegatus* and *M. l. luteus* floral tissues: the nectar guide region, which is anthocyanin- pigmented in both taxa, and the petal lobe region, which is anthocyanin-pigmented only in *M. l. variegatus.* Because exons 3 and 4 are partial duplicates of each other, primers Myb5_64F and Myb5_57R (Supplemental Table S1) amplified both splice variants (exon 1-2-3 and 1-2-4) from *M. l. luteus*, though only the exon 1-2-4 variant from *M. l. variegatus*. See Supplemental Figure S6 for an illustration of primer binding sites for both taxa.

### Nucleic acid extraction, PCR, and cloning

Genomic DNA was extracted from young leaves and floral buds using the Zyppy DNeasy Extraction kit (Zymo Research, CA, USA) according to the manufacturers’ protocol. RNA was extracted from buds using the E.Z.N.A. Plant RNA Kit (Omega Bio-Tek, GA, USA) with the DNAse I digestion protocol added to it. cDNA was synthesized using the qScriptTM cDNA Synthesis Kit (Quanta BioSciences, Inc., MD, USA). Quality and concentration of DNA and RNA were quantified using a nanodrop.

Fragments of *MYB5a* spanning one or both adenine/guanine polymorphic sites were PCR amplified using the primers listed in Supplemental Table S1 with G-Biosciences Taq polymerase (St. Louis, MO, USA). Reactions were run with 10µM forward and reverse primers, G-Biosciences 10x buffer, and 2.5µM dNTPs. Annealing temperatures were set to 3℃ below the primer’s lowest melting temperature and the number of PCR cycles ranged from 30-32.

PCR products were purified and cloned into pGEM vectors in *E. coli* as described in Zheng et al. (2021). Colonies were PCR-screened for inserts of the correct size using primers M13F(-20) and M13R(-24). Sanger sequencing was performed by Eton Biosciences (San Diego, CA, USA) and sequences were visualized using Geneious R9 and R10.

### Strategy for testing for functional equivalence of two coding sequences

Once the “AA” allele had been identified as the only genomically encoded *MYB5a* sequence present in the magenta-flowered *M. l. variegatus*, transgenes were constructed to test whether the exon 1-2-4 splice variant was functionally equivalent to the corresponding allele from the yellow-flowered *M. l. luteus* (which lacks petal lobe anthocyanins), as described below. Each transgene, as well as a negative control, was transfected into leaves of *Nicotiana tabacum*, and the area of the spot of anthocyanin pigment produced following each infiltration was quantified.

The heterologous *N. tabacum* system was selected because our pilot studies in *M. l. luteus* and another monkeyflower species, *M. lewisii*, failed to produce visible anthocyanin pigment.

### Bacterial culturing for transgene construction

*Escherichia coli* cultures were grown at 37°C in Luria-Bertani (LB) broth: 10g/L tryptone, 5g/L yeast extract, 10g/L NaCl in demineralized water, sterilized by autoclaving. *Agrobacterium tumefaciens* cultures were grown at 28 °C in LB broth with the NaCl concentration reduced from 10g/L to 5g/L.

Cells containing plasmids with a Kanamycin-resistance gene were grown in media containing 50 µg/mL kanamycin. The *A. tumefaciens* strain, GV3101, used in these studies contains gentamicin- and rifampicin-resistance genes; these cultures were grown in media additionally containing 50 µg/mL gentamicin and 25 µg/mL rifampicin. Liquid cultures were grown at the appropriate temperature with shaking at 200 rpm. To isolate individual colonies, cells were grown on plates containing LB media with appropriate selective antibiotics and 15 g/L agar.

### Construction of Gateway® Entry Vectors

The exon 1-2-4 splice variant of *MYB5a* was amplified from *M. l. variegatus* petal cDNA, using primers cacc10F and Myb5_69R (Supplemental Table S1) and New England Biolabs® Phusion® High-Fidelity DNA Polymerase. Amplicons were transformed into the pEARLEYGATE101 Gateway vector (Earley et al. 2006). From there, the coding sequence without a stop codon was amplified from plasmid DNA containing *M. l. variegatus MYB5a* CDS using the same primers and polymerase.

The pENTR-D/TOPO Cloning Kit was used to produce directionally-cloned Gateway® entry clones carrying *M. l. variegatus MYB5a* CDS. The reaction mixture was transformed into TOP10™ *E. coli* cells, and colonies were screened for the presence of the insert via PCR, using the Myb5_10F, M13R(-24) primer pair. Three colonies that gave a band at ∼1-kb were selected for sequencing. The M13F(-20), M13R(-24) primer pair was used to amplify and sequence the insert. Sanger sequencing was performed by Eton BioScience® (San Diego, CA, USA) and checked for errors against reference sequence in Geneious® version 9.1.8 (https://www.geneious.com).

For *M. l. luteus*, amplification of the exon 1-2-4 splice variant was attempted without success using cDNA extracted from young bud tissue of *M. l. luteus* lines EY1 and EY7. *M. l. luteus* EY7 is known to express *MYB5a* in the anthocyanin-spotted nectar guide region of the flower bud, but expression levels are low (Zheng et al. 2021). Instead, this protein-coding region was synthesized by GENEWIZ (Plainfield, NJ, USA) based on the published genomic sequence of *M. l. luteus* (Edger et al., 2017). The sequence was delivered in a pUC57 vector, but with attL1 and attL2 homology sites added to the 5’- and 3’- ends of the gene, respectively, to facilitate Gateway recombination. Upon receipt, the plasmid was transformed into TOP10™ chemically competent *E. coli* cells, and colonies were screened for the presence of the insert via PCR, using Myb5_12F internal forward primer and M13R(-24) reverse primer (primer table). Sanger sequencing was performed by Eton BioScience, and sequence was checked for errors against the reference sequence in Geneious.

### Construction of Gateway® Plant Expression Vectors

The LR Clonase II kit (ThermoFisher Scientific, Waltham, MA, USA) was used to transfer each insert from an entry vector to destination vector pEARLEYGATE101 (Earley et al. 2006). Entry vectors used in this study were pENTR with *M.l.variegatus MYB5a* CDS and pUC57 with

### M.l.luteus MYB5a CDS

Reactions were transformed into TOP10™ chemically competent *E. coli* and screened for the presence of the insert via PCR and sequencing, using an insert-specific forward primer (Myb5_10F for *M. l. variegatus* and Myb5_12F for *M. l. luteus*) and a reverse primer, att-R2, that binds to the recombination site at the 3’-end of the insert in recombined pEARLEYGATE vectors.

To exclude colonies with unrecombined entry vector, restriction endonuclease digests were performed as an additional diagnostic on plasmid purified from those colonies. Because it cuts both within the destination vector and within the insert, the HinDIII enzyme was used (Promega Corp., Madison, WI). Reactions were incubated for 60 minutes at 37°C, then heat-inactivated for 15 minutes at 65°C. Colonies that gave the expected digest pattern for recombined destination construct as well as the correct insert DNA sequence were chosen to proceed with this project.

### Transformation into Agrobacterium

GV3101 electrocompetent *A. tumefaciens* cells, mixed with 1 µL isolated plasmid DNA (25-350 ng/µL) from each construct, were briefly exposed to a 2.5 kV, 200 ohm, 25 µF pulse using a BioRad® MicroPulser Electroporator (BioRad Laboratories, Hercules, CA, USA). The mixture was then immediately combined with 1 mL room-temperature LB media without antibiotics, incubated at 28°C for 2-3 hours with shaking at 200 rpm., then plated on selective media. Putative transformants were tested for transgene insertion using a PCR screen with primers pEG-35S- attB1_F and att-R2 (Supplemental Table S1).

### pGFP plasmid for negative control

To screen for non-specific effects of transgene infiltration, the *A. tumefaciens-*compatible GFP expression plasmid pGFPGUSPlus was used as a negative control. pGFPGUSPlus was a gift from Claudia Vickers (Addgene plasmid # 64401; http://n2t.net/addgene:64401; RRID:Addgene_64401) (Vickers et al., 2007). Using the protocol in the previous section, the plasmid was transformed into *A. tumefaciens*.

We initially planned to use GFP fluorescence as a read-out of transgene activity, if anthocyanin pigment failed to be produced by either the *luteus* or *variegatus* coding sequence. However, we found that the focal trait of anthocyanin pigmentation was indeed reliably produced. GFP fluorescence, beyond that produced by autofluorescence, was not evident in a pilot study of infiltrated *N. tabacum* leaves (Supplemental Figure S5), and thus was not examined further.

### Transient transformation

A video documenting our transformation methodology, adapted from Ding and Yuan (2016), is available upon request.

*Agrobacterium* colonies were PCR-screened to verify that they contained the desired transgene (pGFPGUSPlus, or *MYB5a* CDS from *M. l. luteus* or *M. l. variegatus*), and were then grown 16- 24 hours in 5 mL LB (*Agrobacterium* recipe) at 28°C with appropriate antibiotics. The small culture was brought up to 50 mL with LB plus antibiotics and grown 12-16 hours at 28°C.

The 50-mL cultures were centrifuged in a Beckman-Coulter Allegra 25R temperature-controlled benchtop centrifuge (Beckman Coulter Inc, Brea, CA, USA) at 4°C and 6,000 RCF for 15 minutes. The pellet was resuspended in a volume of 5% sucrose (w/v) solution equivalent to between one half and one time the original culture’s volume. Resuspensions were adjusted, through dilution with the sucrose solution, to have the same optical density at 600 nm (OD_600_) across all three transgene types within each trial, with an OD_600_ range of 1.6-1.9 across trials. Departing from Ding and Yuan (2016), acetosyringone and Silwet L-77 were not included in the resuspension solution; omitting these reagents was proposed as a possible solution for leaf tissue damage previously observed in infiltrated *Mimulus lewisii* leaves (B. Ding, personal communication).

A B-D 1-mL slip tip disposable SubQ syringe (Becton, Dickinson and Company, Franklin Lakes, NJ, USA) with the needle removed was used to deliver *A. tumefaciens* cells to the leaves of young (1-3 months) *N. tabacum*. Using a gloved finger, the top of the leaf was held firmly while the underside of the leaf was injected with the syringe until the liquid had visibly spread past the site of injection. A volume of 100-200μL resuspended cells was injected per spot. The *M. l. luteus MYB5a* transgene and the *M. l. variegatus MYB5a* transgene were infiltrated in pairs, alternating with each leaf which transgene was injected into the left versus the right side of the leaf. A smaller number of negative controls was performed, with pGFPGUSPlus infiltrations approximately evenly distributed between the two sides of other leaves on the same plants.

### Image acquisition and preparation

Leaves were imaged starting three days after infiltration, since Li et. al. (2009) reported that maximum expression of *Agrobacterium*-delivered transgenes occurs 3 days after infiltration. Because accumulation of visible gene product (anthocyanin) continued for several days after infiltration in some samples, leaves were imaged until a maximum of twelve days after infiltration.

Digital photographs of infected leaves were then taken in a dark room with a Nikon D3500 DSLR camera with 18-55mm lens. The camera was fixed on a stand and the leaf was illuminated by a Sylvania Ceramic Metal Halide bulb, which exceeds 15,000 lumens, as the light source. All exposures produced both RAW and jpeg images, with RAW images used for the analysis and jpeg images used for interoperability with image annotation software. VGG Image Annotator was used to demarcate the regions of interest covering the extent of the infiltrated leaf tissue and the center of each injection site.

### Analysis of images and data

S_green_, which is the strength of the green channel relative to the total of all three color channels, was previously found to correlate highly (R^2^ >= 0.63, p<.001) with anthocyanin concentration across a range of taxa and plant tissues (del Valle et. al. 2018). This is because anthocyanin absorbs light in the green region of the visible spectrum. The index, S_green_, is given by:

S_green_ = N_green_ / (N_green_ + N_blue_ + N_red_)

where N is the intensity value for the green, blue, or red color channel (del Valle et al. 2018).

A custom Python program imported the RAW image files for processing. The Python program used a modified version of the MacDuff color chart detection algorithm (https://github.com/mathandy/python-macduff-colorchecker-detector) to automatically detect the panels of known broad-spectrum reflectance values on a reference color chart. Image pixel values were converted into normalized reflectance values based on a linear fit of the red, green, and blue signal strengths in those panels of known reflectance. The program then averages the relative greenness (S_green_) value over all pixels of the annotated region of interest minus a circular region 20 pixels in radius at the injection site, which typically exhibited tissue damage from the injection syringe. This yielded a single S_green_ value for each sample.

To simultaneously evaluate S_green_ values across all three treatments (the negative controls, the *luteus* transgenes, and the *variegatus* transgenes), a one-way ANOVA was performed in R 4.1.2, followed by a Tukey’s post hoc test. To further compare effects of only the *luteus* and *variegatus* transgenes to each other, taking advantage of the paired design of infiltrations for these two treatments, a two-tailed paired t-test was used.

A number of leaves infiltrated with *MYB5a* did not produce visible quantities of red pigmentation. A *luteus-variegatus* treatment pair was categorized as “no visible pigment” if the S_green_ value for either member of the pair showed equal or lower amounts of red pigmentation than the average of all the negative controls. This categorization corresponded well with a by-eye assessment. Data were analyzed both with and without these apparently-unsuccessful infiltration pairs. Overall rates of success (defined as production of visible pigment) were compared between the *luteus* and *variegatus* transgenes using a χ^2^ contingency test in R 4.1.2.

### Data availability

A detailed transient-transformation methods video is available upon request. Images of all leaves analyzed are linked as Supplemental Figure S9.A, S9.B, and S9.C in the Supplemental Data. Full S_green_ (pigment intensity) data are available as Supplemental Table S2. *MYB5a* sequences from the exon 1-2-4 splice variants of *M. l. luteus* and *M. l. variegatus* have been previously published: https://www.ncbi.nlm.nih.gov/nuccore/MT361119.1 (*M. l. luteus*, also listed as *E. l. lutea*) and https://www.ncbi.nlm.nih.gov/nuccore/2019733960 *(M. l. variegatus*, also listed as *E. l. variegata*). Code for image analysis is available at https://github.com/WhitmanOptiLab/PigmentSpotting.

**Table 2.** The M. l. luteus allele of MYB5a more often produced visible anthocyanin pigmentation in tobacco leaves than did the M. l. variegatus allele, although the difference is not statistically significant. χ^2^ = 3.249, df = 1, p = 0.0715.

| Transgene | Leaves with visible anthocyanin | Leaves without visible anthocyanin |
| --- | --- | --- |
| *luteus MYB5a* | 79 | 19 |
| *variegatus MYB5a* | 67 | 31 |

## Results

### Two sites in MYB5a are occasionally sequenced as guanine rather than adenine

The locations of two putatively edited sites, in exon 4 of *M. l. variegatus MYB5a*, are shown in Figure 2, along with chromatograms from the “AA” variant versus the “GG” variant from our initial, fortuitous discovery of the sequence difference. The novel “GG” variant was found in two different sequencing reactions, originating from two cDNA synthesis reactions from a single mRNA extraction of *M. l. variegatus* young bud outer-petal tissue.

### The “GG” allele of MYB5a from M. l. variegatus is not genomically encoded

Cloning *MYB5a* gDNA from a *variegatus x luteus* F1 hybrid yielded 29 colonies containing a *MYB* sequence. Of these, 13 contained the “AA” variant of the *M. l. variegatus* allele. The remaining 16 contained the *M. l. luteus* allele. The “GG” variant was not discovered in these genomic DNA samples.

Cloning *MYB5a* gDNA from a highly inbred line of *M. l. variegatus* yielded 37 colonies containing a *MYB* sequence. All of these were the “AA” variant of *M. l. variegatus MYB5a*. The “GG” variant was not discovered in these genomic DNA samples. Sample PCR colony screens, from both F1 hybrid gDNA and *M. l. variegatus* gDNA, are shown in Supplemental Figure S7.

After collecting new floral bud tissues, and cloning and sequencing *MYB5a* cDNA from them, we identified one additional colony containing a G at the edited site encompassed by our primers (Table 1). As before, the variant was obtained from the petal lobes (“outer petal”) of *M. l. variegatus*. The other 29 *M. l. variegatus* colonies contained the “AA” variant. In *M. l. variegatus,* only the exon 1-2-4 splice variant was recovered, as expected based on the utilization of primers Myb5_64F and Myb5_57R (Supplemental Figure S6). These same primers were, however, competent to amplify both 1-2-3 and 1-2-4 splice variants from *M. l. luteus*, and they did. We found 28 colonies containing the exon 1-2-4 splice variant of *M. l. luteus MYB5a*, and 13 containing the exon 1-2-3 splice variant, with both splice variants appearing in both inner and outer petal tissue (Table 1).

Recovering *M. l. luteus MYB5a* sequence from outer petal tissue was unexpected, and may reflect imprecise separation of the two tissue types during floral bud dissection. The finding is consistent with RT-PCR of *MYB5a* from our four cDNA samples, which indicated the presence of some *MYB5a* transcript in the *M. l. luteus* outer petal sample (Supplemental Figure S8).

### Infiltration of transgenes into N. tabacum leaves

Of the 124 leaves that received paired infiltrations of the 1-2-4 splice variant of *MYB5a* (*M. l. variegatus* on one side of the central vein, and *M. l. luteus* on the other), 26 were eliminated due to tearing or inadvertent marking over the pigmented area. The remaining 98 were scored for the presence of visible anthocyanin pigmentation, and also quantitatively analyzed for pigment abundance. Of the 25 leaves infiltrated on each side of the midvein with pGFPGusPlus as a negative control, five were eliminated due to tearing or marking errors and the remaining 20 leaves (40 infiltrations) were quantitatively analyzed for anthocyanin pigment abundance (Supplemental Table S2, Supplemental Figure S9).

### MYB5a from both M. l. luteus and M. l. variegatus drives strong anthocyanin production

Both *MYB5a* transgenes resulted in significantly redder leaf tissue, as indicated by lower S_green_ values, than did the negative control (Figure 3 and 4; F(2, 233) = 12.43; p<0.0001). Surprisingly, the allele of *MYB5a* from the yellow-flowered *M. l. luteus* drove significantly greater anthocyanin production than the corresponding allele from the magenta-flowered *M. l. variegatus* (Tukey’s post hoc test: p=0.0102). After excluding the negative control to compare only the *luteus* and *variegatus* transgenes via a paired t-test, the difference was even more strongly significant (t = - 6.3656, df = 97, p = 6.461*10^-9^; mean difference = 6.456*10^-3^ S_green_ units).

**Figure 3.**
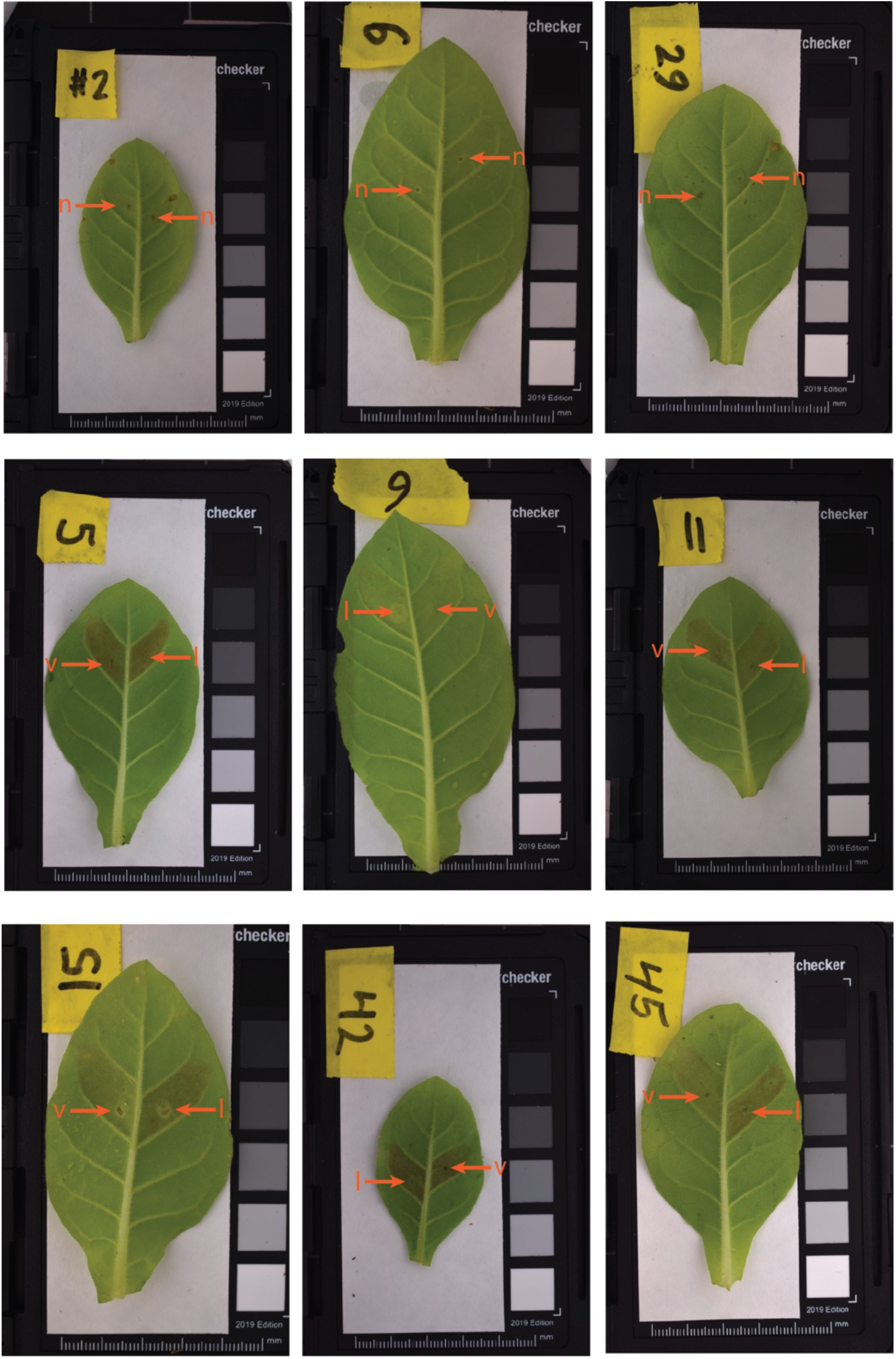
*MYB5a* transgenes yielded a range of anthocyanin biosynthesis levels in *N. tabacum* leaves. In each image, the injection site is indicated with an arrow, labeled as n (negative control), l (*luteus* allele of *MYB5a*), or v (*variegatus* allele of *MYB5a*). Leaf photos for this figure were uniformly brightened by 30% in Powerpoint to make anthocyanin pigmentation and injection sites easier to see, although analyses were performed on un-brightened images.

**Figure 4.**
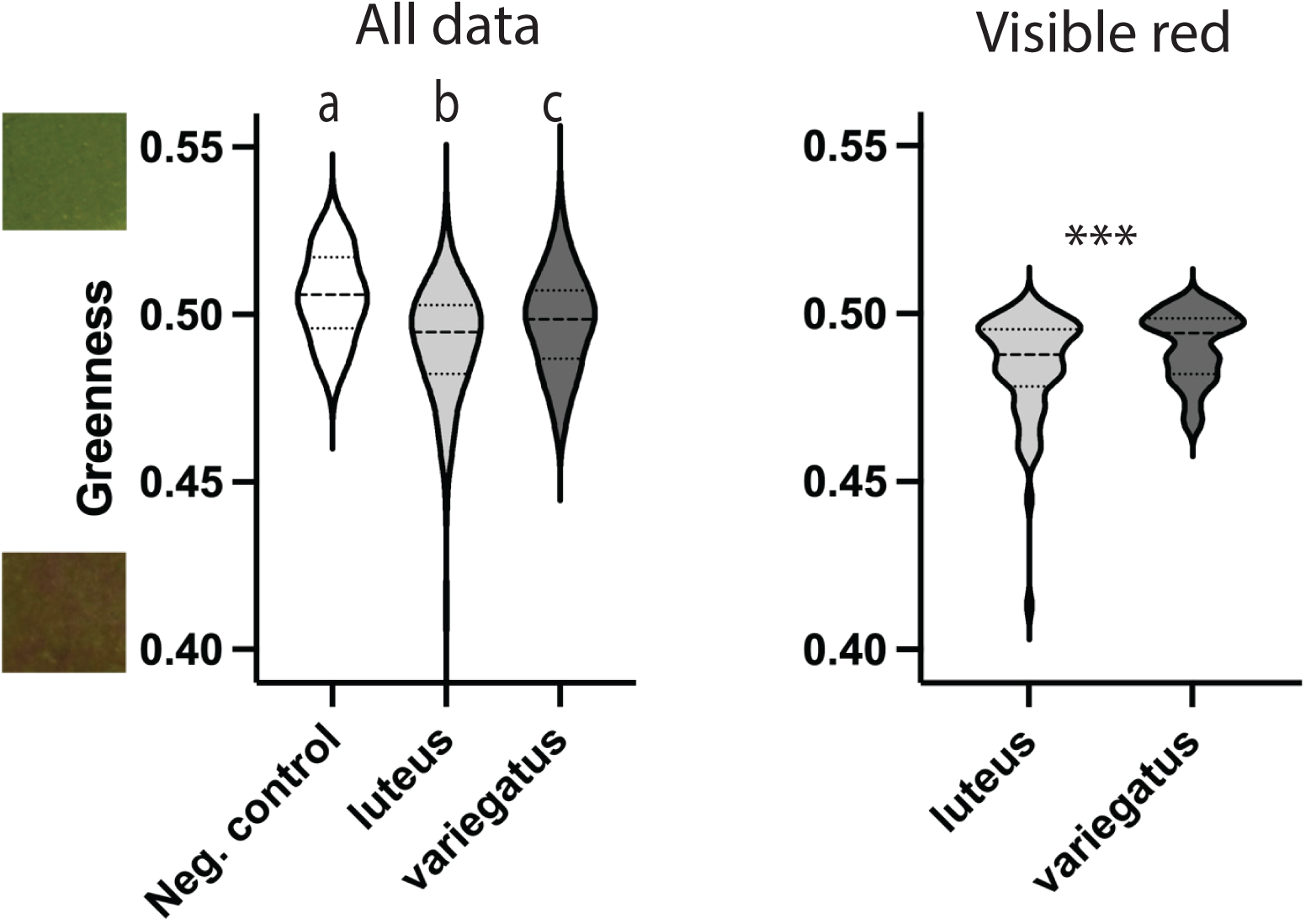
*MYB5a* alleles from both the yellow-petaled *M. l. luteus* and the magenta-petaled *M. l. variegatus* drive production of red anthocyanin pigmentation in *N. tabacum* leaf tissue. Left, data from all samples; *N* = 40 (negative control), 98 (*luteus* coding sequence), 98 (*variegatus* coding sequence). Right, data only from matched *luteus*-*variegatus* pairs for which both members of the pair produced visible pigmentation; *N* = 66 pairs. Each graph is a violin plot with the median shown as a dashed line and the first and third quartiles as dotted lines. On the Y-axis, low values correspond to redder leaf tissue and high values correspond to greener tissue. Color swatches on the Y-axis were taken from injection sites with Sgreen values of 0.53 and 0.41 respectively. Swatches were taken from leaf photos that had been uniformly brightened by 30% in Powerpoint, to maintain consistency with the previous figure, although analyses were performed on un-brightened images. Letters a, b, and c are significance groupings at p<0.05 based on a one-way Anova with Tukey’s post hoc test; *** indicates p<0.001 based on a two-tailed paired t-test.

We wondered whether this result might have been influenced by different rates of infiltration success, rather than by different quantities of pigment produced in successful infiltration events. Overall, 79/98 *luteus* infiltrations, but only 67/98 *variegatus* infiltrations, resulted in the accumulation of visible anthocyanin pigmentation in a tobacco leaf (Table 2). The difference in success rate between the two alleles approached but did not reach statistical significance (χ^2^ = 3.249, df = 1, p = 0.0715).

To exclude possible effects of varying infiltration success, we analyzed pigment production only from the pairs of infiltrations for which both members of the pair produced visible amounts of leaf anthocyanin pigment (Figure 4 right panel). In these pairs, a slightly smaller but still highly significant difference was found, again with the *luteus* coding sequences corresponding to lower S_green_ values and thus more-intense red pigmentation (t = -4.4855, df = 65, p = 3.032*10^-5^; mean difference = 6.170*10^-3^ S_green_ units).

## Discussion

When closely related taxa show phenotypic divergence, is the molecular mechanism a mutational difference in the protein-coding region of a gene, or in the noncoding, *cis*-regulatory region? We used transient transgenic assays to investigate this question for the MYB5a anthocyanin-activating transcription factor from two varieties of monkeyflower: the yellow-flowered *Mimulus luteus* var. *luteus*, which lacks anthocyanin pigment in its petal lobes, and the magenta-flowered *M. l. variegatus*, which recently evolved petal lobe anthocyanin via an unknown change within the *MYB5a* gene. Using quantitative image-analysis based methods, we report that there is a significant functional difference between the two protein-coding regions when expressed in *N. tabacum* leaves, but in the opposite direction of what would be expected if coding sequence evolution were responsible for the increased pigmentation observed in *M. l. variegatus*. Together with a previous finding that *MYB5a* is more strongly expressed in *M. l. variegatus* petal lobes than in its conspecific (Zheng et al. 2021), this result strongly indicates that *cis*-regulatory evolution is responsible for the recent expansion of pigmentation in *M. l. variegatus*.

### Improved tools for rapid transgenic assays in Mimulus

The success of the two *Mimulus* transgenes at activating anthocyanin production in *N. tabacum* is encouraging for future functional studies in *Mimulus*, though not unprecedented. One factor that can limit the ability of a MYB to function in a heterologous system is the availability of a functional bHLH co-factor. In some cases, co-expression of the focal MYB gene’s native bHLH partner has been necessary for successful anthocyanin activation (Espley et al., 2009; Lin-Wang et al., 2010), but in other cases, anthocyanins have been induced in *N. tabacum* without also expressing bHLH from the same system (Fraser et. al., 2013). In one such study, Montefiori et. al. (2015) expressed *AcMyb110* from Kiwifruit (*Actinidia* sp., order Ericales) in *N. tabacum* leaves and successfully stimulated anthocyanin production. They identified two endogenous bHLH transcription factors in *N. tabacum*, NtJAF13 and NtAN1, that associated with AcMYB110 to stimulate expression of anthocyanin biosynthetic pathway genes, and that may also have interacted with the *Mimulus* MYB5a protein in our experiment. While stable transgenics are more useful for many applications, the ease and rapidity of transient transformation approaches makes it a valuable tool when studying a readily quantifiable phenotype such as pigment production.

One caveat to the quantification of anthocyanin activation by the *luteus* and *variegatus* transgenes is that high levels of transgene expression could potentially alter patterns of functionality that only appear at lower concentrations (Koes et. al., 2005). Developing a more closely related species as a platform for functional tests would also be beneficial. In our hands, preliminary tests with both *M. luteus* and congener *M. lewisii* resulted in high levels of tissue death and damage, but the latter appears to have promise as a host for transient transgenic assays (Ding and Yuan 2016).

Using *N. tabacum* for direct side-by-side comparison of *Agrobacterium-*delivered transgenes was first reported by Van der Hoorn et. al. (2000). This strategy takes advantage of leaf symmetry and the clearly delineated leaf sectors in *N. tabacum* to compare two genes side-by-side in an identical biological background. Coupled with a nondestructive way to quantify the resulting phenotype, we believe this remains an underutilized strategy for functional comparisons between genes. Finally, we note that the significant difference in pigment production between the *luteus* and *variegatus* transgenes would almost certainly not have been detected using a qualitative assessment of pigmentation, highlighting the benefits of even relatively simple computer-vision based programs to quantify color traits in plants.

### Possible A-to-I editing of mRNA

In the process of building *MYB5a* overexpression transgenes, we discovered what appears to be the first documented case, to our knowledge, of post-transcriptional editing in an anthocyanin-related gene. Two sites within the *M. l. variegatus* allele are encoded as adenine in the genome, yet occasionally produce mRNA sequences that read as a guanine in Sanger sequencing.

Inosine is a guanine analog, most often created in cells by the deamination of adenine (Srinivasan et al., 2021), that is reported as guanine in Sanger sequencing (Cattenoz et al., 2013). Adenine-to-inosine (A-to-I) editing was first discovered in *Xenopus laevis* mRNA by Bass and Weintraub (1988), and is abundant in metazoans (Cattenoz et al., 2013), with one study predicting over 36,000 A-to-I editing sites in the human genome (Li et al., 2009).

Although A-to-I editing of mRNA transcripts does not yet appear to have been directly investigated in plants, we hypothesize that the two new bases are in fact inosine, given that inosine is reported as guanine by Sanger sequencing (Cattenoz et al., 2013). Nuclear A-to-I post-transcriptional editing has been reported in plant tRNA (Delannoy et al., 2009; Karcher and Bock 2009; Zhou et al., 2014), and the deaminase enzymes required for A-to-I editing have been putatively discovered in *Arabidopsis thaliana* (Zhou et al., 2014). Though mRNA A-to-I editing has not been described in plants, it is widespread across the domains of life, including fungi (reviewed in Teichert 2018), animals (reviewed in Knoop 2011), and bacteria (first reported by Bar-Yaacov et al., 2017).

In contrast, reports of A-to-G editing in plants appear to be based solely on sequencing-based approaches that would report inosines incorrectly as guanines (Pan et al., 2022), with A-to-I editing apparently first proposed by Meng et al. (2010) on the basis of A-to-”G” mRNA editing discovered by sequencing plant transcriptomes. True A-to-G editing does not appear to be a verified biological phenomenon in any taxon. We therefore consider A-to-G editing to be less likely than A-to-I editing in our study.

A variety of methods exist for confirming A-to-I editing, including chromatographic approaches (Wolf et al., 2002; Chan et al., 2010) and “inosine chemical erasing” (ICE)-Seq (Sakurai et al., 2010). However, A-to-I editing is commonly detected and quantified by the simple method used here, in which reverse transcription and sequencing of mRNA reveals unexpected “guanines” in some proportion of transcripts (e.g. Gu et al., 2012).

When inosine is present in tRNA, it can pair promiscuously with A, C, or U. In mRNA transcripts, in contrast, it is translated as though it were guanine (Srinivasan et al. 2021). Regardless of whether the edited bases result in guanine or inosine, then, they are likely to be interpreted by the translational machinery of the cell as guanine. In the *M. l. variegatus* allele of *MYB5a*, both edited sites would result in an amino acid change: from asparagine to serine at nucleotide position 584, and from glutamic acid to glycine at position 686.

How conservative are these changes? One metric is Grantham’s Distance, based on composition, polarity and molecular volume (Grantham 1974). Using this metric, amino acid pairs have similarity scores ranging from 5 for the highly similar leucine-isoleucine pair to a maximum of 215 for cysteine-tryptophan. The first putative editing site reported here (asparagine-serine, both of which have polar side chains) has a modest Grantham’s Distance of 46. The putative glutamic acid to glycine substitution - replacing a negatively charged side chain with a single hydrogen - has a larger Grantham’s Distance of 98. The implications for MYB5a protein folding and function are, however, unknown. Overall, the mechanisms for A-to-I (or A-to-G) editing in plant mRNAs, and their functional impacts, comprise a barely-explored area within plant molecular biology, which seems likely to yield new discoveries upon further investigation.

Nanopore native RNA sequencing methods, recently used by Nguyen et. al. (2022) to globally identify inosine in human, mouse, and *Xenopus* transcriptomes, might be applied fruitfully to plant transcriptomes with the same aim.

## Conclusions

*Mimulus luteus* var. *luteus* and *M. l. variegatus* differ strikingly in floral phenotype, thanks to a derived loss of yellow carotenoid pigment and gain of magenta cyanidin pigment in the latter. The expansion of cyanidin to the petal lobes of *M. l. variegatus* has previously been tracked to the *MYB5a* transcription factor gene, for which the patterns of petal expression correlate well with the presence versus absence of cyanidin pigment. Here, we use transient transgenics in *N. tabacum*, coupled with quantitative digital image analysis, to show that the protein-coding region of *MYB5a* from the high-anthocyanin *M. l. variegatus* does not show any increased anthocyanin- activation ability (in fact, the reverse) compared to the orthologous sequence from the low- anthocyanin *M. l. luteus*. This finding adds further support to the hypothesis that evolution in *cis* to *MYB5a* is the molecular mechanism for the gain of this novel anthocyanin trait in *M. l. variegatus*.

We additionally report the discovery of what appears to be post-transcriptional mRNA editing. The edits are reported as A-to-G by Sanger sequencing, but we argue that A-to-I editing is more likely based on what is known about RNA editing in plants and other organisms. Overall, our work highlights the utility of floral diversification for identifying the molecular mechanisms of evolution, as well as the scope for continued new discoveries in the realm of plant molecular genetics.

## Acknowledgements and Funding

The authors thank N. Forsthoefel, B. Ding, and Y.-W. Yuan for assistance with and advice on the transgenic techniques. We thank D. Vernon, B. Moss, J. Puzey, and their students for discussion of some of the results presented here. AMC was supported by NSF-DEB-1655311, NSF-DEB-1754075, and NSF-IOS-2031272. AEP was additionally supported by a Whitman College Abshire Award for undergraduate research.

## Supplemental tables

**Supplemental Table S1.**
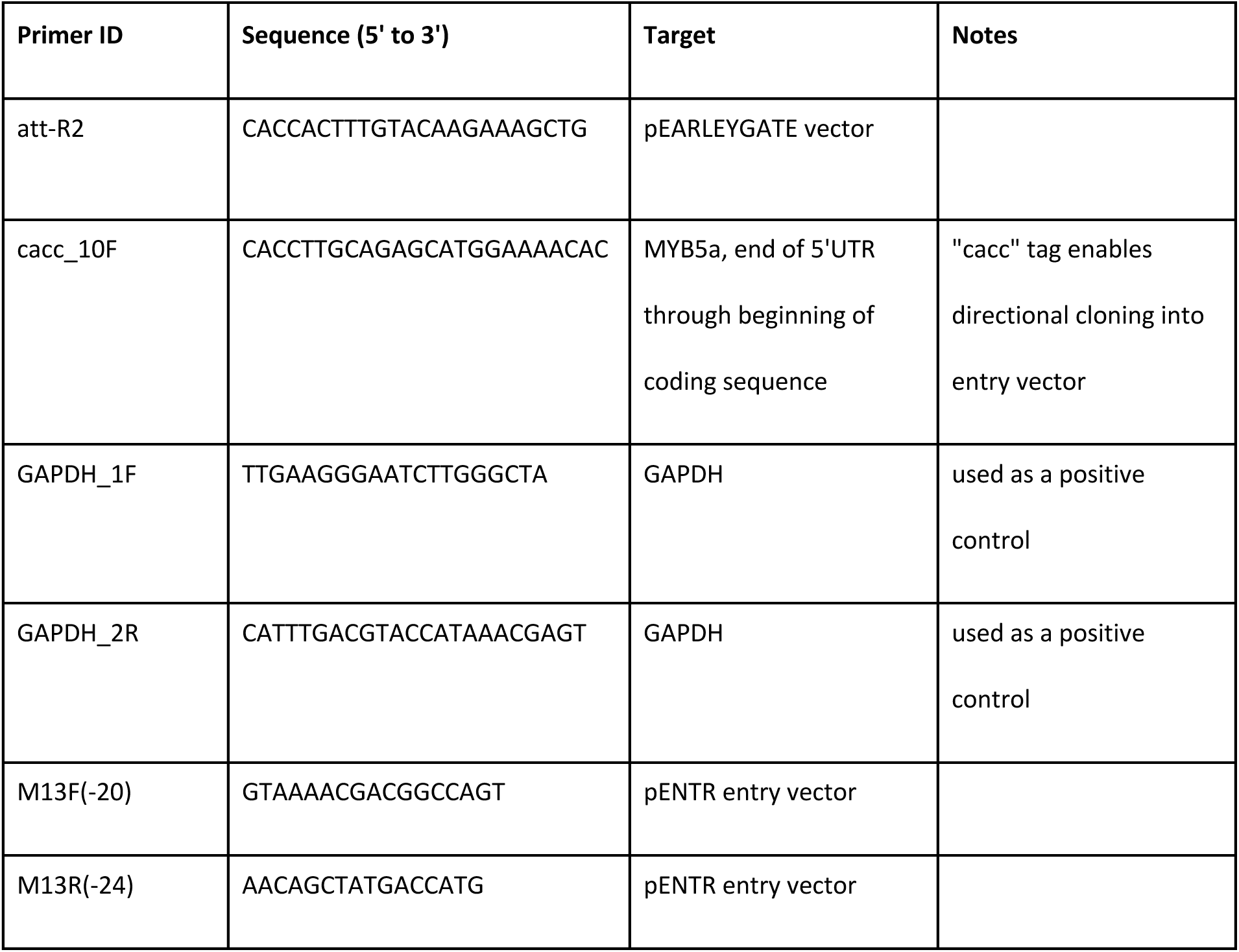
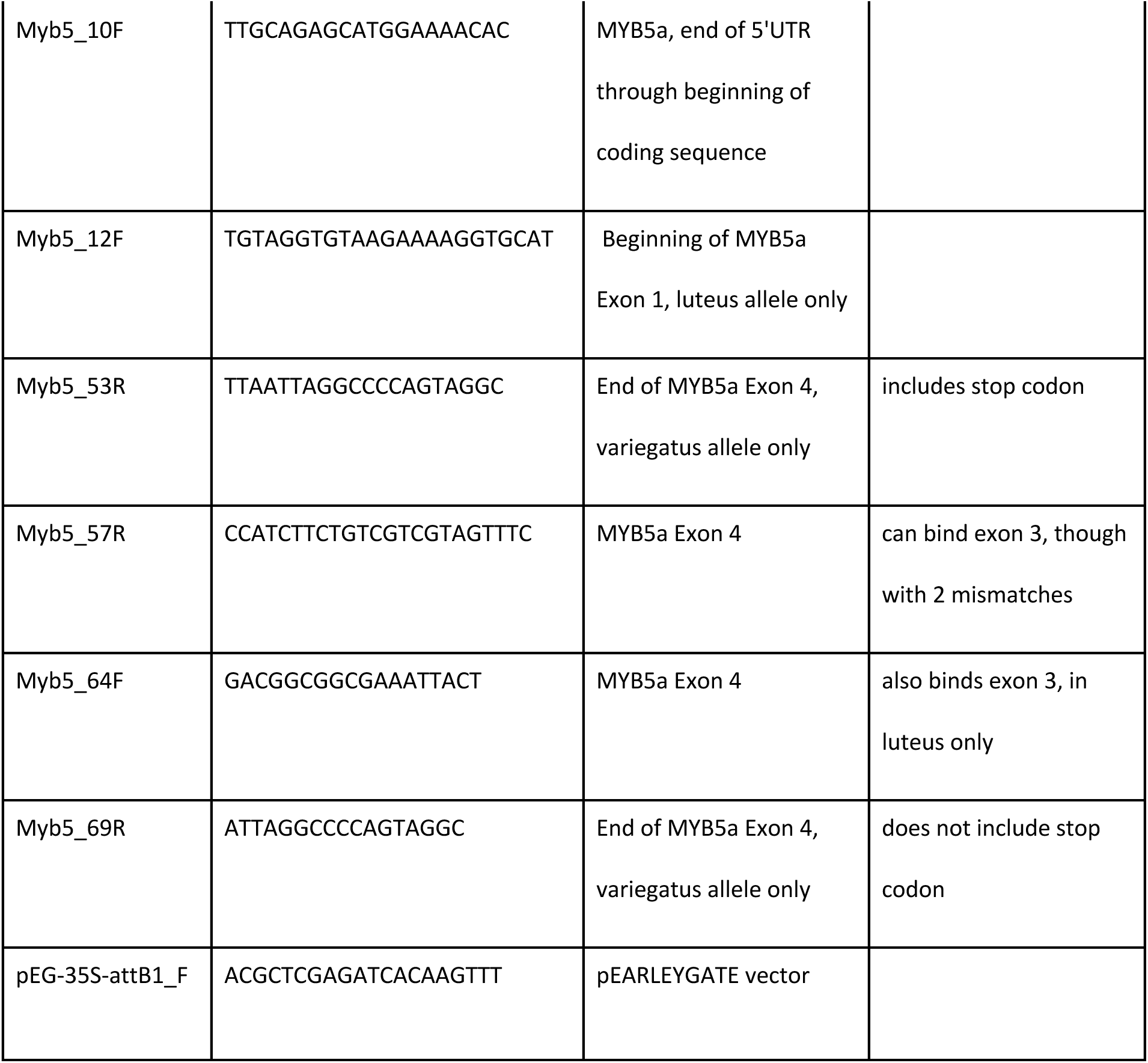
Primers used. F and R in primer names indicate forward and reverse primer directions with respect to the direction of transcription. *MYB5a* primers bind to both *M. l. variegatus* and *M. l. luteus* alleles unless otherwise noted.

**Supplemental Table S2.**
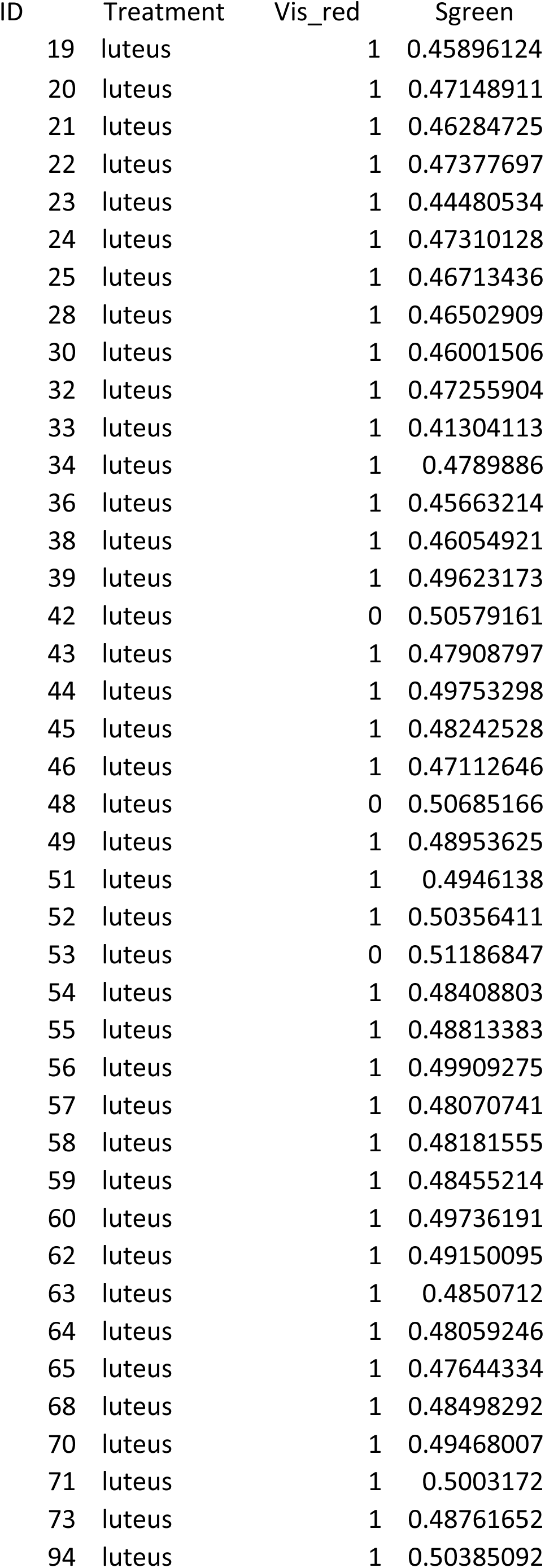
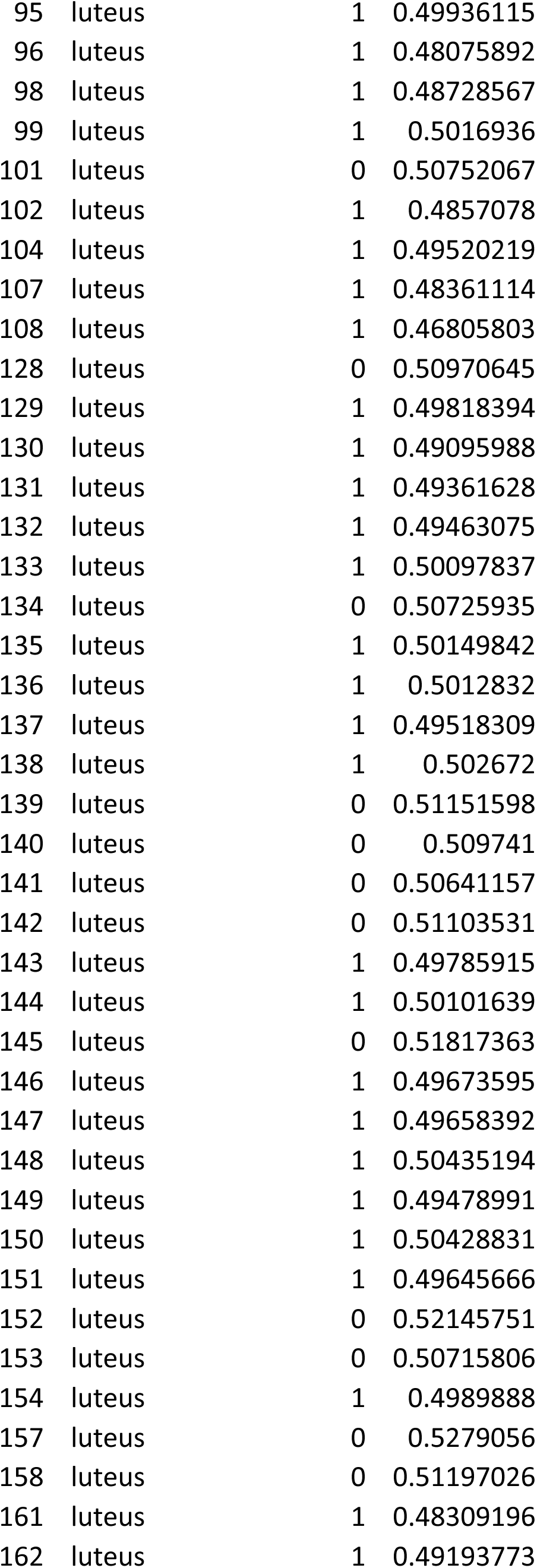
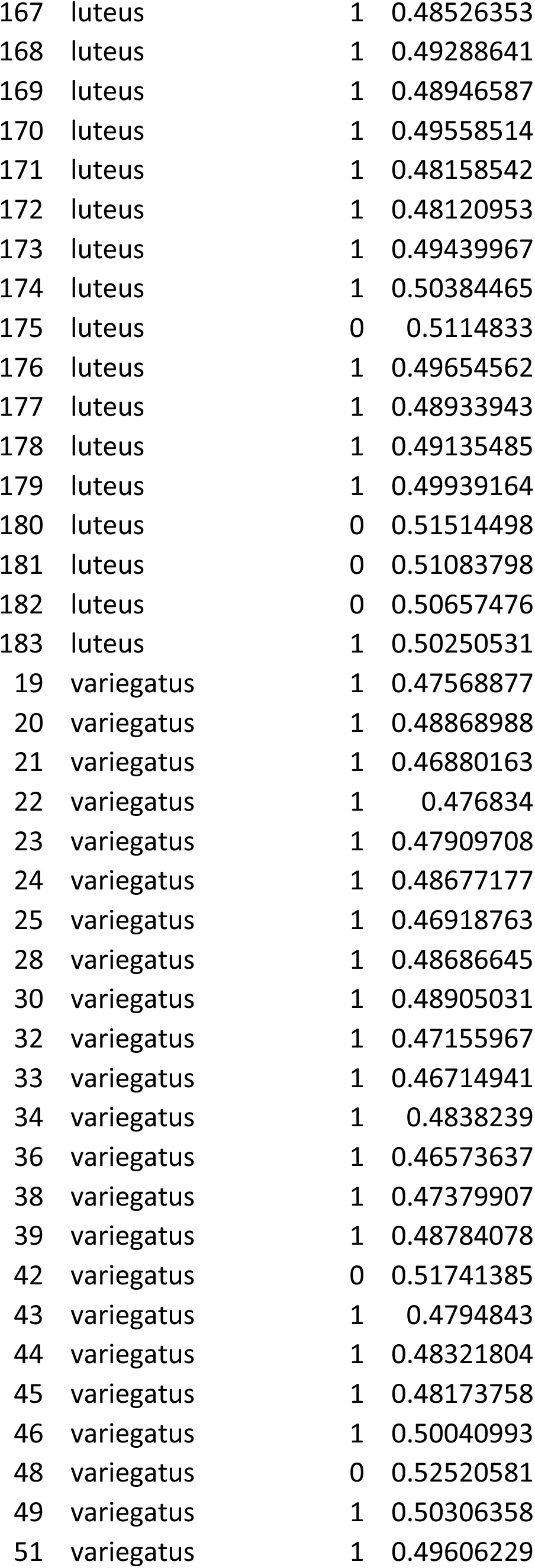
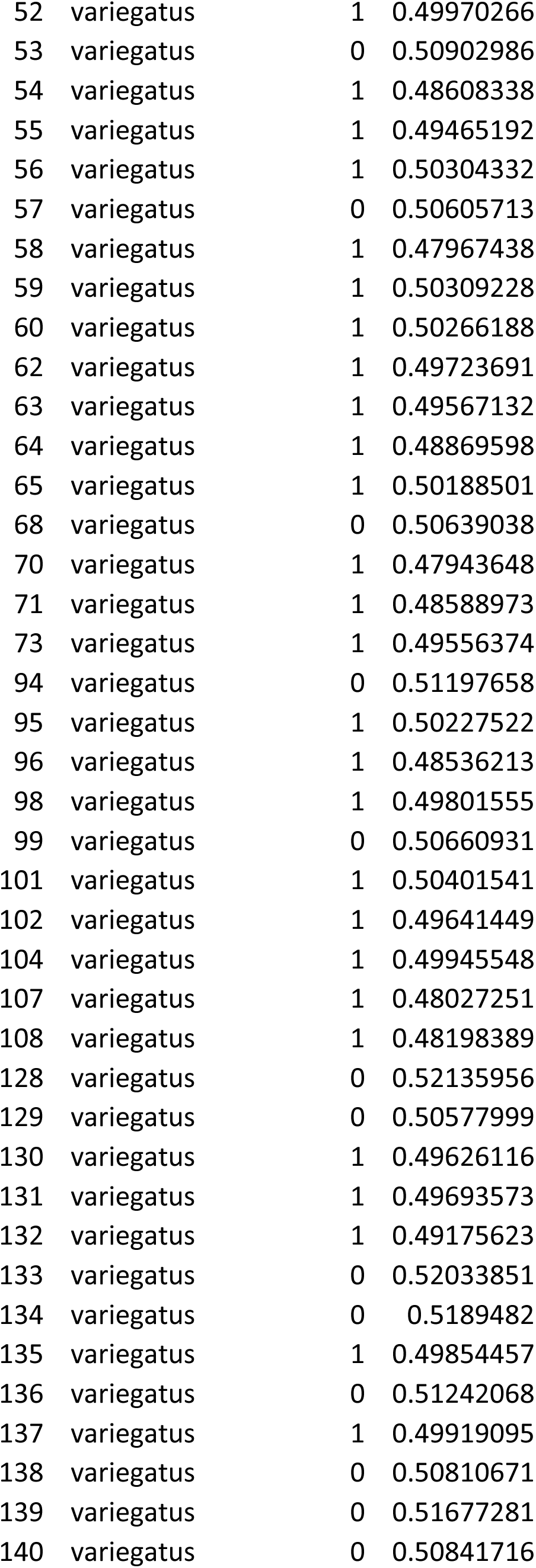
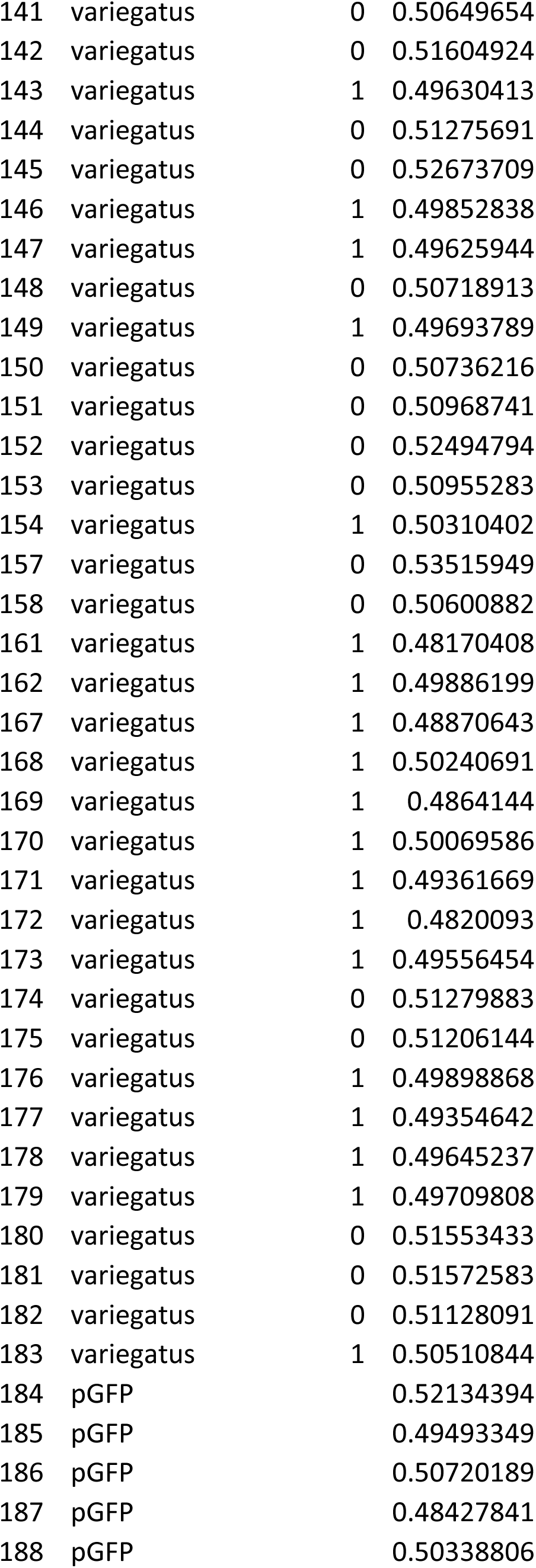
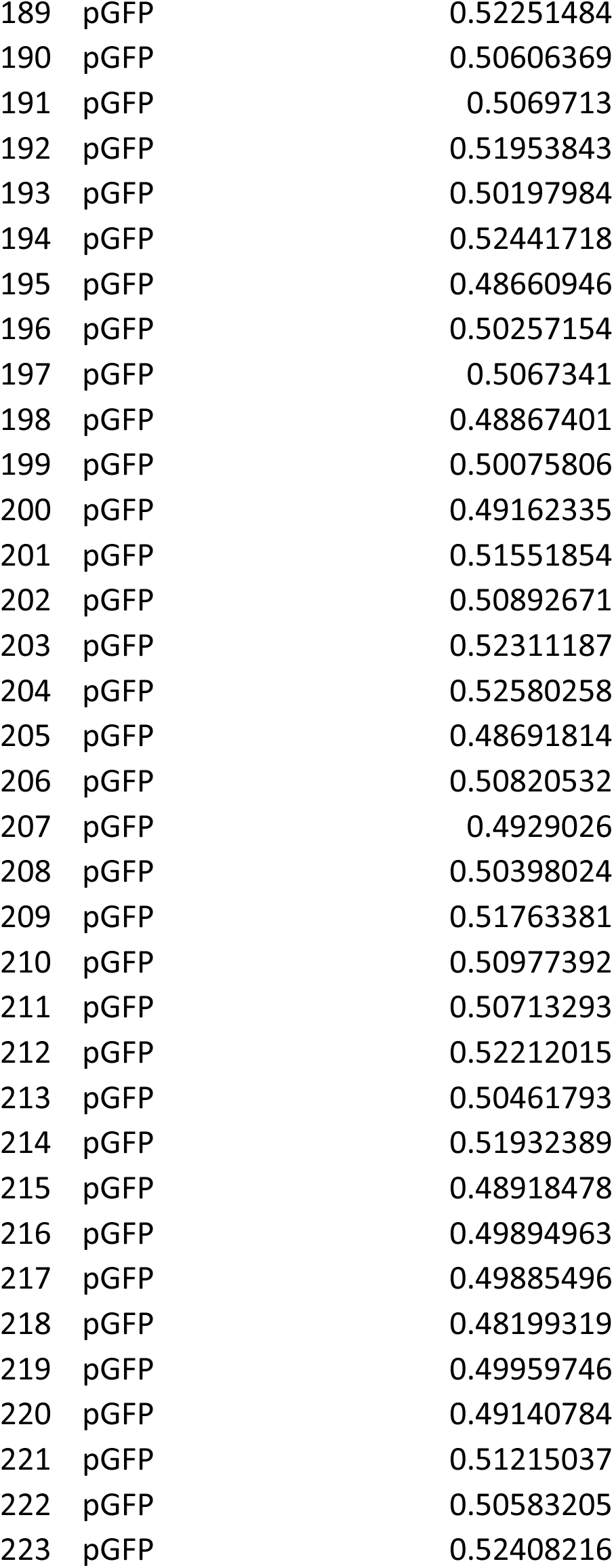
S_green_ pigment intensity data for all analyzed samples. Damaged leaves were excluded from the dataset prior to analysis. A lower S_green_ value corresponds to a greater intensity of red pigmentation. Each pair of *luteus* and *variegatus* transgenes that were infiltrated into the same leaf also share an ID number. In the “Vis_red” column, 1 indicates pigment intensity higher than the average for the negative controls.

## Supplemental figures

**Supplemental Figure S1.**
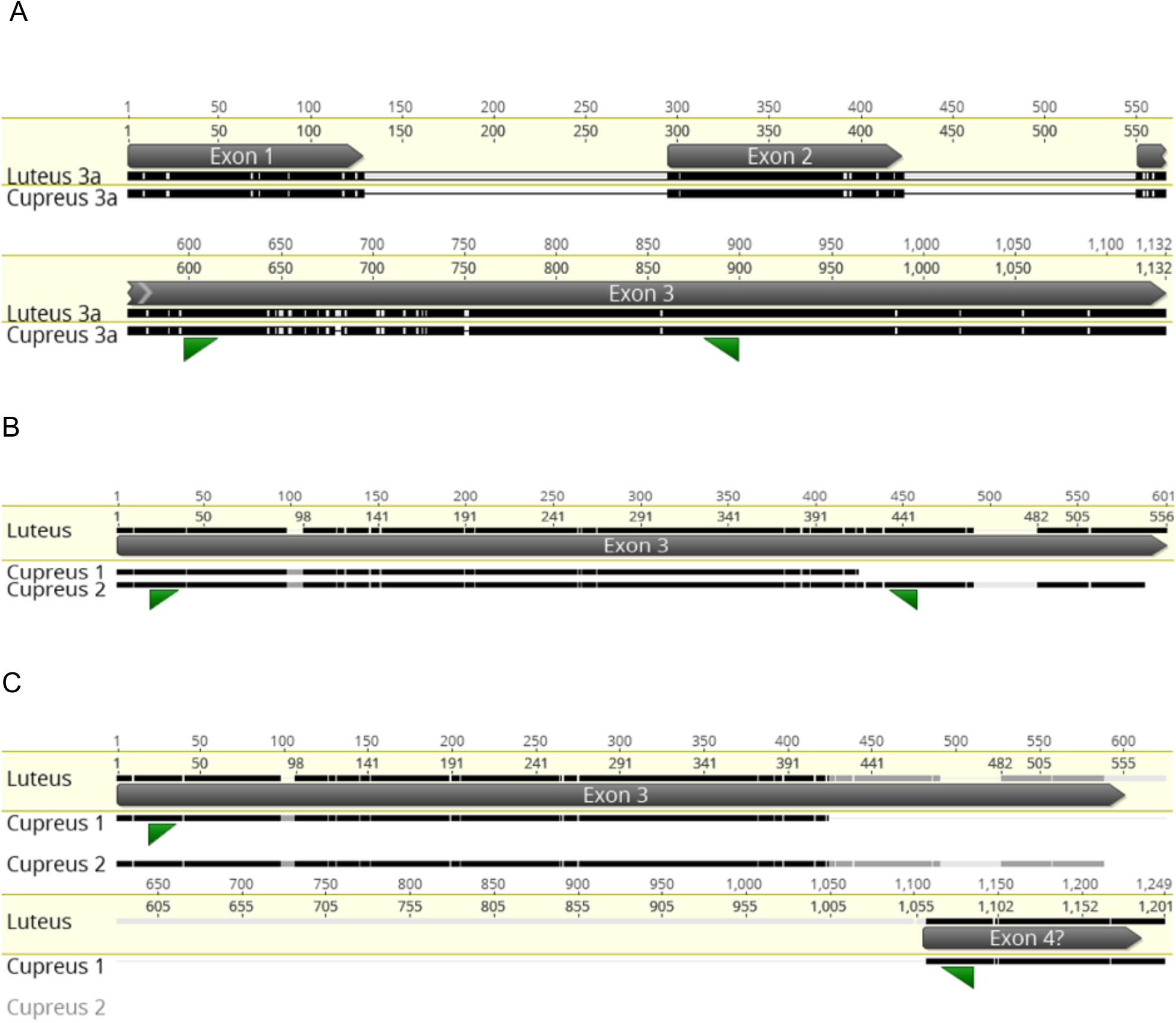
M*i*mulus *cupreus MYB3a* and *MYB2* transcripts aligned to *M. luteus* genomic sequence. A. *MYB3a*. Primers Myb2/3_1F and Myb3a_2R are shown in green. B. *MYB2*. Two *M. cupreus* transcripts were recovered from transcriptome sequence; the region in which they diverge is labeled as “Cupreus 1” and “Cupreus 2,” possibly corresponding to alternative splice variants. Primers Myb 2_1F and Myb 2b_5R shown in green. These primers are expected to amplify only transcript 2. C. The same *MYB2* transcripts with primers Myb 2_1F and Myb 2b_7R shown in green.

**Supplemental Figure S2.**
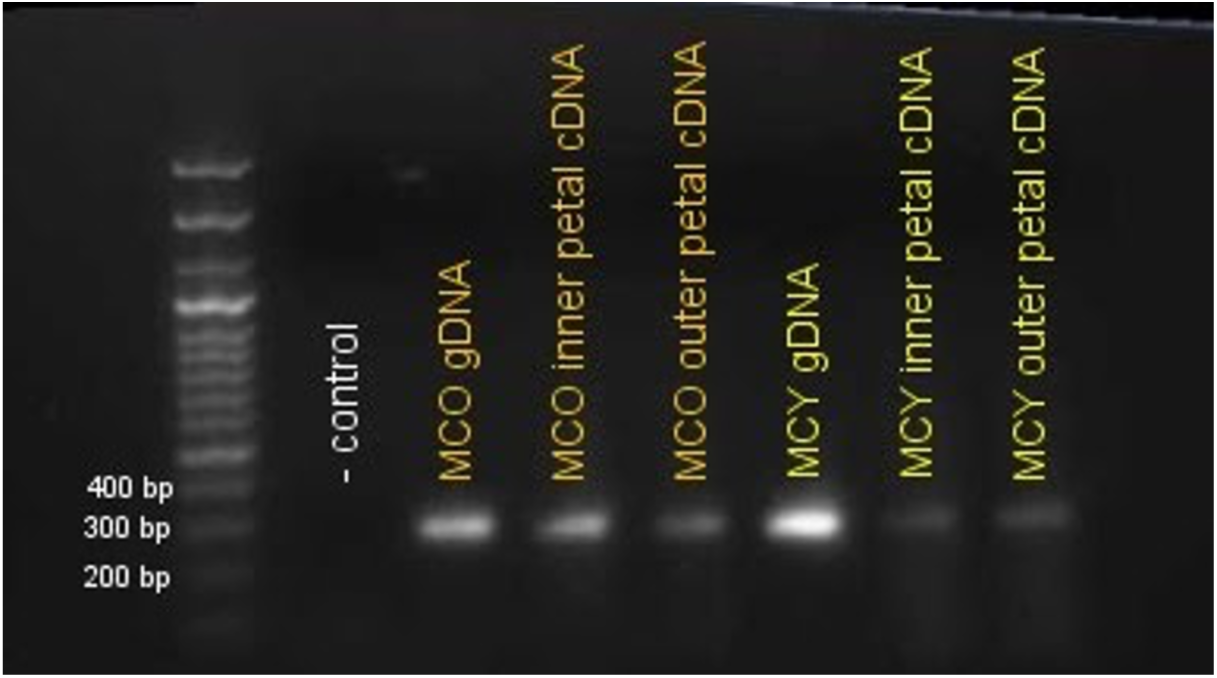
Primers Myb 2/3_1F and Myb 3a_2R were used to amplify a portion of *Myb3a* exon 3, encoding a transcription factor gene that is one of two candidates for the gain of petal anthocyanin pigmentation in the orange-flowered morph of *M. cupreus* (Cooley et al., 2011). The recent loss of pigmentation in the rare yellow morph of *M. cupreus* also maps to the same region. A product of the expected length (300 bp) was amplified out of gDNA for both orange and yellow-flowered *M. cupreus.* The product was also amplified out of cDNA for both inner and outer petal tissue from both morphs, indicating that Myb 3a **is** expressed in all of these tissues.

**Supplemental Figure S3.**
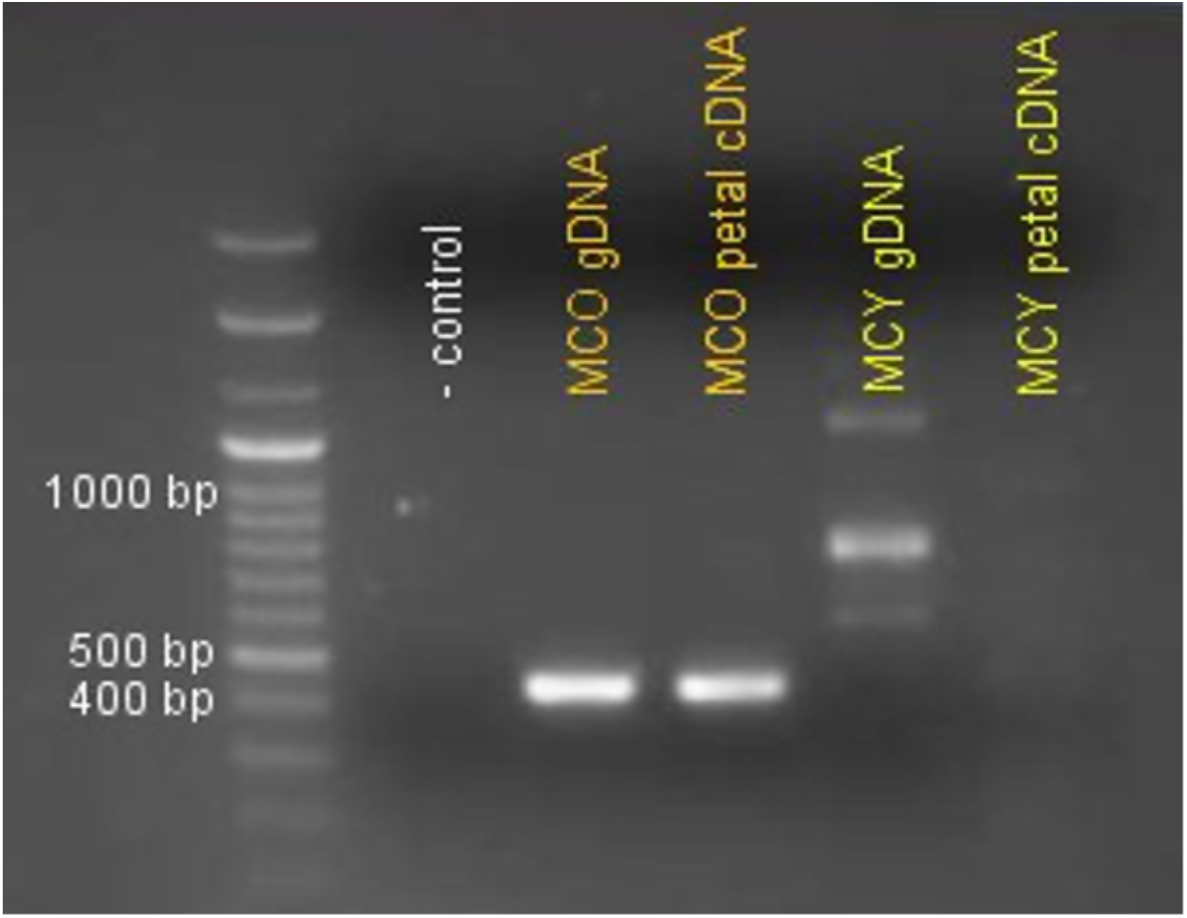
Primers Myb 2_1F and Myb 2b_5R were used to amplify a portion of *Myb2* transcript 2, exon 3, encoding a transcription factor gene that is the second of two candidates for the derived gain of petal anthocyanin pigmentation in the orange-flowered morph of *M. cupreus*, and the even more recent loss of petal anthocyanin in the rare yellow morph. A product of expected length (around 450 bp) was amplified out of orange-flowered *M. cupreus* gDNA. The product was also amplified out of cDNA from both inner and outer petal from orange-flowered *M. cupreus,* indicating that it is expressed in the orange morph. However, no product of the expected length was amplified out of yellow-flowered *M. cupreus* gDNA. Several longer products were amplified less brightly, and are likely due to nonspecific annealing. No product was seen in the yellow-flowered petal cDNA either.

**Supplemental Figure S4.**
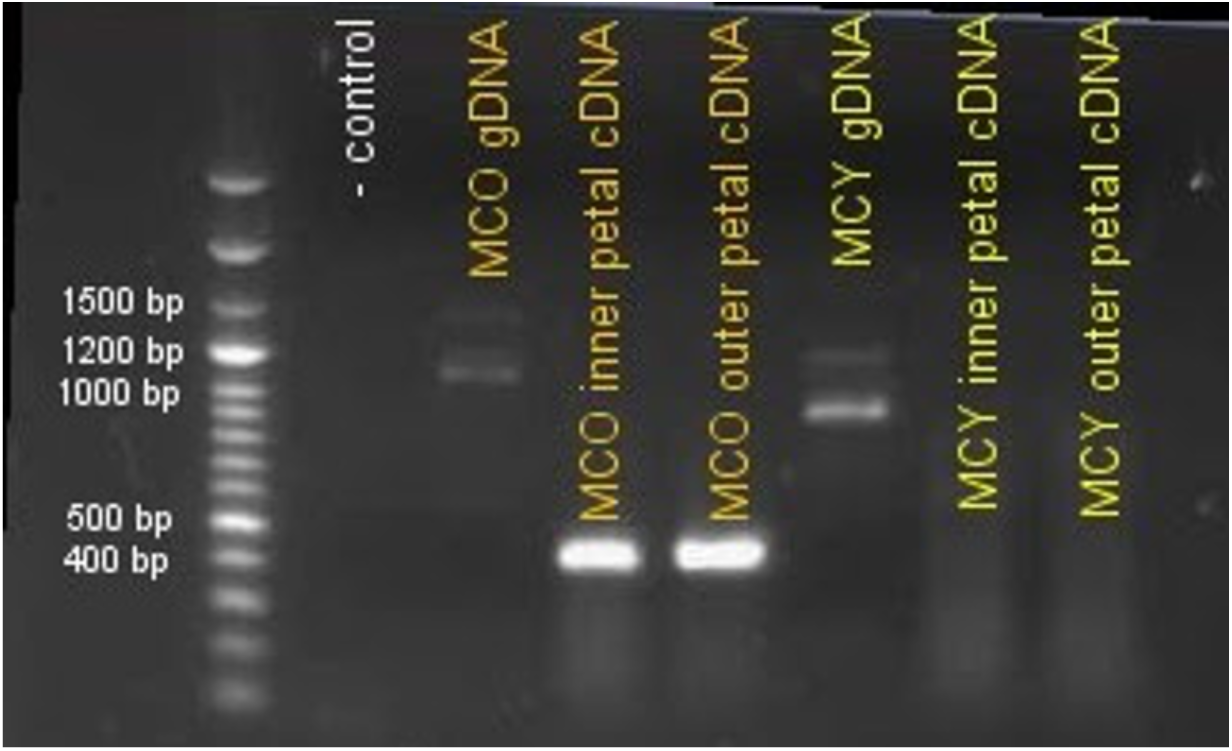
Primers Myb 2_1F and Myb 2b_7R were used to amplify a region of *Myb2* extending from the beginning of the third exon through the beginning of the predicted fourth exon. The expected length of the product spanning the third intron of Myb2 was estimated to be approximately 1050 bp based on the *M. luteus* genome. A PCR amplification done using standard Taq polymerase showed bands at 1050 bp and 1500 bp in orange-flowered *M. cupreus* gDNA and at 950 bp and 1300 bp in yellow-flowered *M. cupreus* gDNA. Amplification using long-amp Taq polymerase revealed that a 950 bp product was also present in gDNA from orange-flowered *M. cupreus*. Sequencing of these products showed that the 1050 bp fragment from orange *M. cupreus* corresponds to *Myb2*, while the two bands seen in yellow *M. cupreus* contain sequences with no resemblance to any anthocyanin-related Myb gene. In orange-flowered *M. cupreus* cDNA from both inner and outer petal, a product of approximately 430 bp was amplified. This indicates that the transcript is spliced as expected, and is expressed in inner and outer petal tissue of orange-flowered *M. cupreus.* No product was observed in the amplification out of yellow-flowered *M. cupreus* petal cDNA, indicating that the Myb 2b transcript is not expressed in yellow *M. cupreus* petals.

**Supplemental Figure S5.**
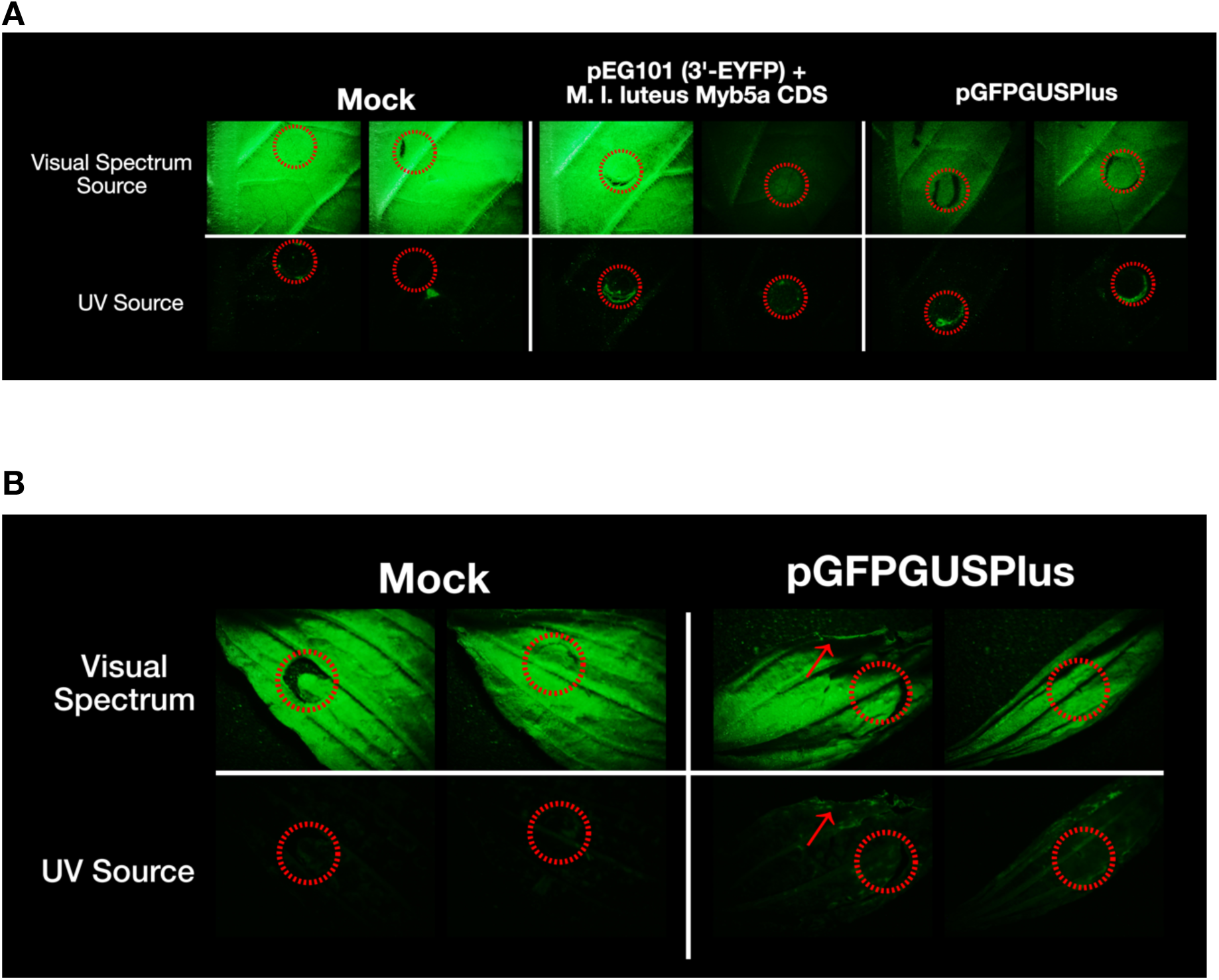
Screen for GFP fluorescence in transfected N. tabacum and M. lewisii leaves. Fluorescence microscopy was performed on a Jenco GL7-3L zoom stereo microscope (Jenco International, Portland, OR, USA) equipped with a NIGHTSEA green filter (NIGHTSEA LLC, Lexington, MA, USA). Light was provided by both a full-spectrum light source supplied with the microscope as well as an external NIGHTSEA ultraviolet light (UV) light source, which was used to excite fluorescence in tissue infiltrated with GFP-expressing plasmids. Leaves infiltrated with pGFPGUSPlus were used as a positive control; leaves infiltrated with a mock control and YFP-expressing pEARLEYGATE101 with M. l. luteus Myb5a CDS were used as negative controls. A) Fluorescence microscopy with Nicotiana tabacum leaves. Left, two replicates treated with 5% sucrose; middle, two replicates treated with pEARLEYGATE101 with M. l. luteus Myb5a CDS in Agrobacterium tumefaciens; right, two replicates treated with pGFPGUSPlus in A. tumefaciens. Top row, visual spectrum light source; bottom row, UV source. Some leaves imaged with visual spectrum light source appear dimmer; this is an artifact caused by the inadvertent use of a different light setting. Red circles correspond to injection sites. B) Fluorescence microscopy with Mimulus lewisii leaves. Left, two replicates treated with 5% sucrose; right, two replicates treated with pGFPGUSPlus in A. tumefaciens. Top row, visual spectrum light source; bottom row, UV source. Red circles correspond to injection sites. Note apparent tissue damage at edge of leaves treated with pGFPGUSPlus (red arrows).

**Supplemental Figure S6.**
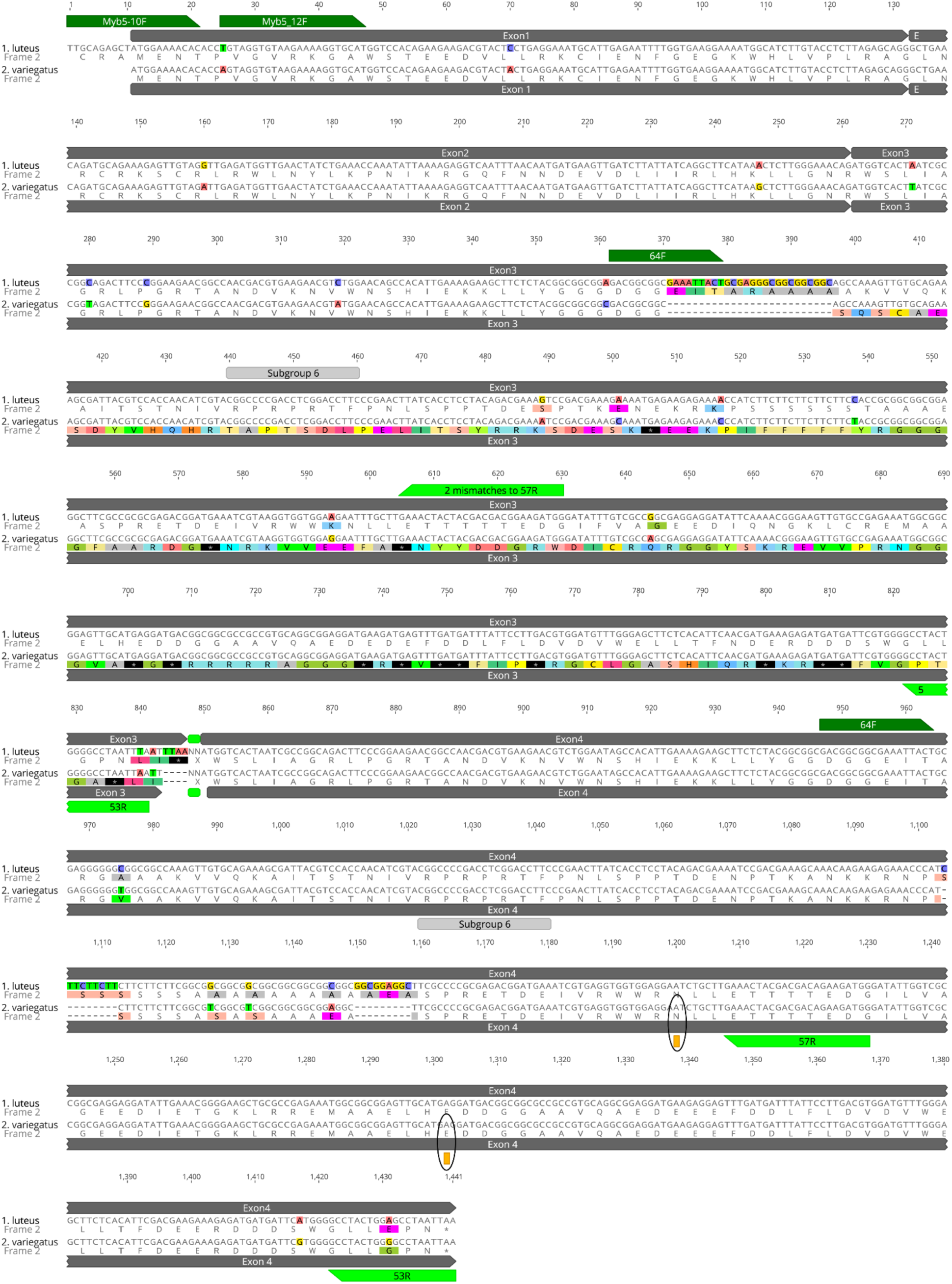
Alleles of *MYB5a* from the yellow-petaled *M. l. luteus* (top sequence on each line) and the magenta-petaled (anthocyanin-pigmented) *M. l. variegatus* (bottom sequence on each line). Primers mentioned in the Methods are shown in dark green (forward primers) and light green (reverse primers). Exons are marked in gray. Two putative A-to-I editing sites are marked in yellow. Note that Subgroup 6, which is a hallmark of all known anthocyanin-activating R2R3 MYBs, is present in both taxa in exon 4 as the amino acid sequence RPRPRTF. In exon 3, however, a frameshift mutation in *M. l. variegatus* eliminates this feature, making it unlikely that the exon 1-2-3 splice variant of *M. l. variegatus* would be capable of anthocyanin activation.

**Supplemental Figure S7.**
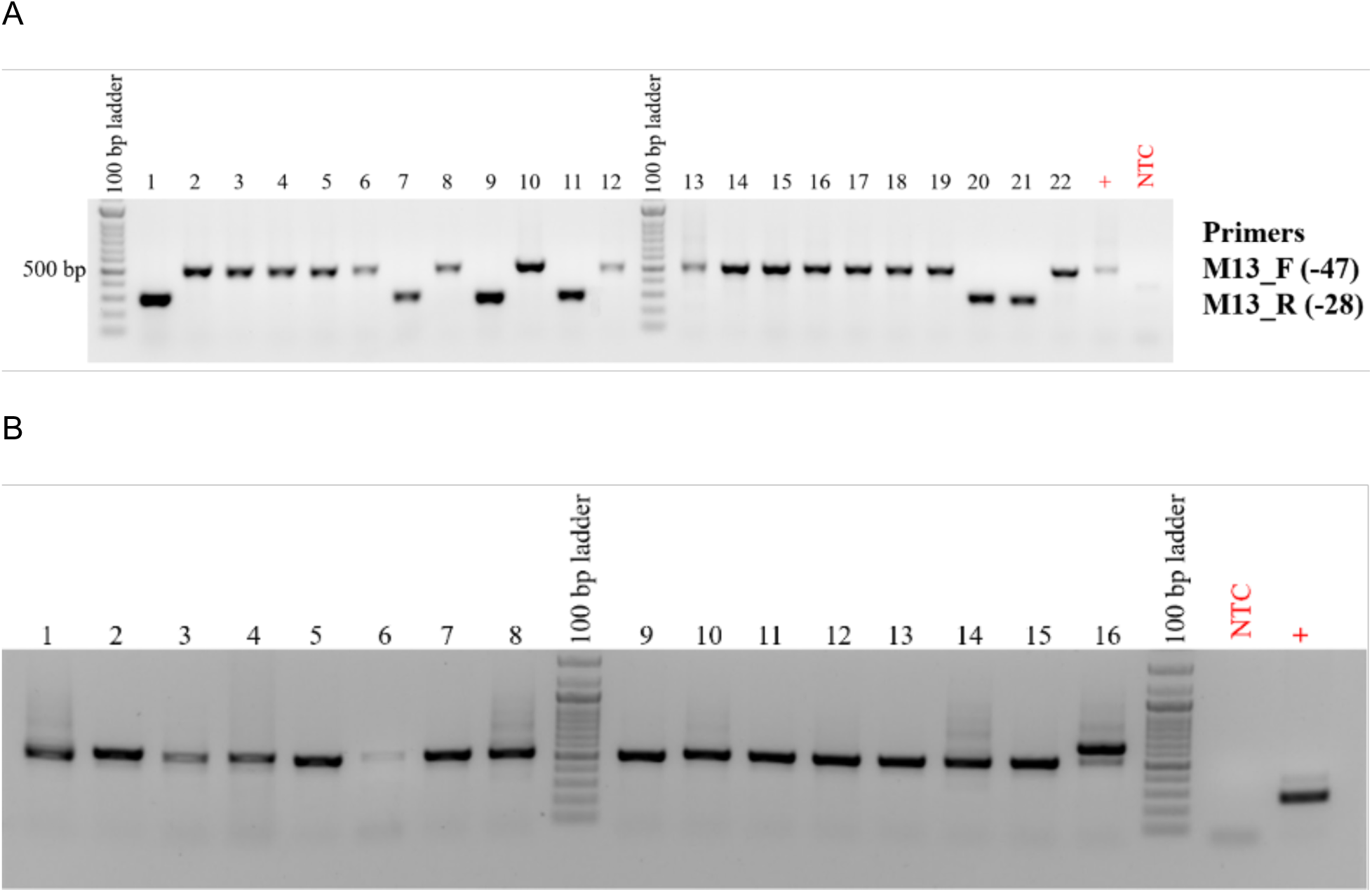
pGEM Cloning of an exon 4 fragment of *MYB5a*. A fragment containing the first edited site was PCR-amplified using primers 64F-57R and cloned, and colonies were screened for fragment insertion. Lanes with band size ∼500 bp are colonies that contain the *MYB5a* exon 4 insert. Lanes with bands ∼300 denote empty vectors. (A) Primers M13_F (-47) and M13_R (-28) were used to screen white colonies from a pGEM cloning attempt with *M. l. variegatus* x *M. l. luteus* F1 hybrid gDNA. 16 colonies appear to have taken up the MYB5 exon 4 insert. (B) Primers M13_F (-47) and M13_R (-28) were used to screen white colonies from a pGEM cloning attempt with Mlv gDNA. 15 colonies appear to have successfully taken up the insert. The band at ∼650bp most likely denotes a vector with a longer, incorrect insert.

**Supplemental Figure S8.**
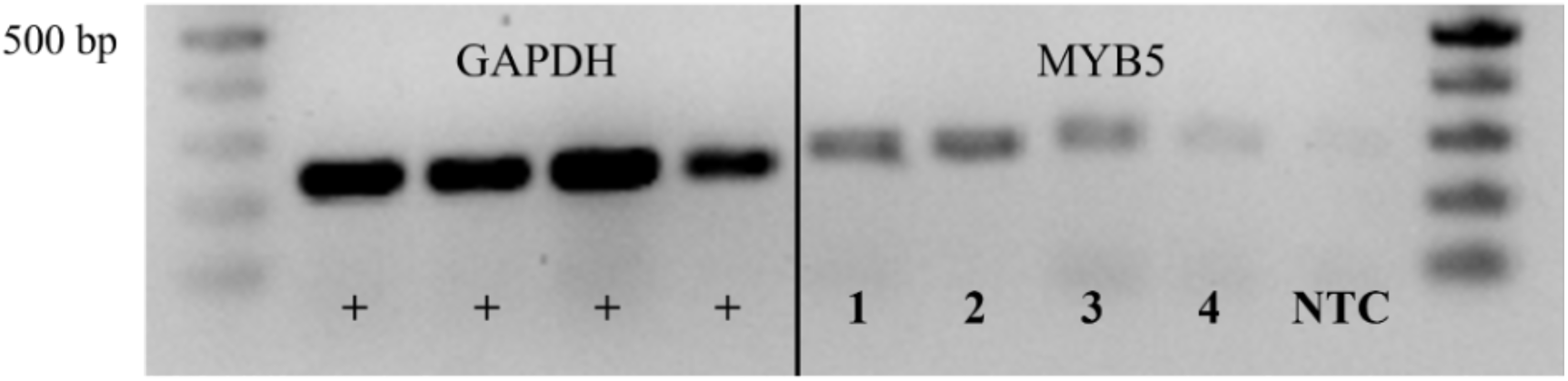
Relative expression of *MYB5a* in *M. l. variegatus* and *M. l. luteus*. Left: positive control using GAPDH primers 1F-2R, on each of the samples shown on the right. Right: An exon 4 fragment of *MYB5a* amplified using Myb5_64F and Myb5_57R from cDNA from *M. l. variegatus* inner petal (1) and outer petal (2); the red-spotted *M. l. luteus* inner petal (3); and the yellow *M. l. luteus* outer petal (4). NTC, No Template Control. All band sizes were as expected.

**Supplemental Figure S9.** Leaf photos utilized in quantitative comparisons of anthocyanin production. Photos excluded due to leaf damage are not included. A. pGFP negative controls, with two transgene injections per leaf. B. Leaves for which the *M. l. variegatus MYB5a* exon 1-2- 4 transgene was injected on the left side of the leaf and the *M. l. luteus* allele on the right. C. Leaves for which the *M. l. variegatus MYB5a* exon 1-2-4 transgene was injected on the right side of the leaf and the *M. l. luteus* allele on the left. Files uploaded separately.

