## Supplementary figures and images for "Coding-sequence evolution does not explain divergence in petal anthocyanin pigmentation between *Mimulus luteus* var. *luteus* and *M. l. variegatus*"

### Supplemental Figure S9.A

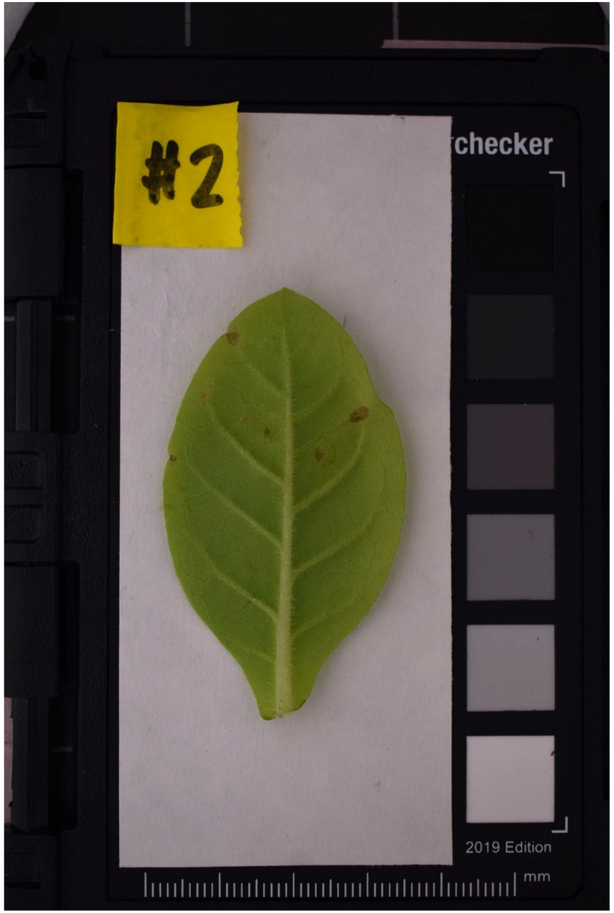

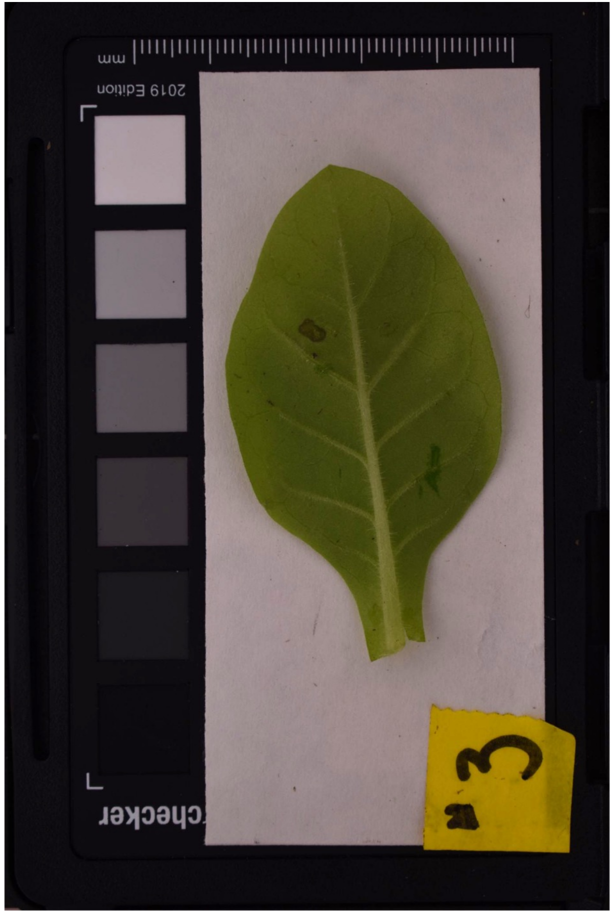

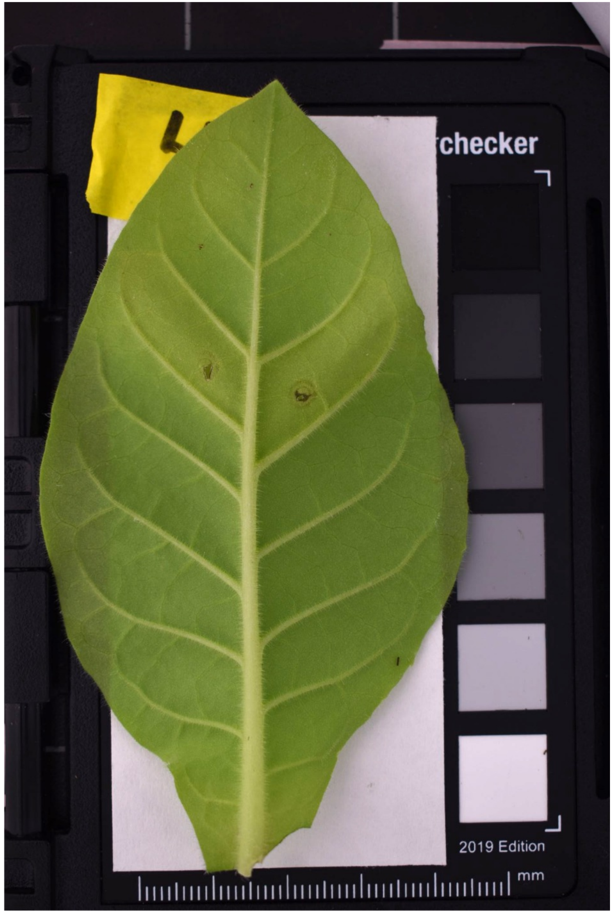

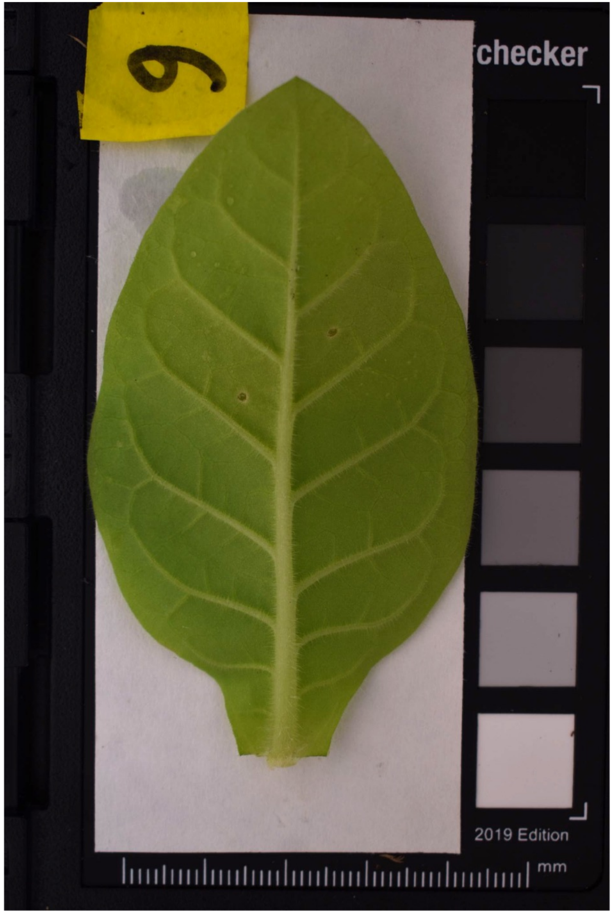

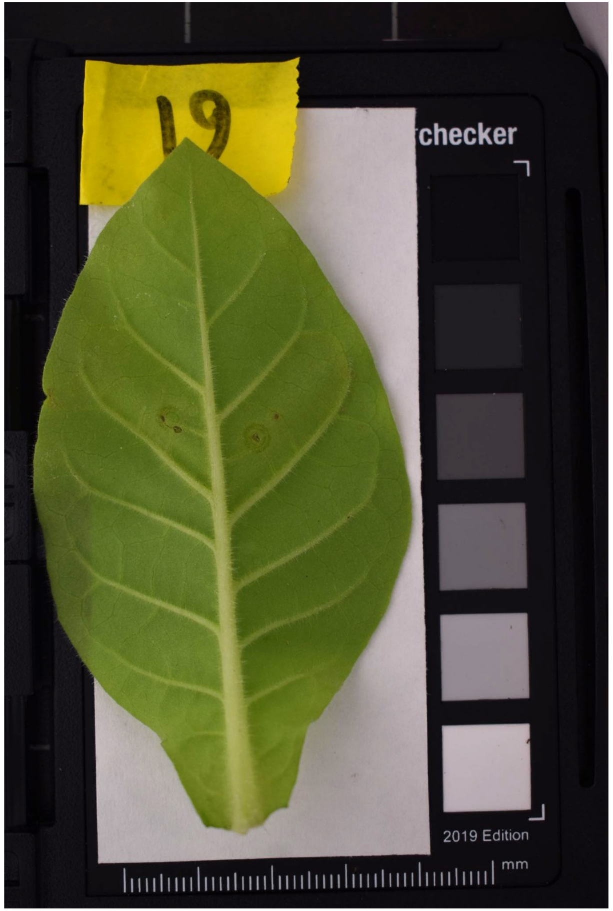

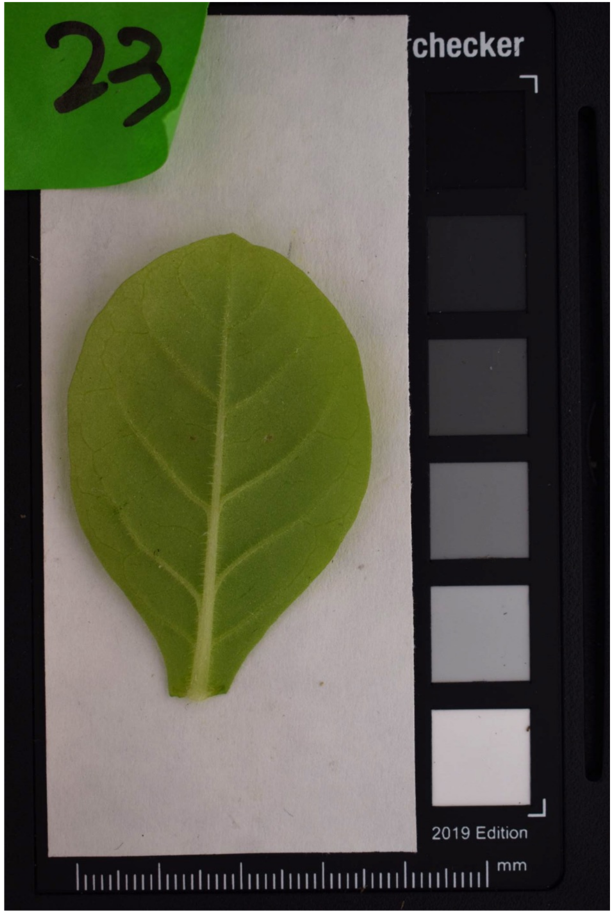

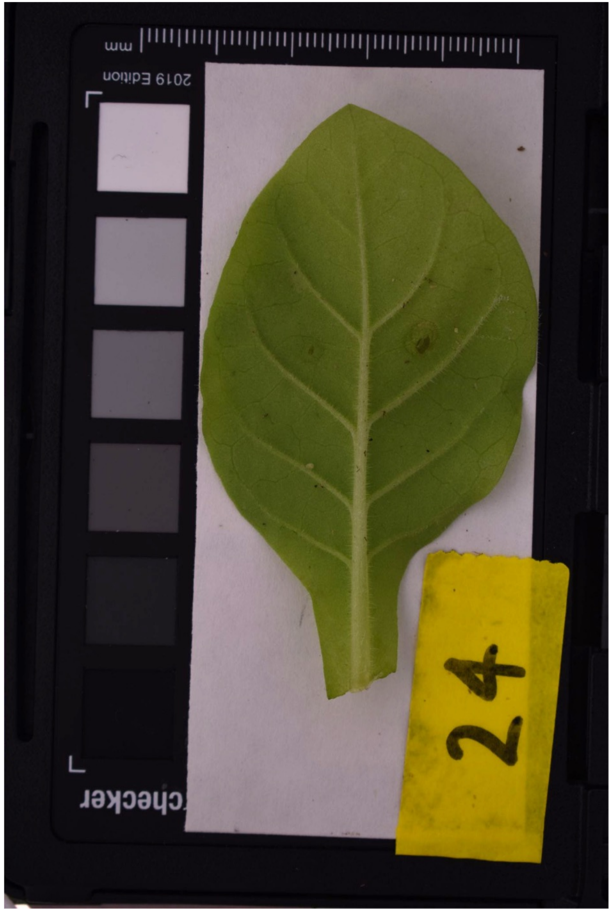

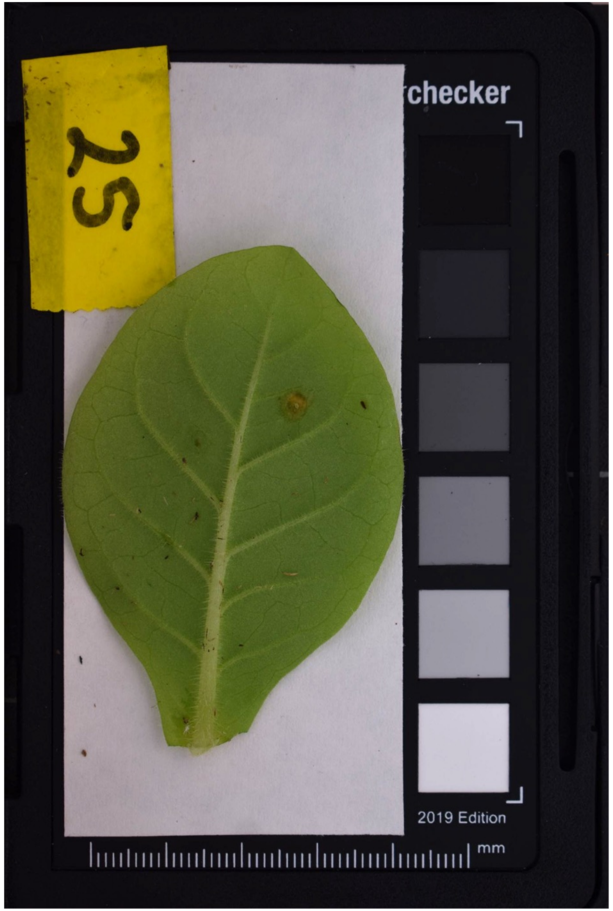

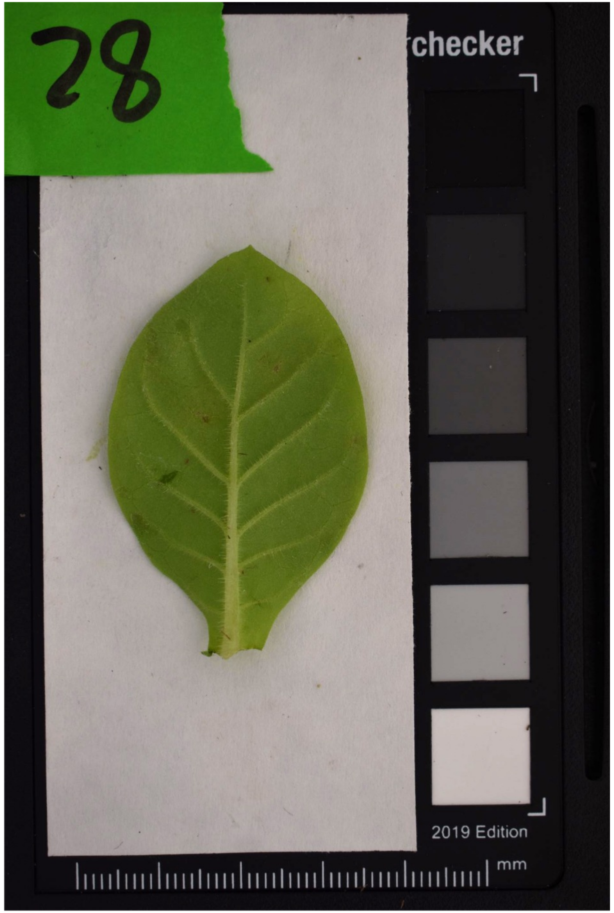

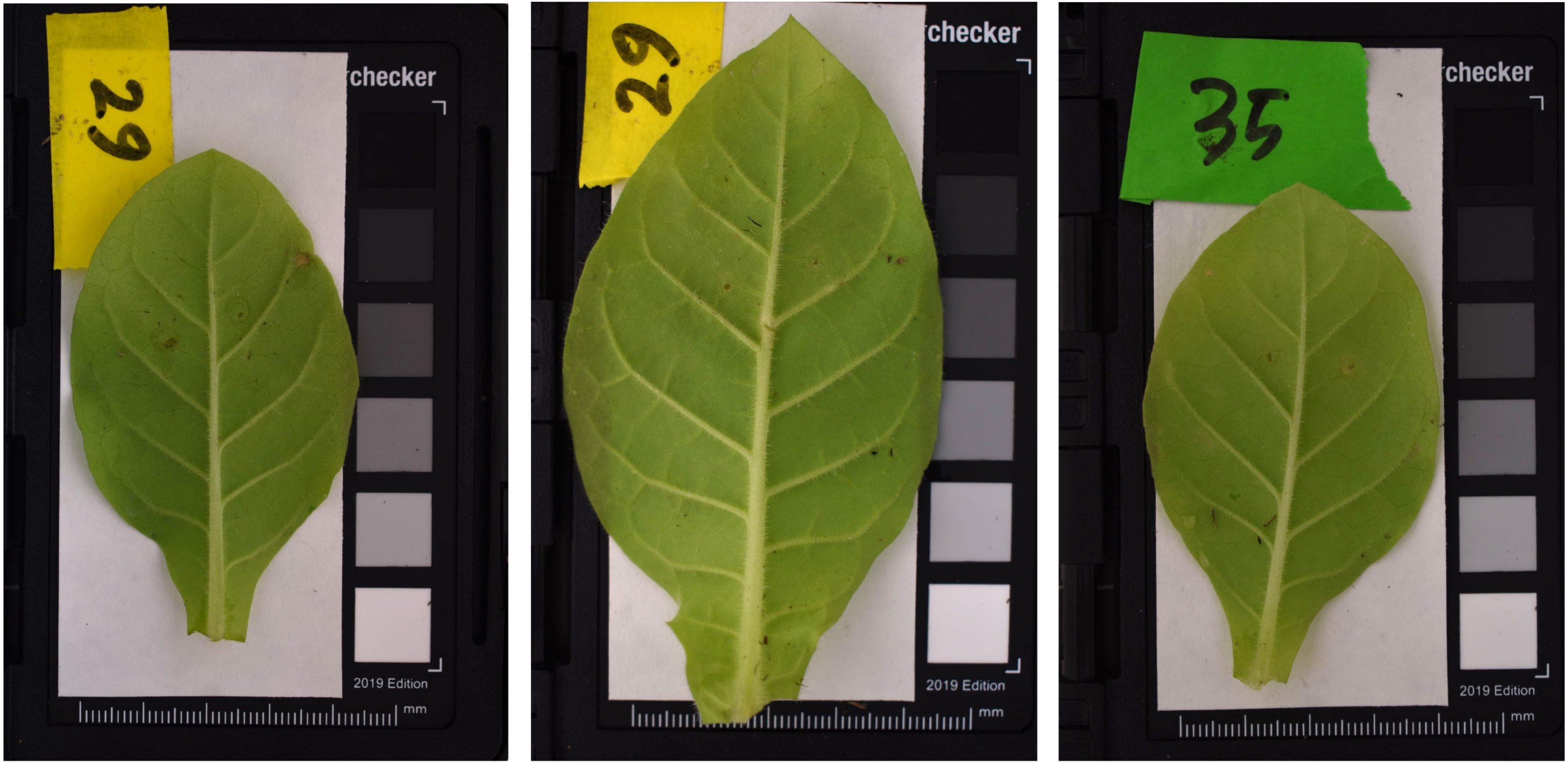

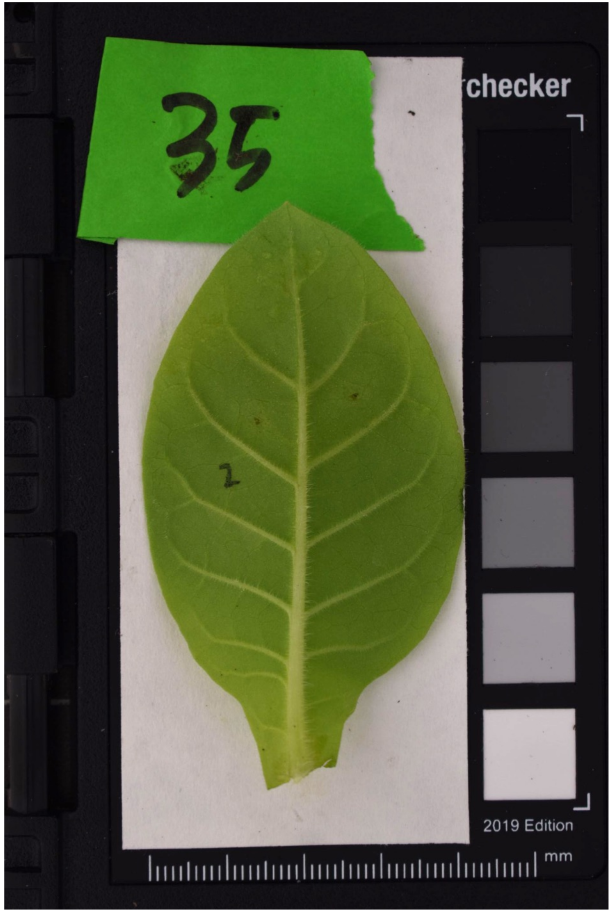

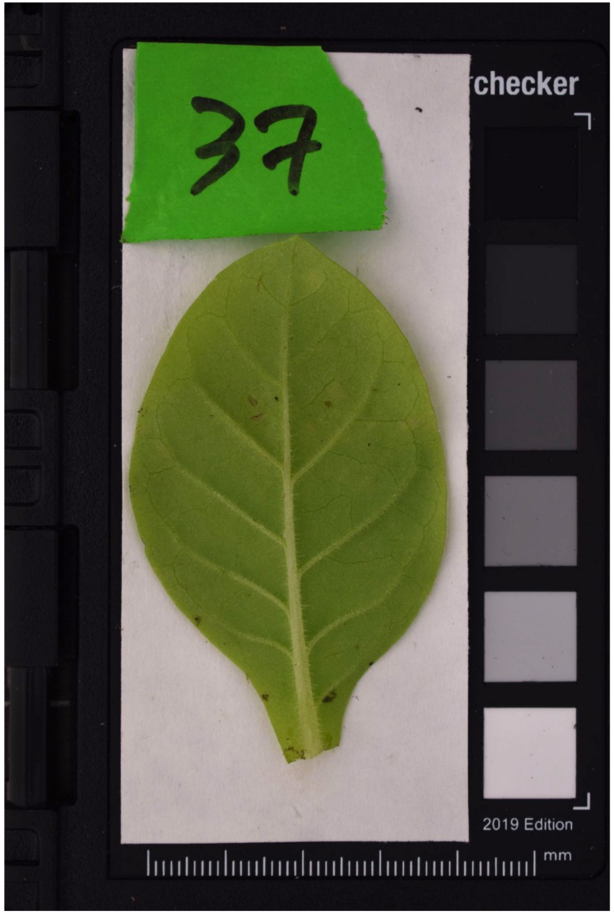

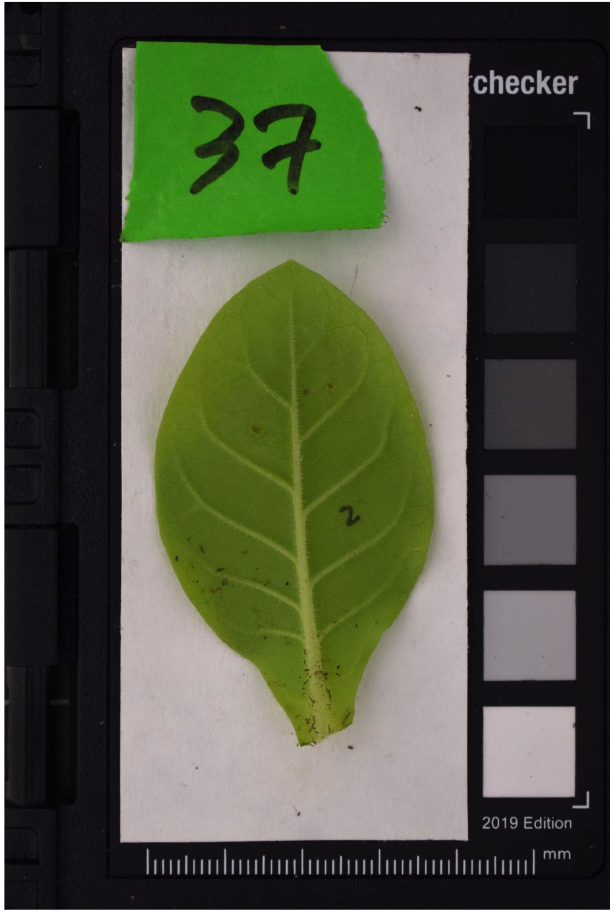

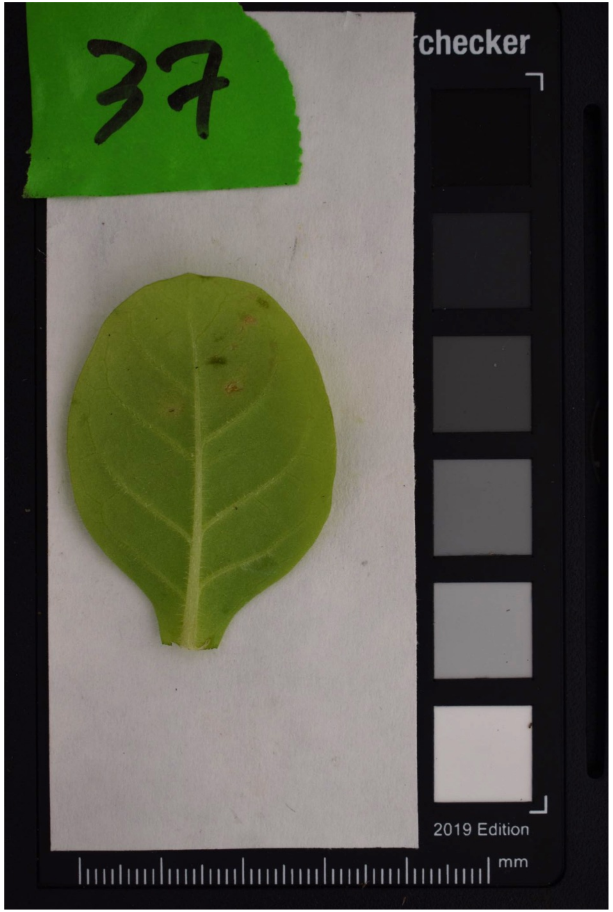

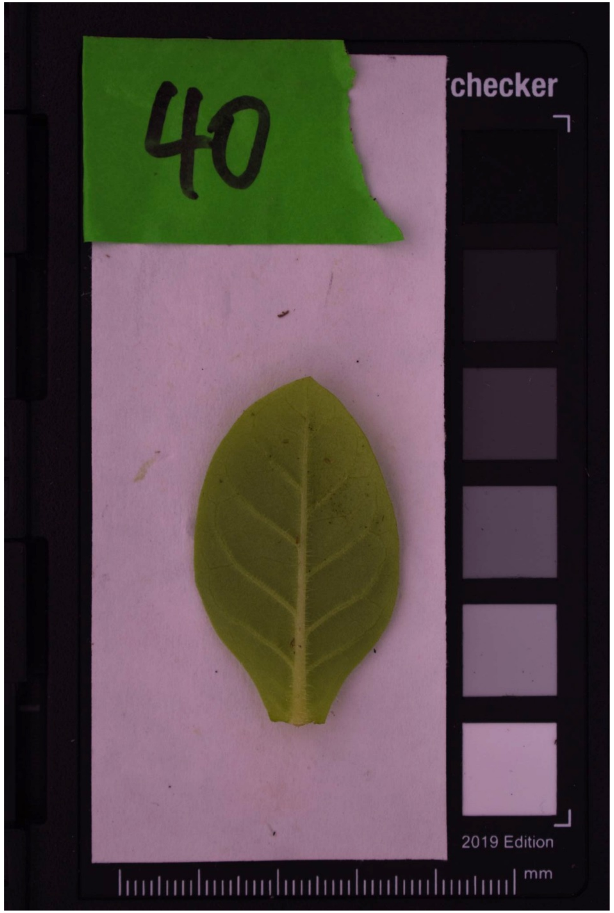

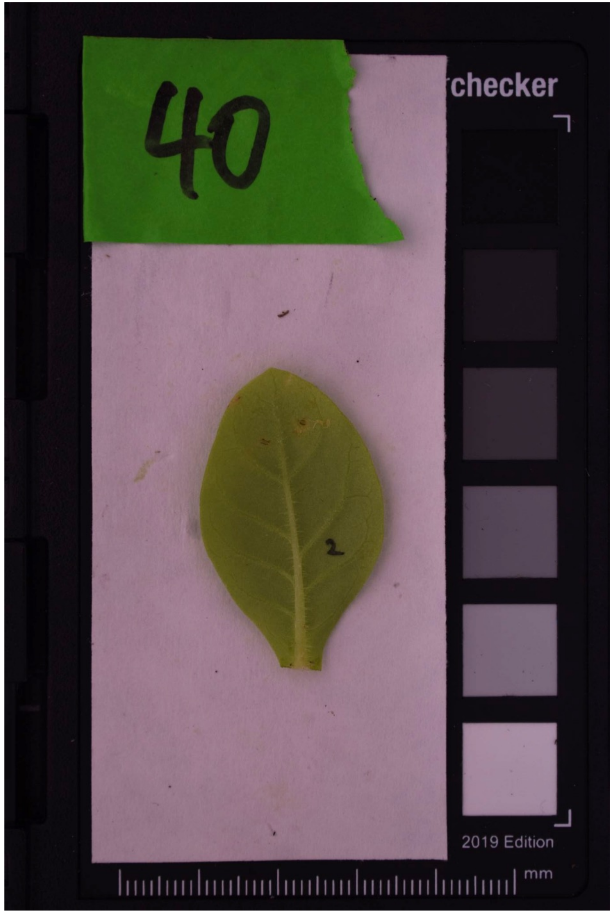

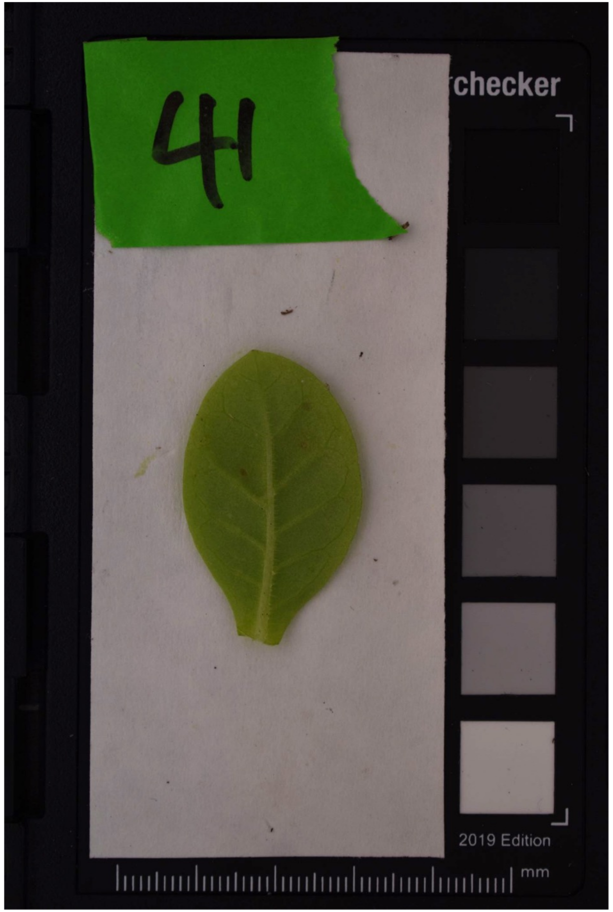

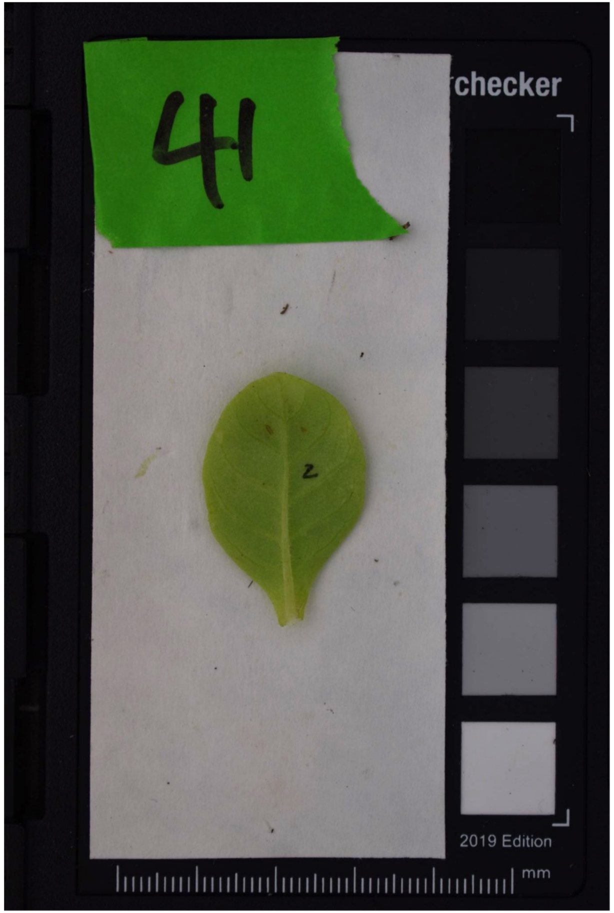
