## Supplemental Figure S9.B for "Coding-sequence evolution does not explain divergence in petal anthocyanin pigmentation between *Mimulus luteus* var. *luteus* and *M. l. variegatus*"

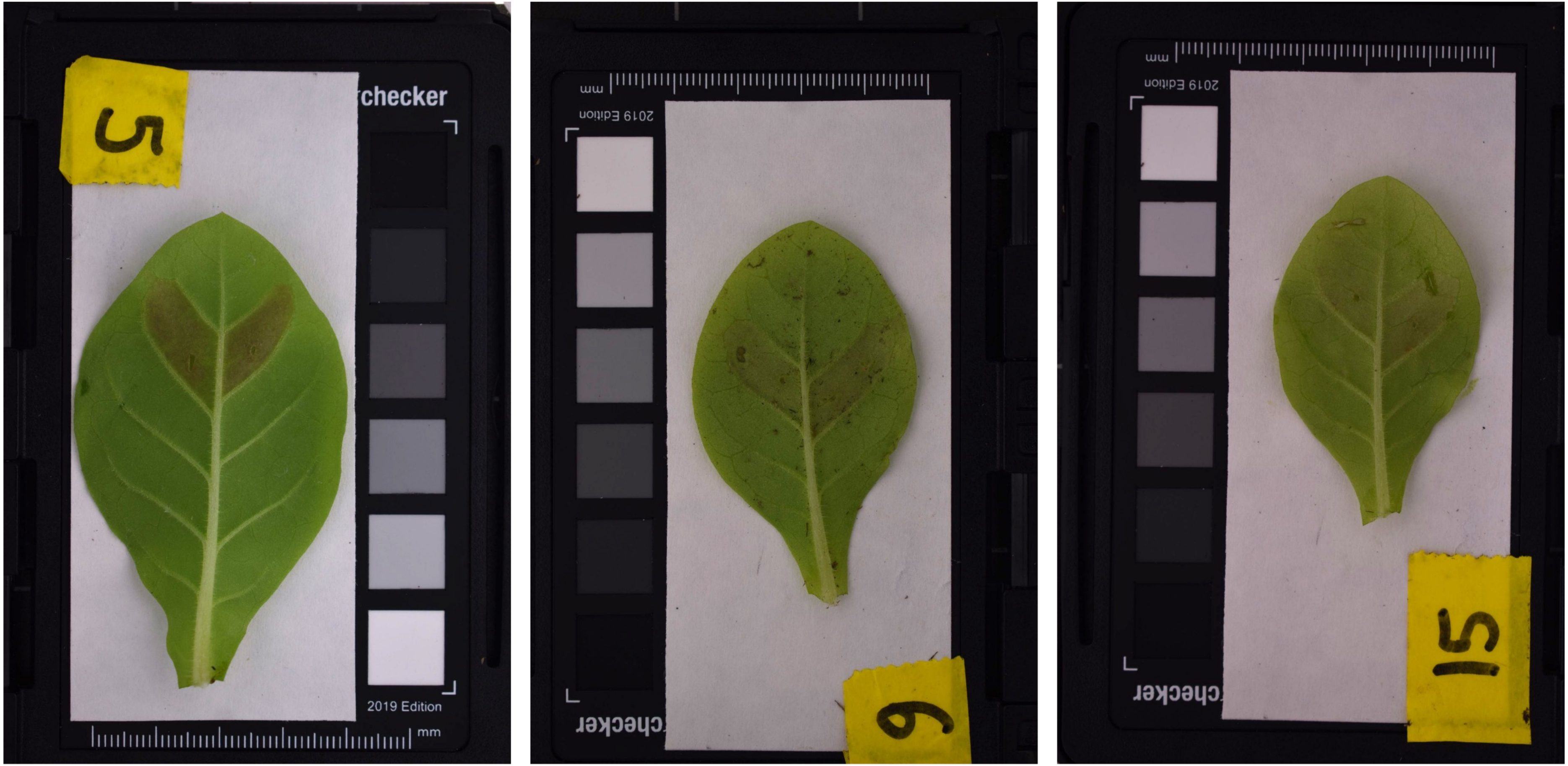
5
checker
2019 Edition
mm

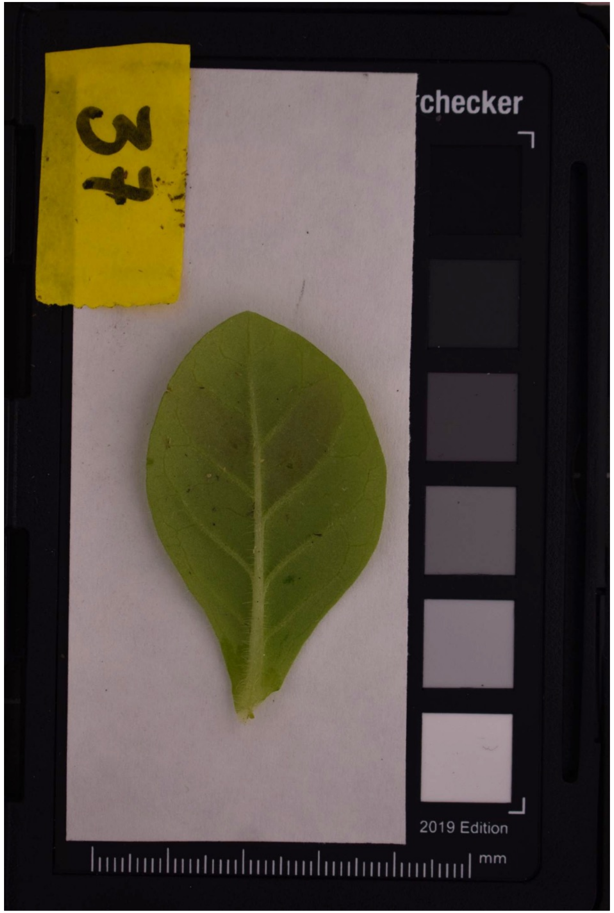
37
checker
2019 Edition
mm

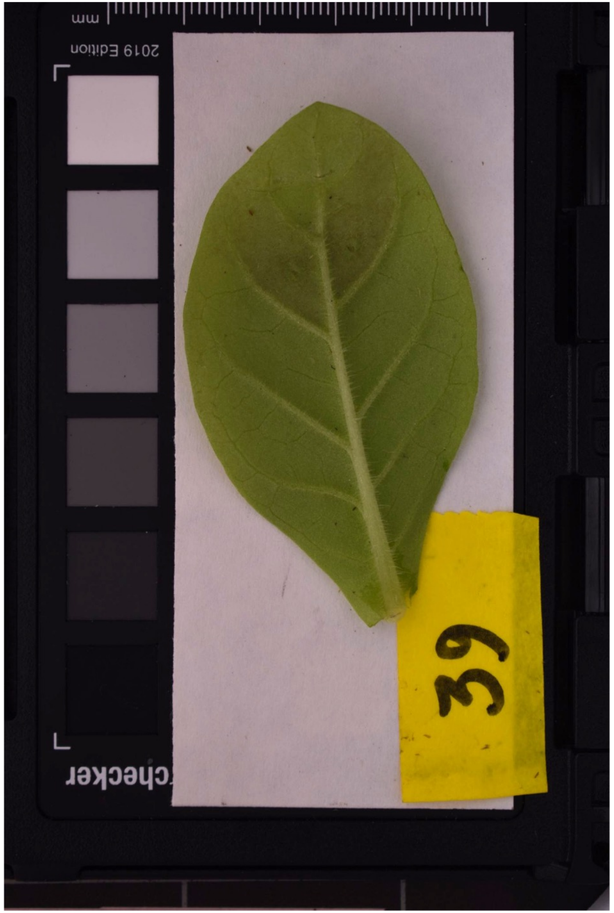
39
checker
2019 Edition
mm

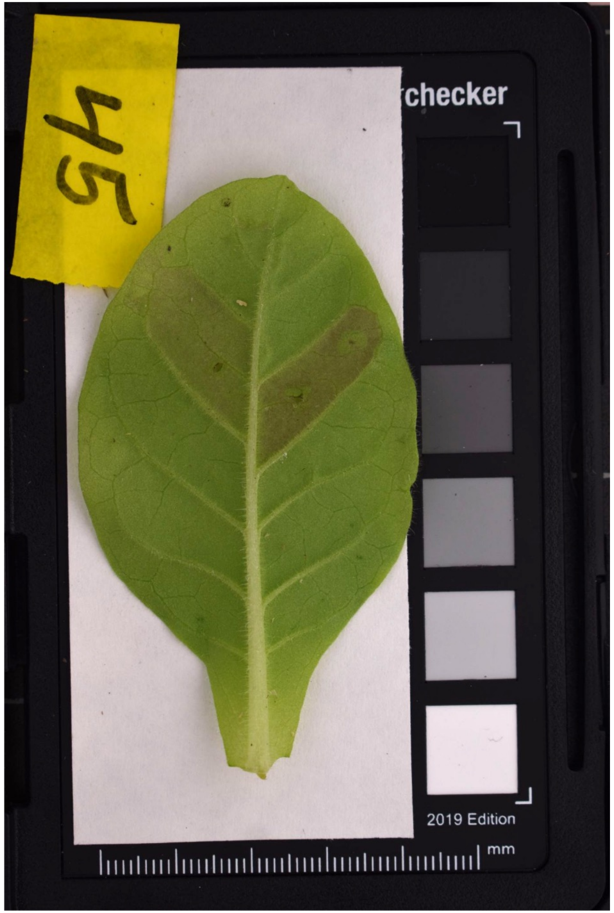
45
checker
2019 Edition
mm

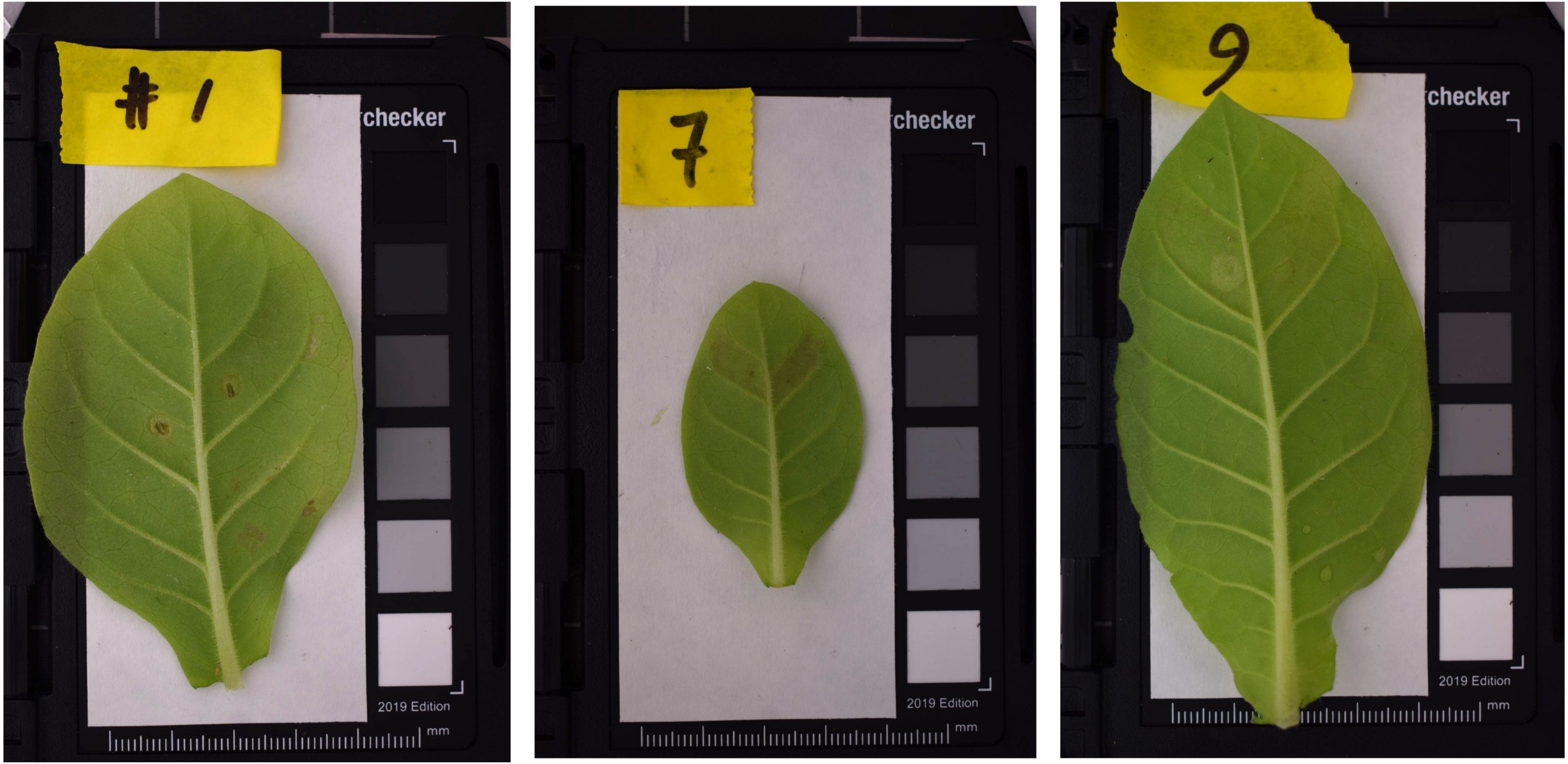
#1
2019 Edition
mm
7
2019 Edition
mm
9
2019 Edition
mm

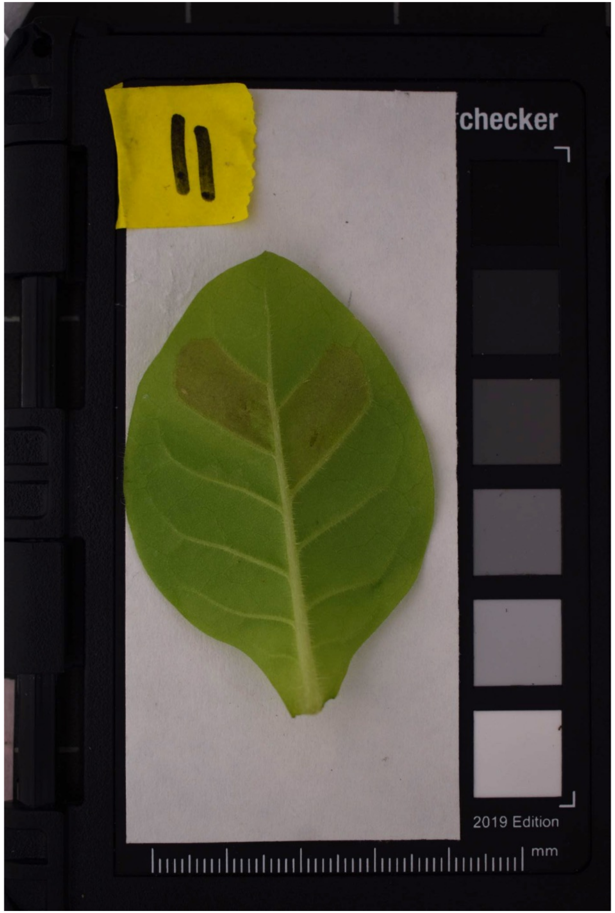
11
checker
2019 Edition
mm

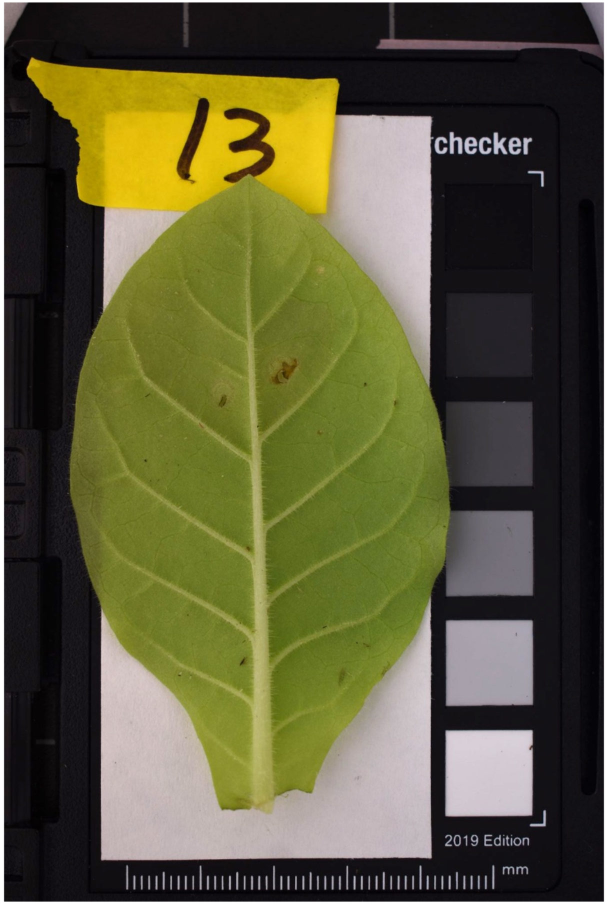
13
checker
2019 Edition
mm

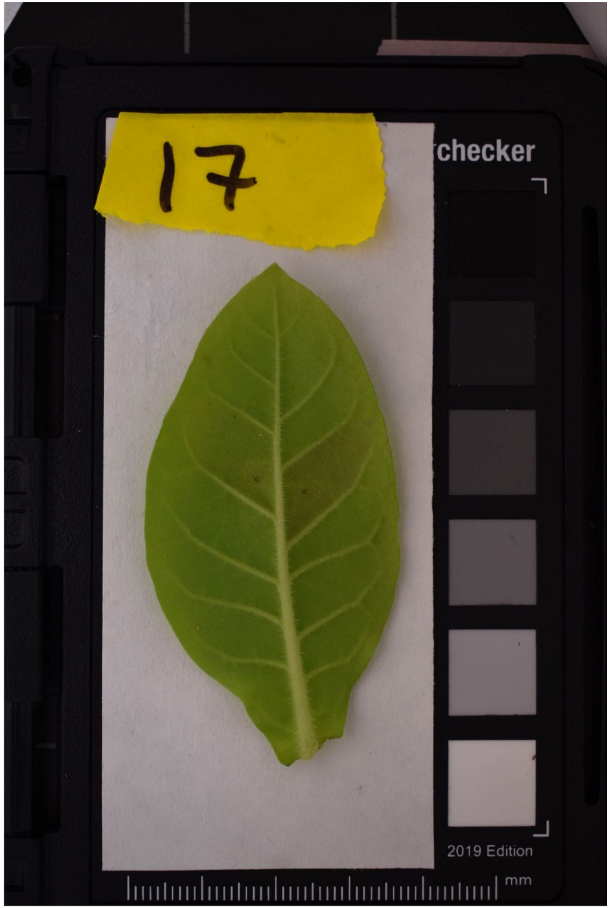
17
checker
2019 Edition
mm

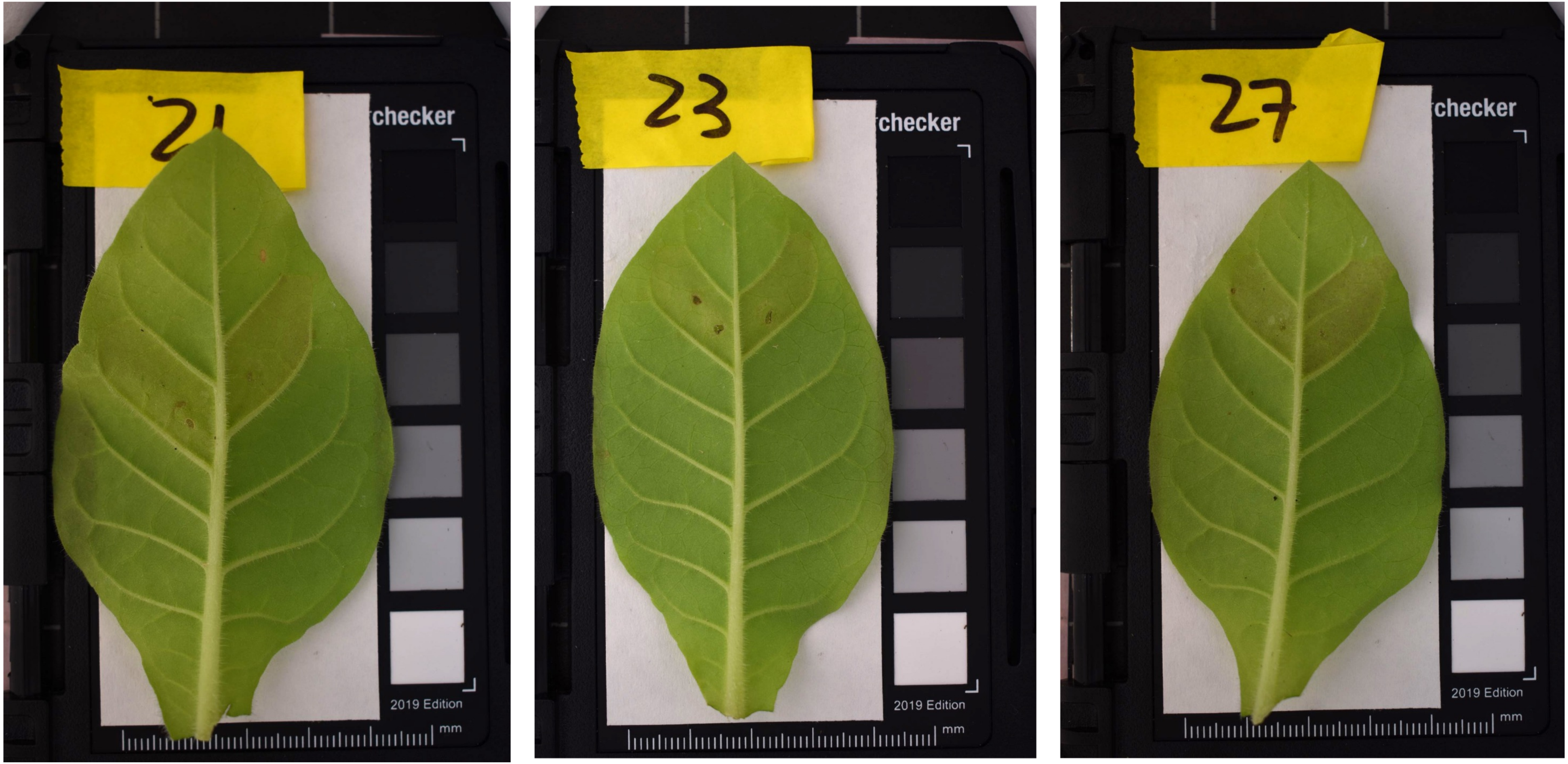
21
checker
2019 Edition
mm
23
checker
2019 Edition
mm
27
checker
2019 Edition
mm

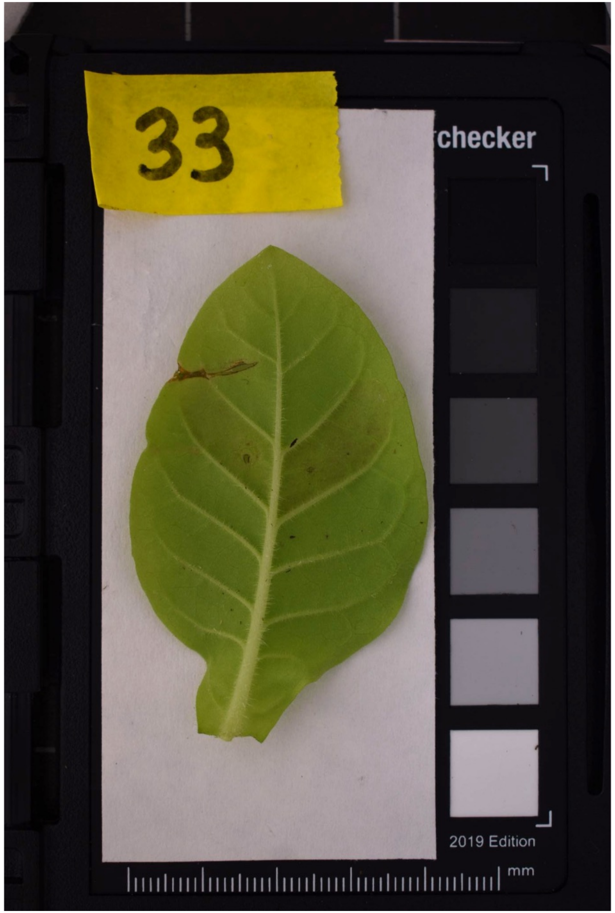
33
checker
2019 Edition
mm
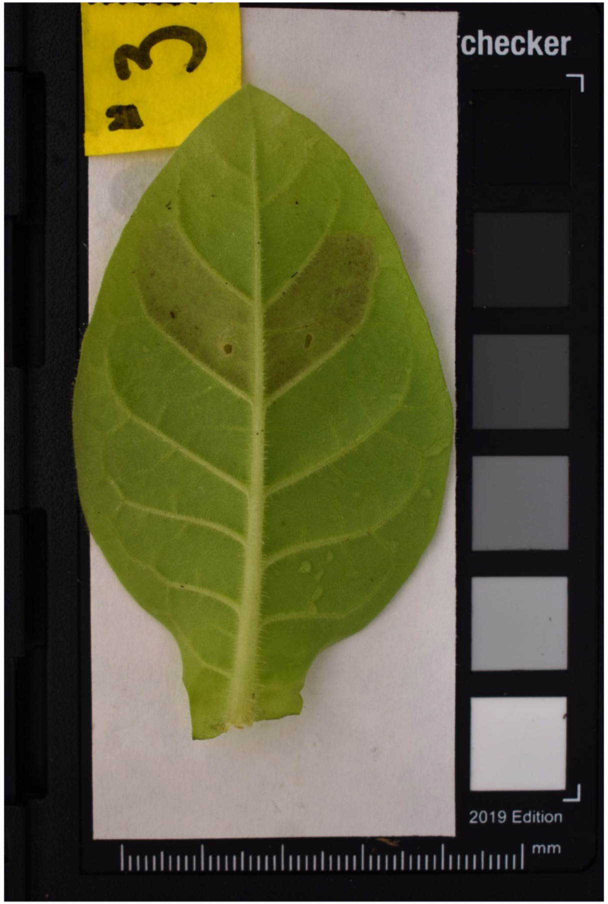
3
checker
2019 Edition
mm
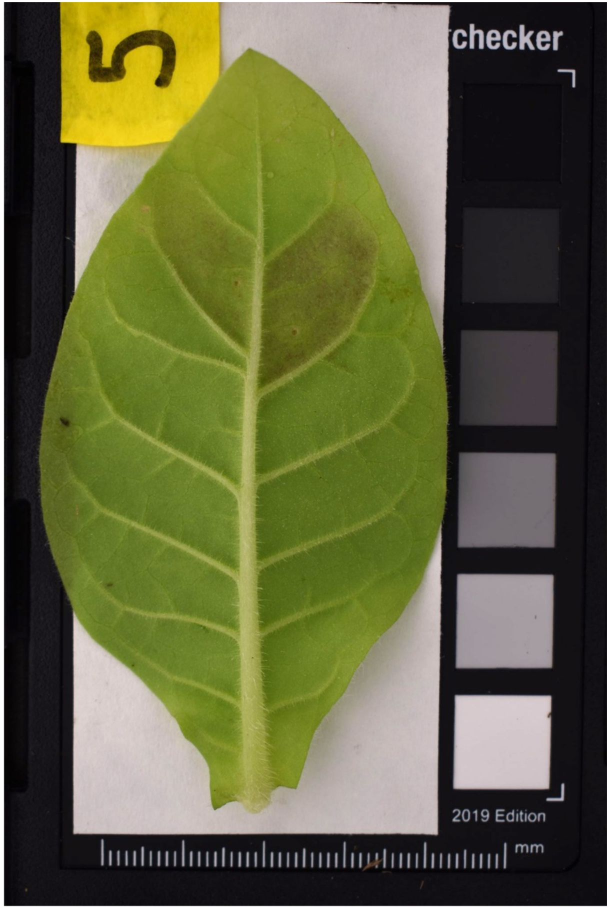
5
checker
2019 Edition
mm

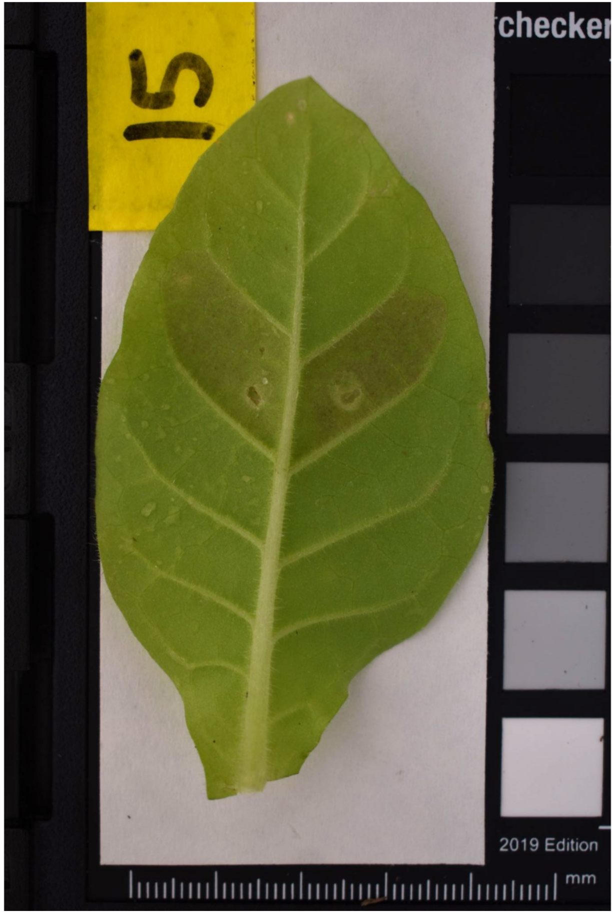
15
checker
2019 Edition
mm
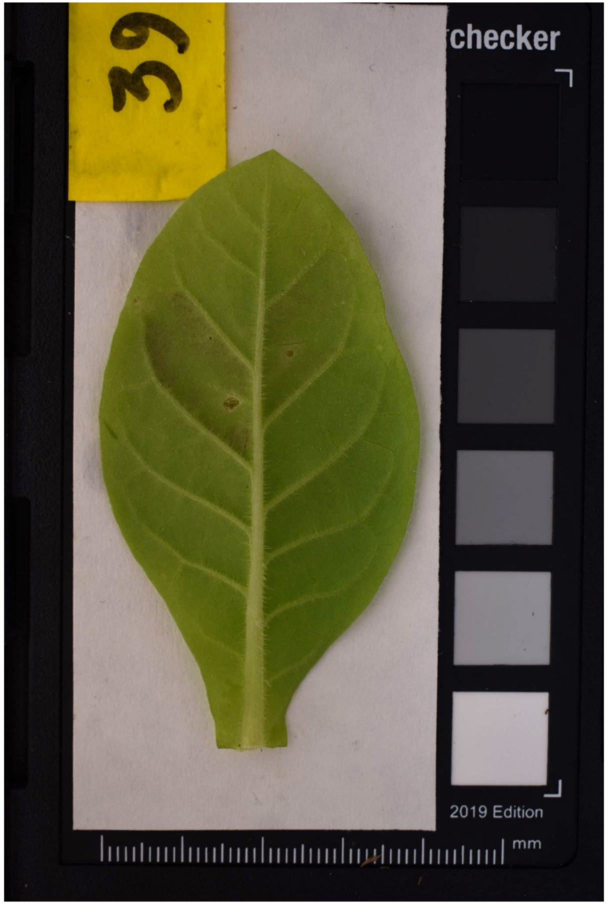
39
checker
2019 Edition
mm
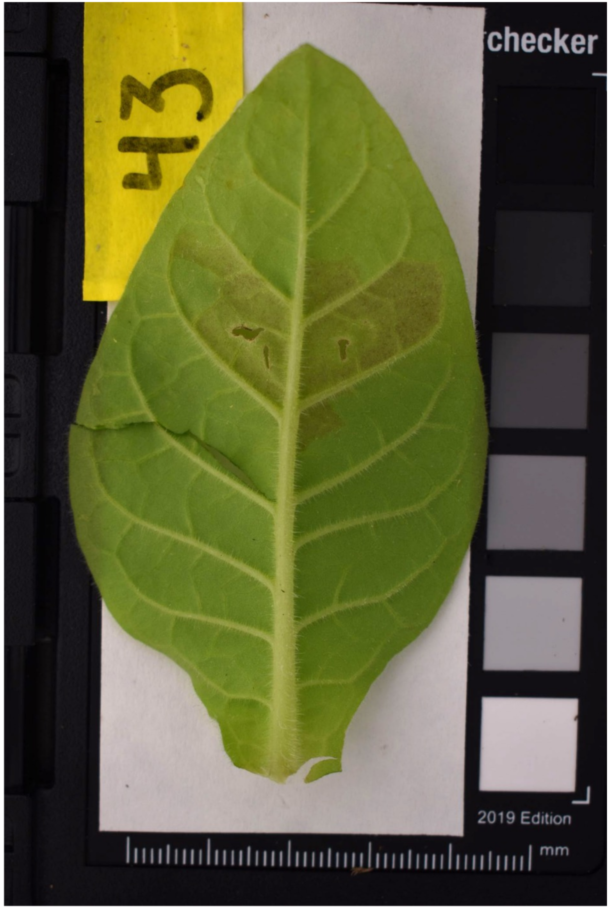
43
checker
2019 Edition
mm

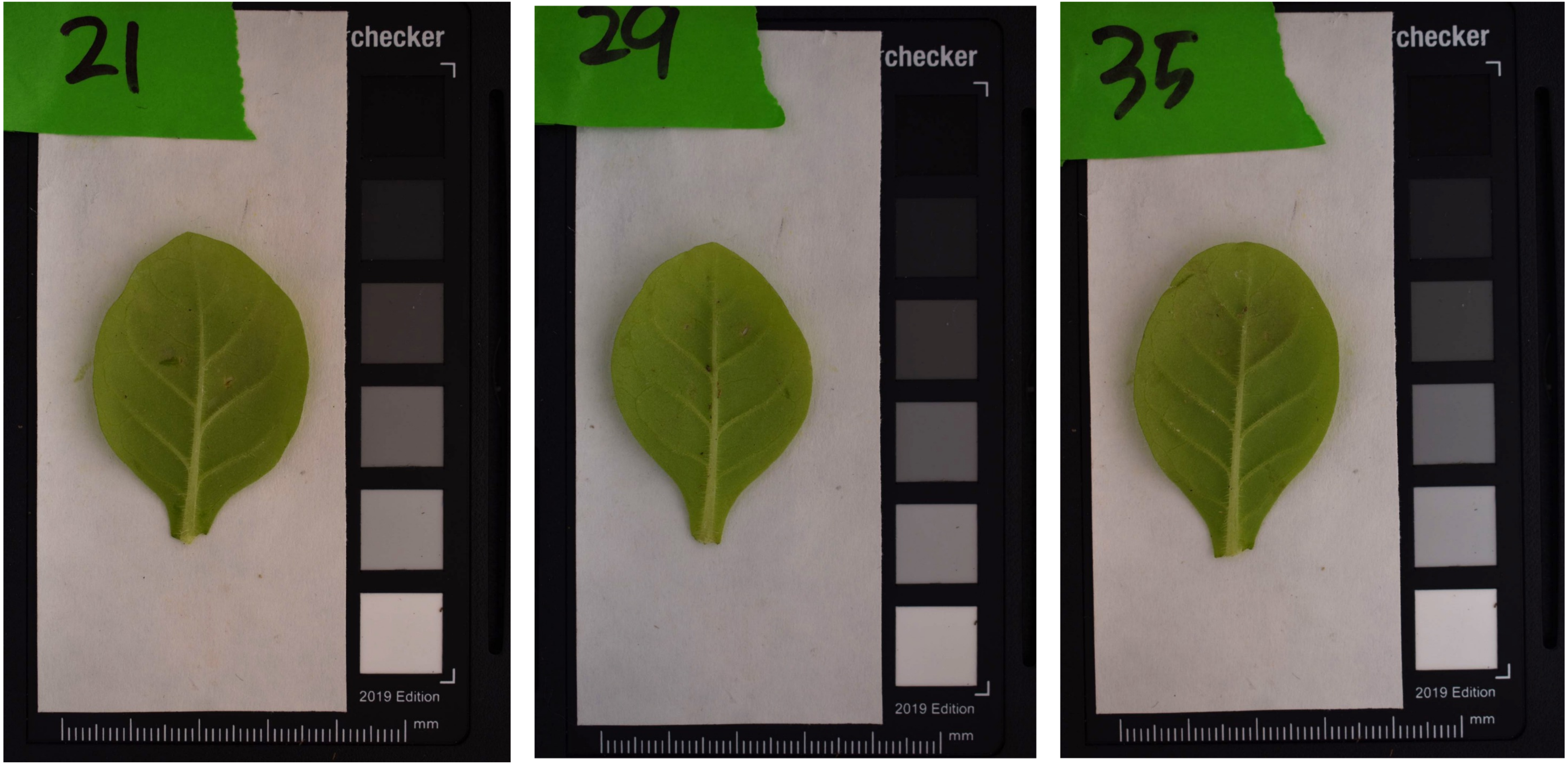
21
checker
2019 Edition
mm
29
checker
2019 Edition
mm
35
checker
2019 Edition
mm

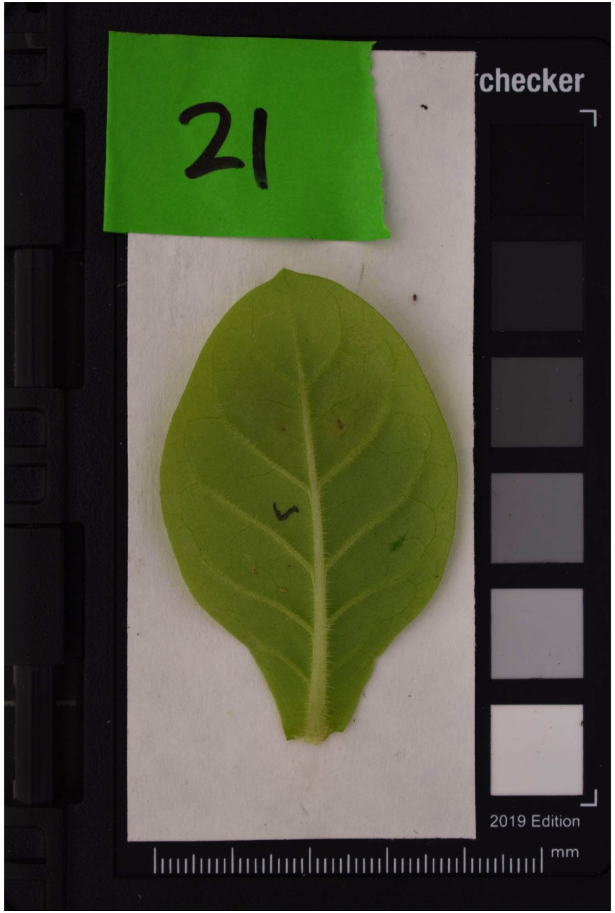
21
checker
2019 Edition
mm
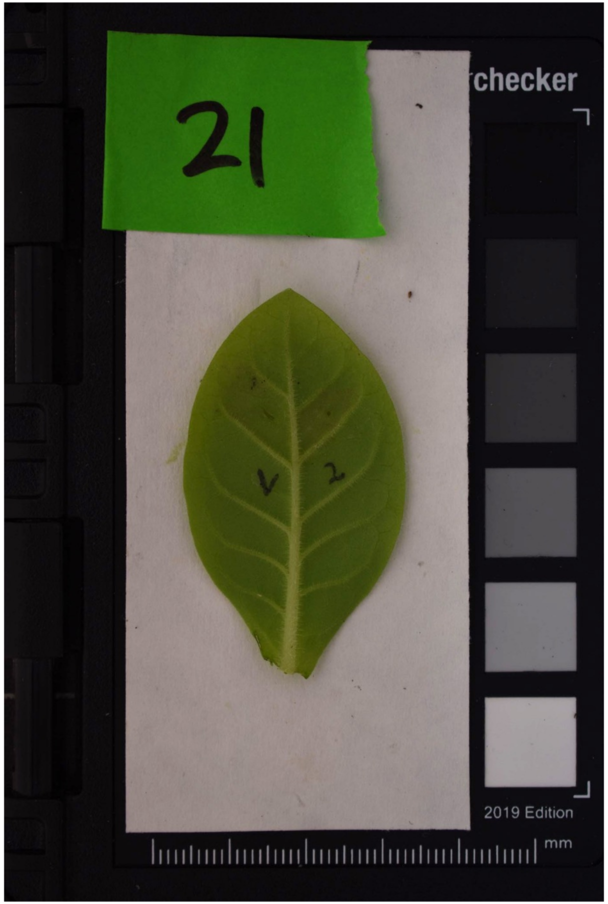
21
checker
2019 Edition
mm
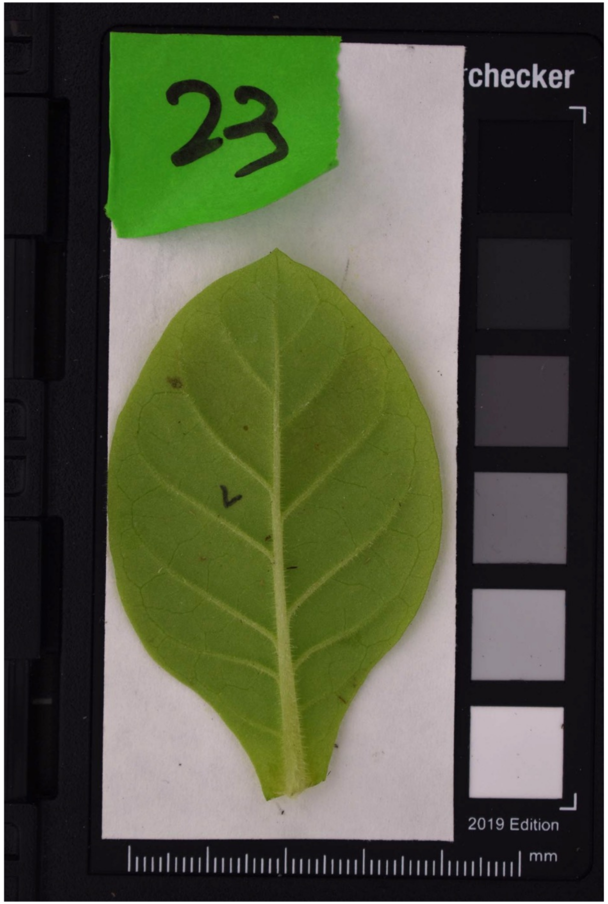
23
checker
2019 Edition
mm

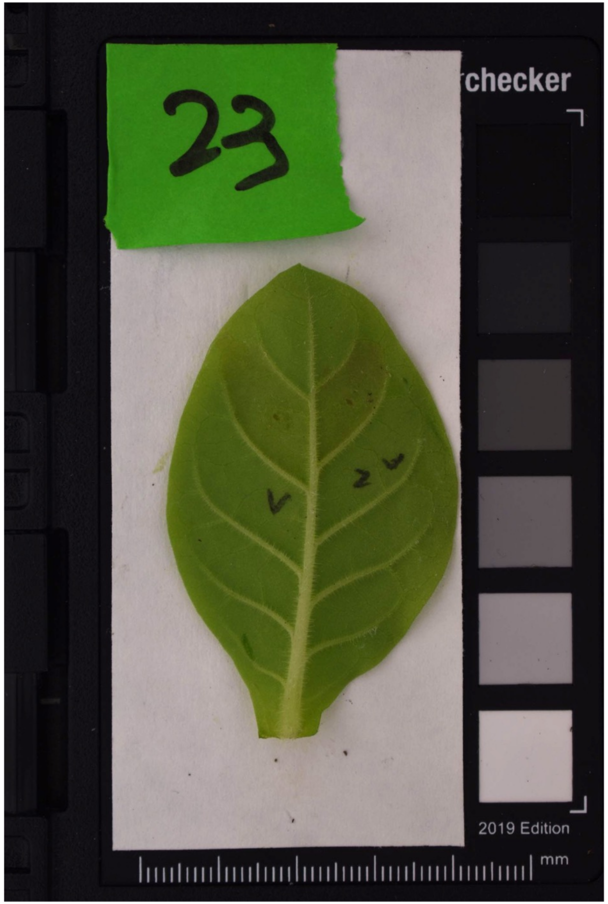
23
checker
2019 Edition
mm
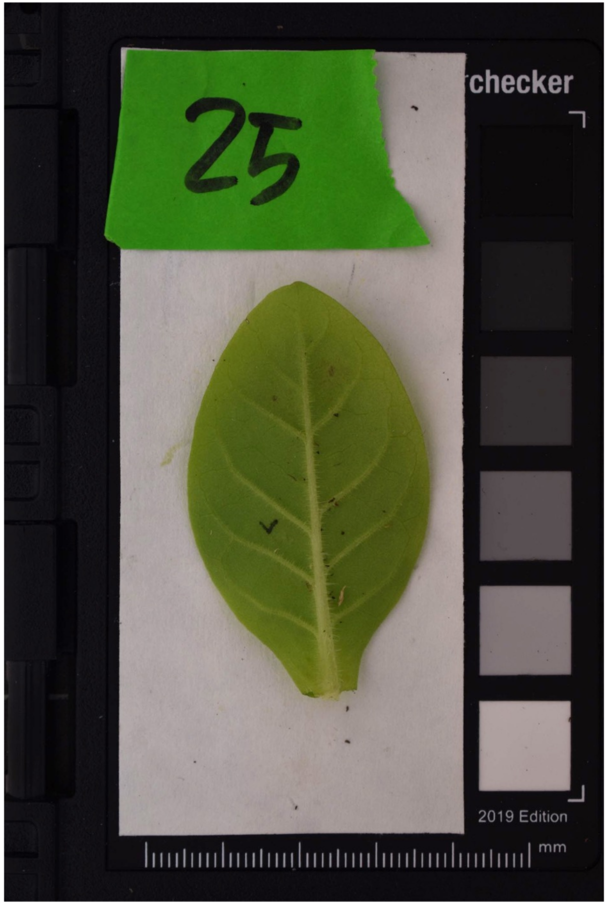
25
checker
2019 Edition
mm
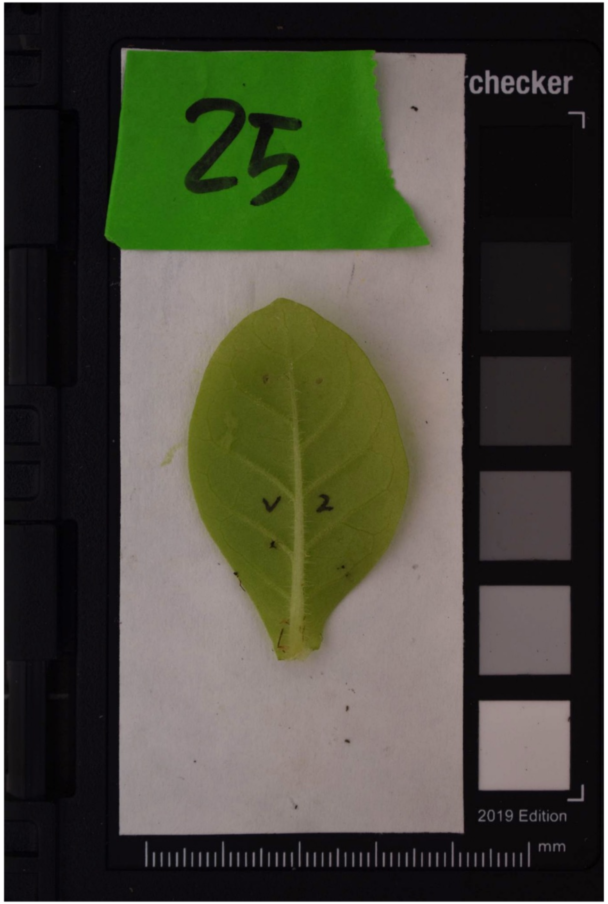
25
checker
2019 Edition
mm

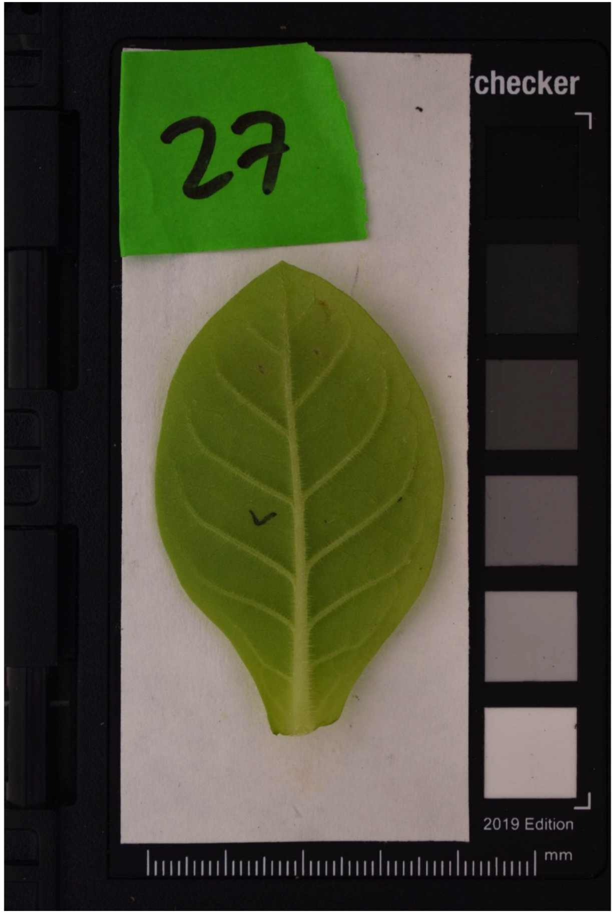
27
checker
2019 Edition
mm

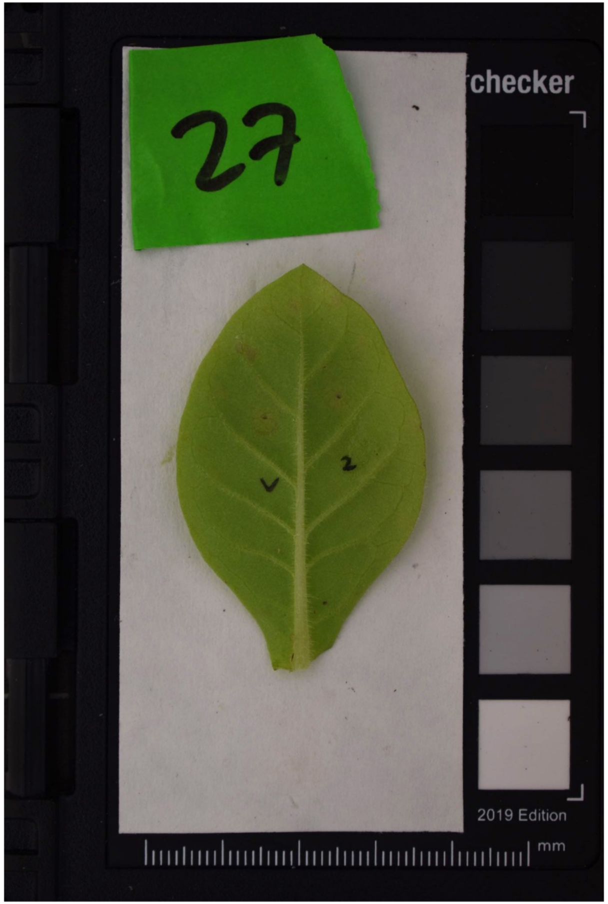
27
checker
2019 Edition
mm

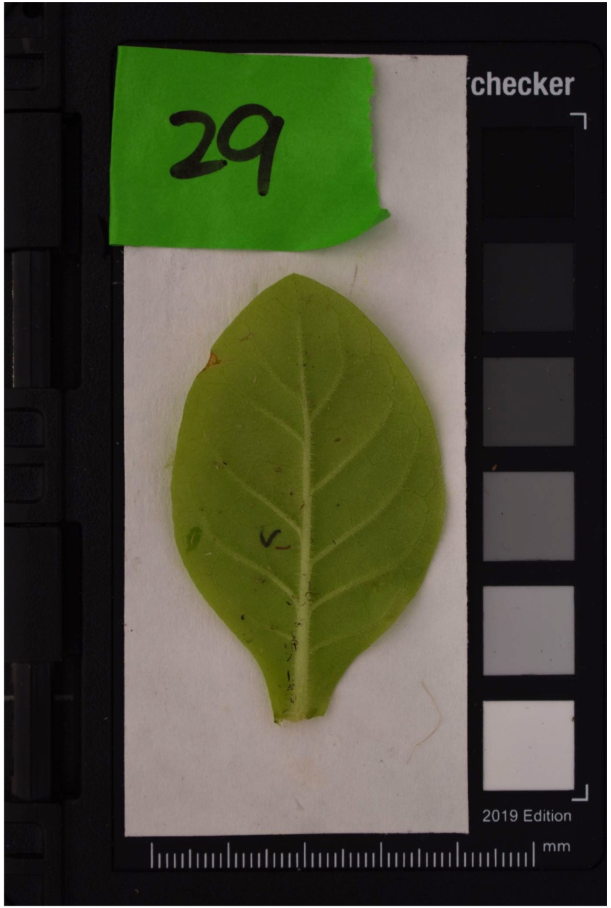
29
checker
2019 Edition
mm

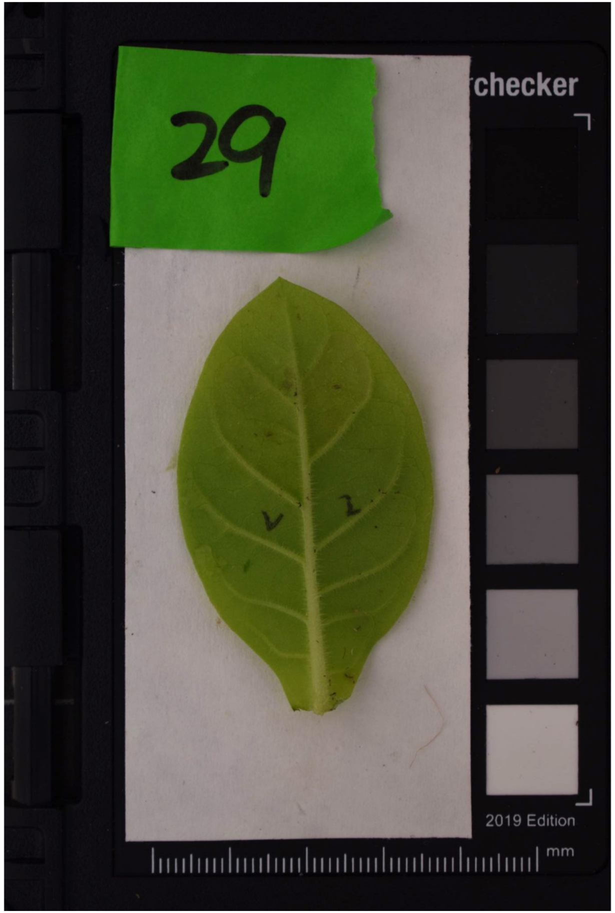
29
checker
2019 Edition
mm

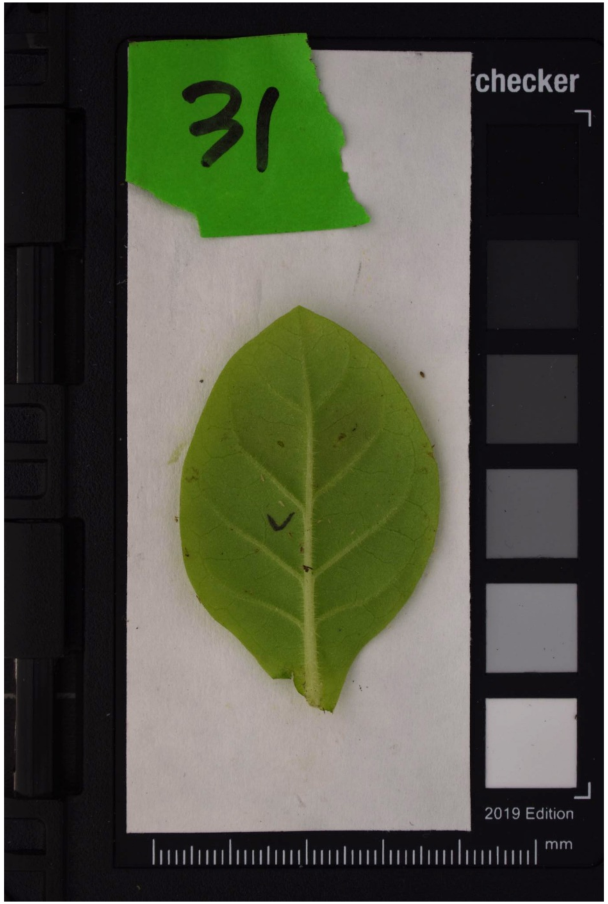
31
checker
2019 Edition
mm

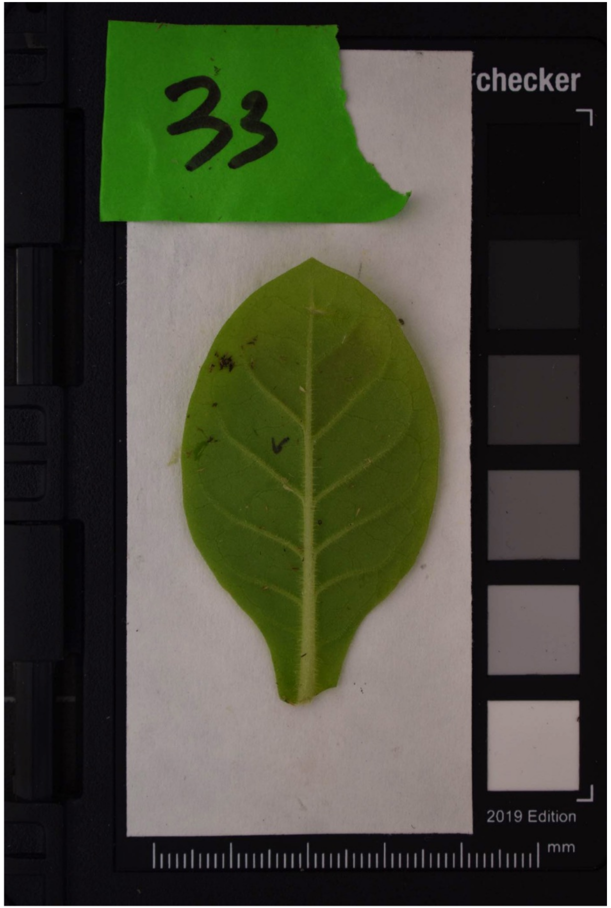
33
checker
2019 Edition
mm

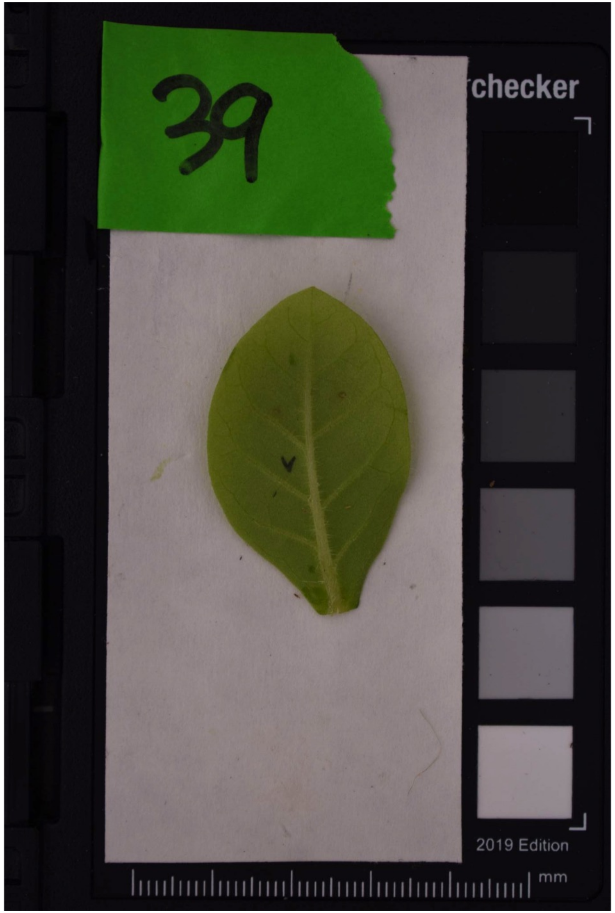
39
checker
2019 Edition
mm
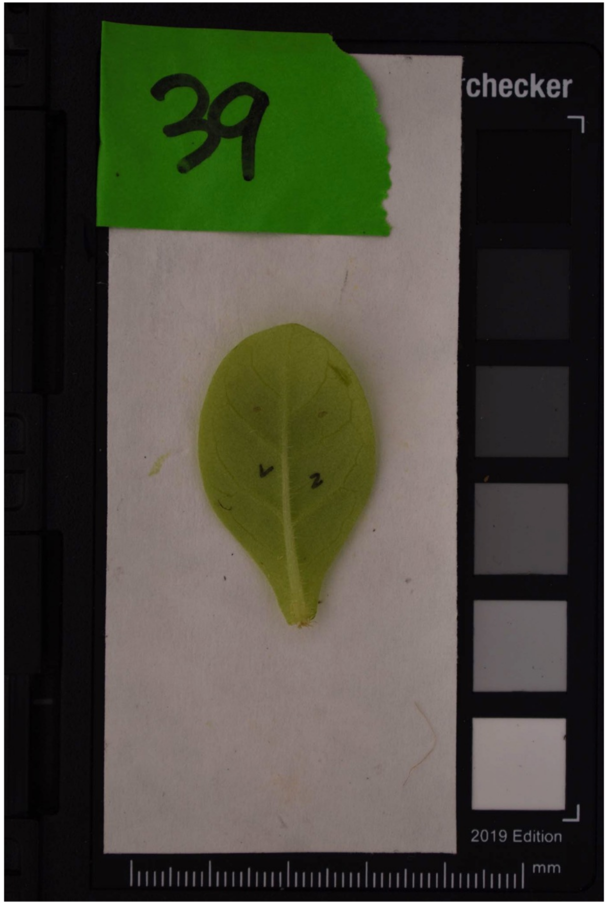
39
checker
2019 Edition
mm
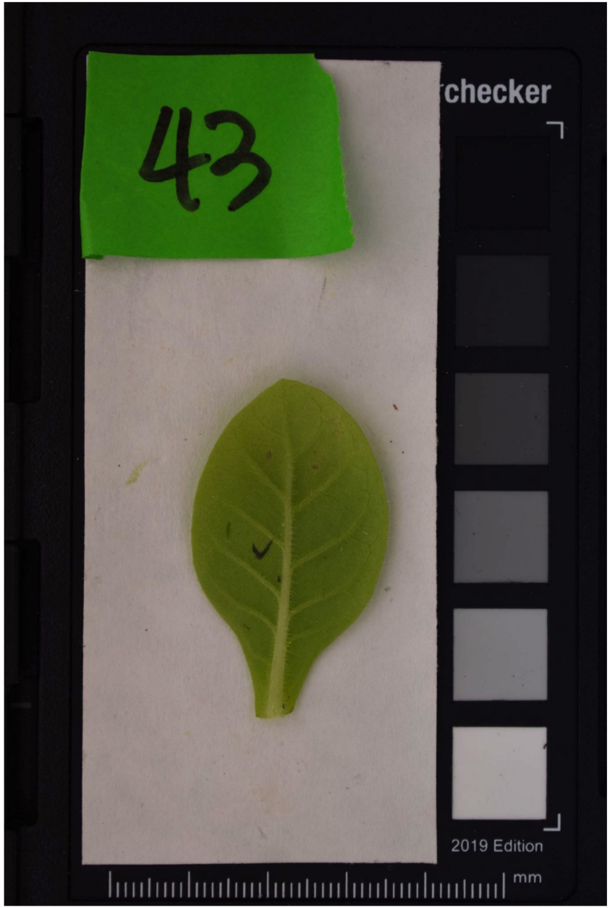
43
checker
2019 Edition
mm

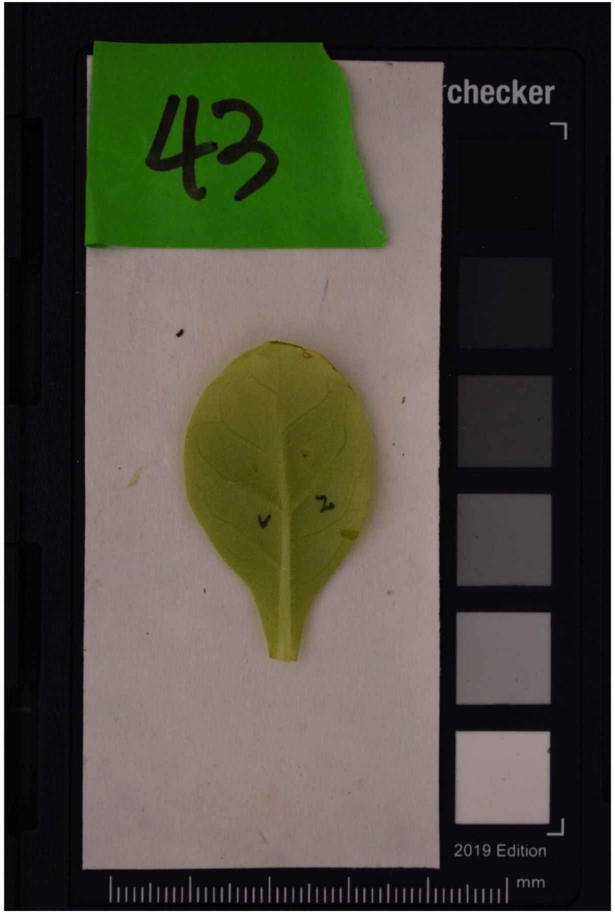
43
checker
2019 Edition
mm

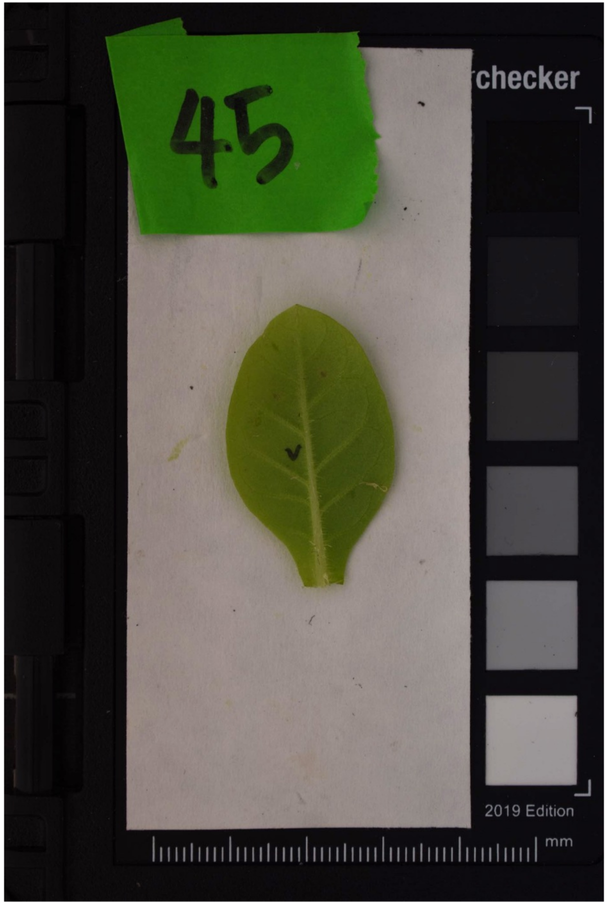
45
checker
2019 Edition
mm

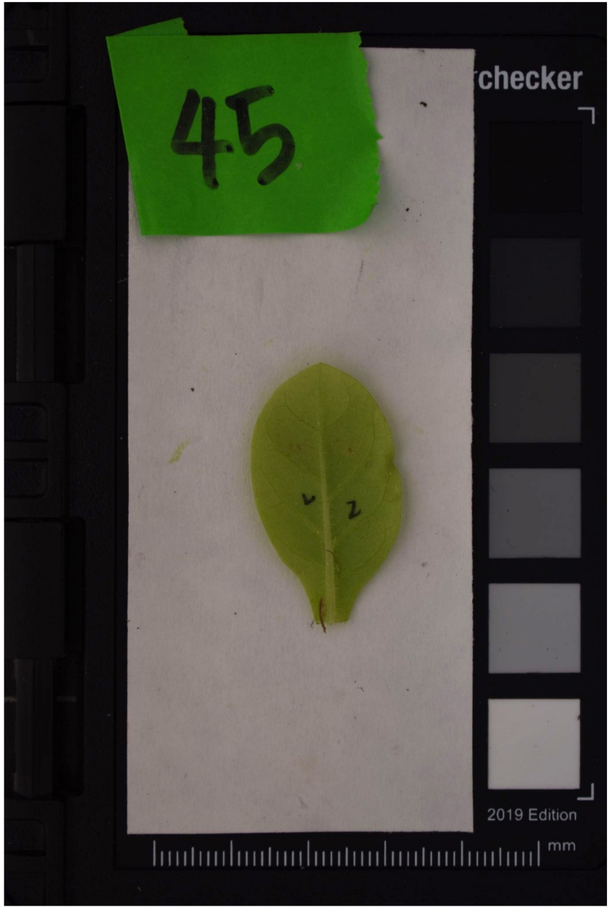
45
checker
2019 Edition
mm

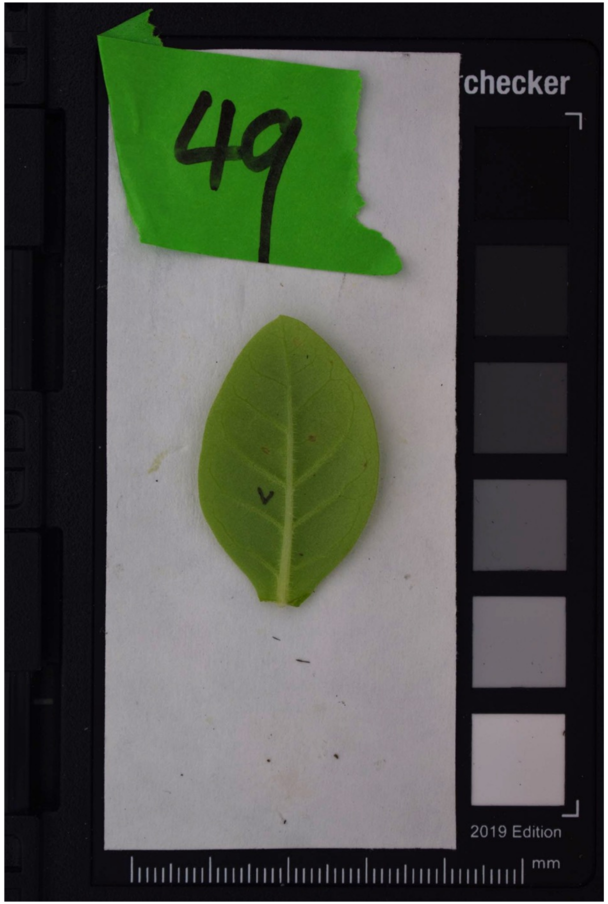
49
checker
2019 Edition
mm

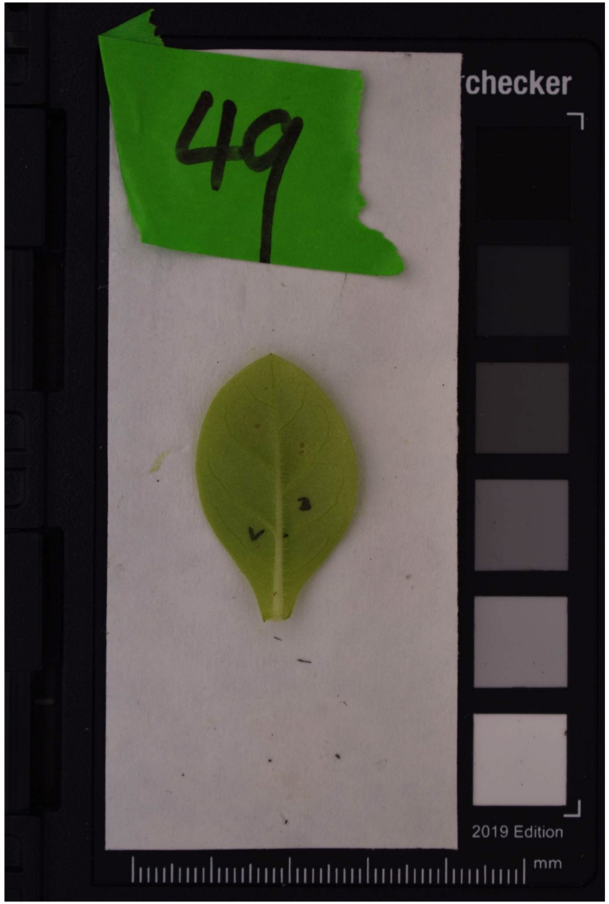
49
checker
2019 Edition
mm

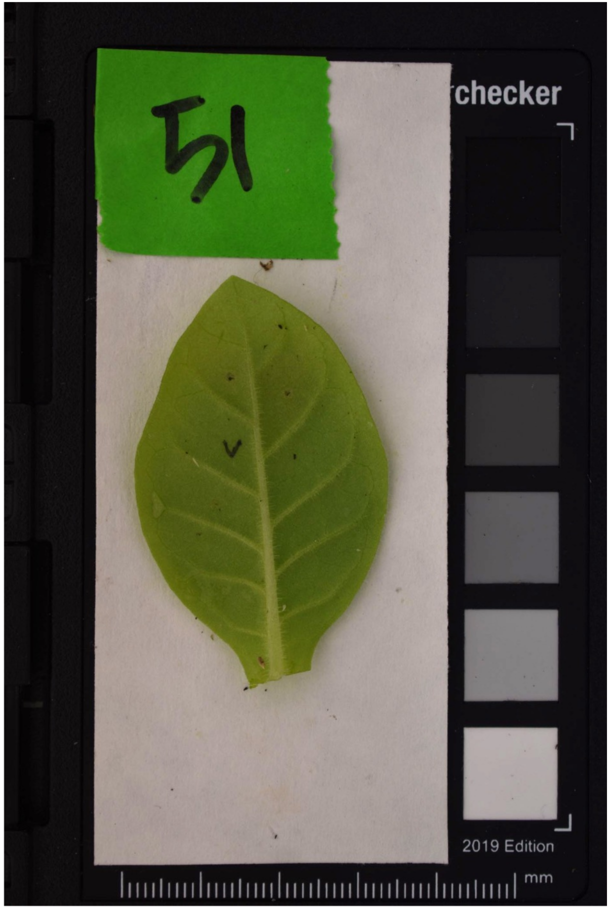
51
checker
2019 Edition
mm

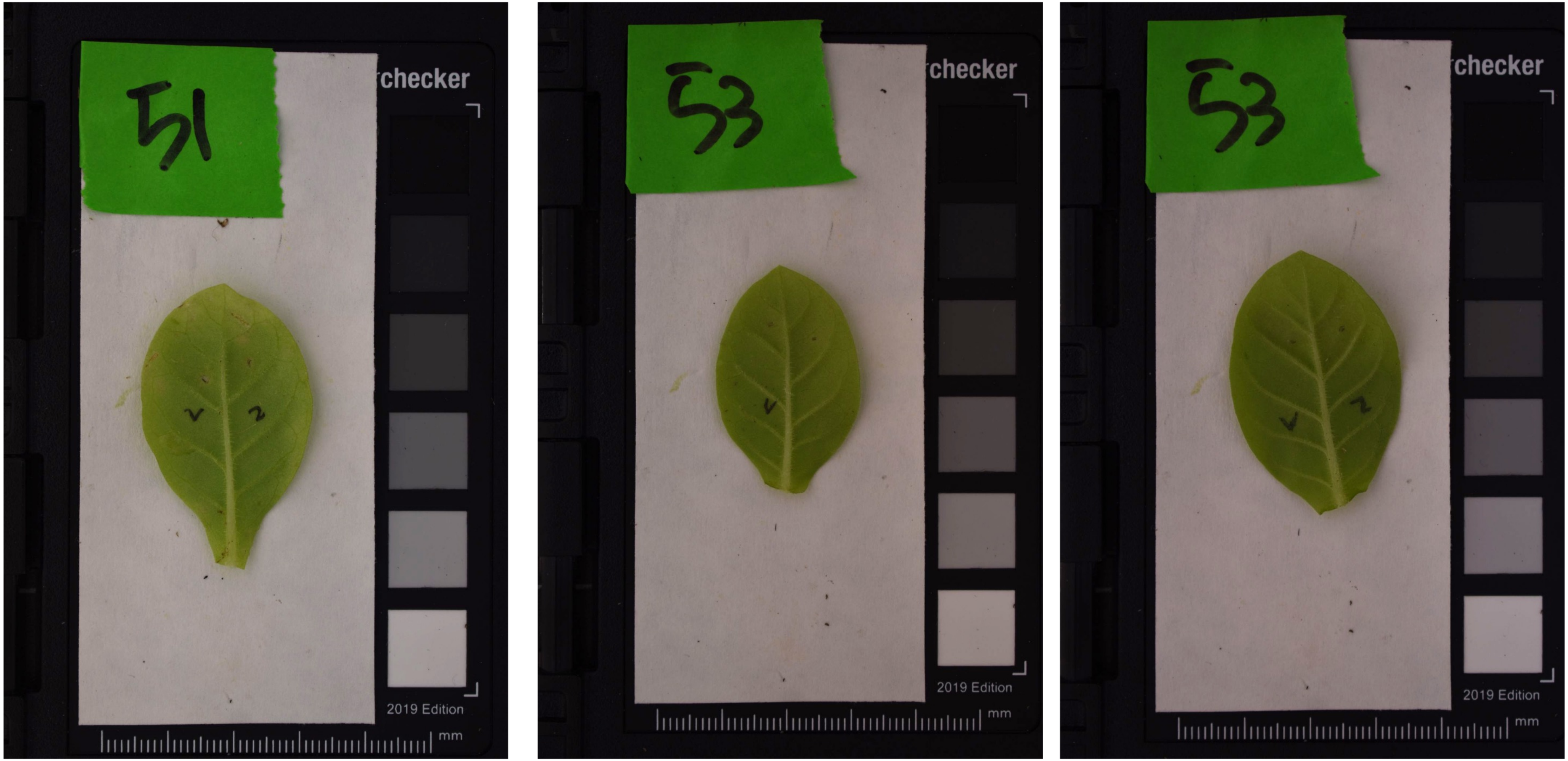
checker
2019 Edition
mm
checker
2019 Edition
mm
checker
2019 Edition
mm

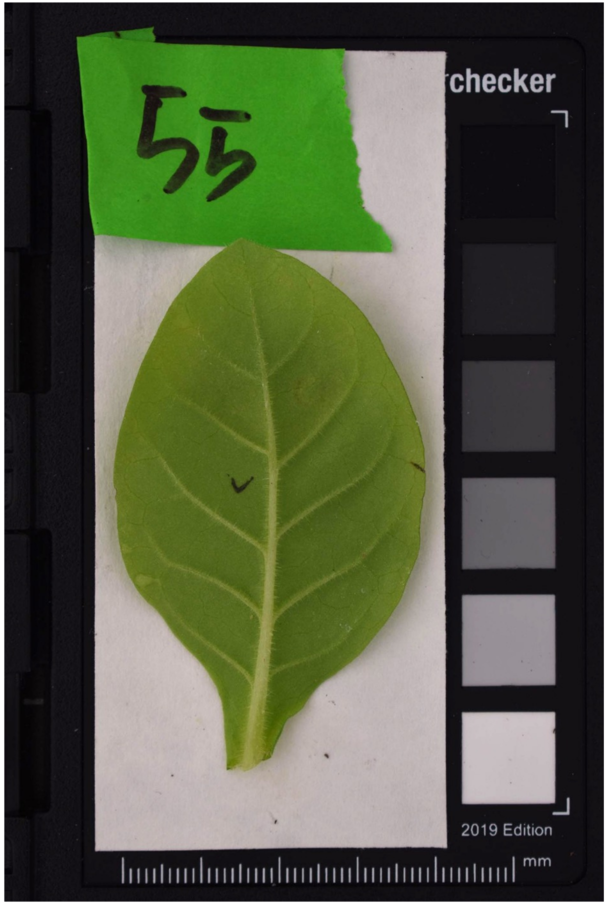
55
checker
2019 Edition
mm
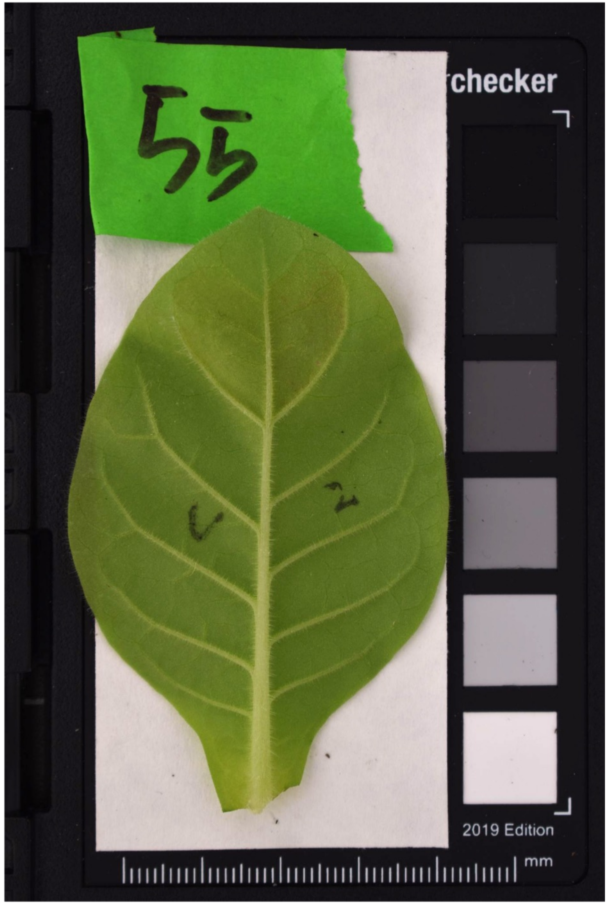
55
checker
2019 Edition
mm
