## Supplemental Figure S9.C for "Coding-sequence evolution does not explain divergence in petal anthocyanin pigmentation between *Mimulus luteus* var. *luteus* and *M. l. variegatus*"

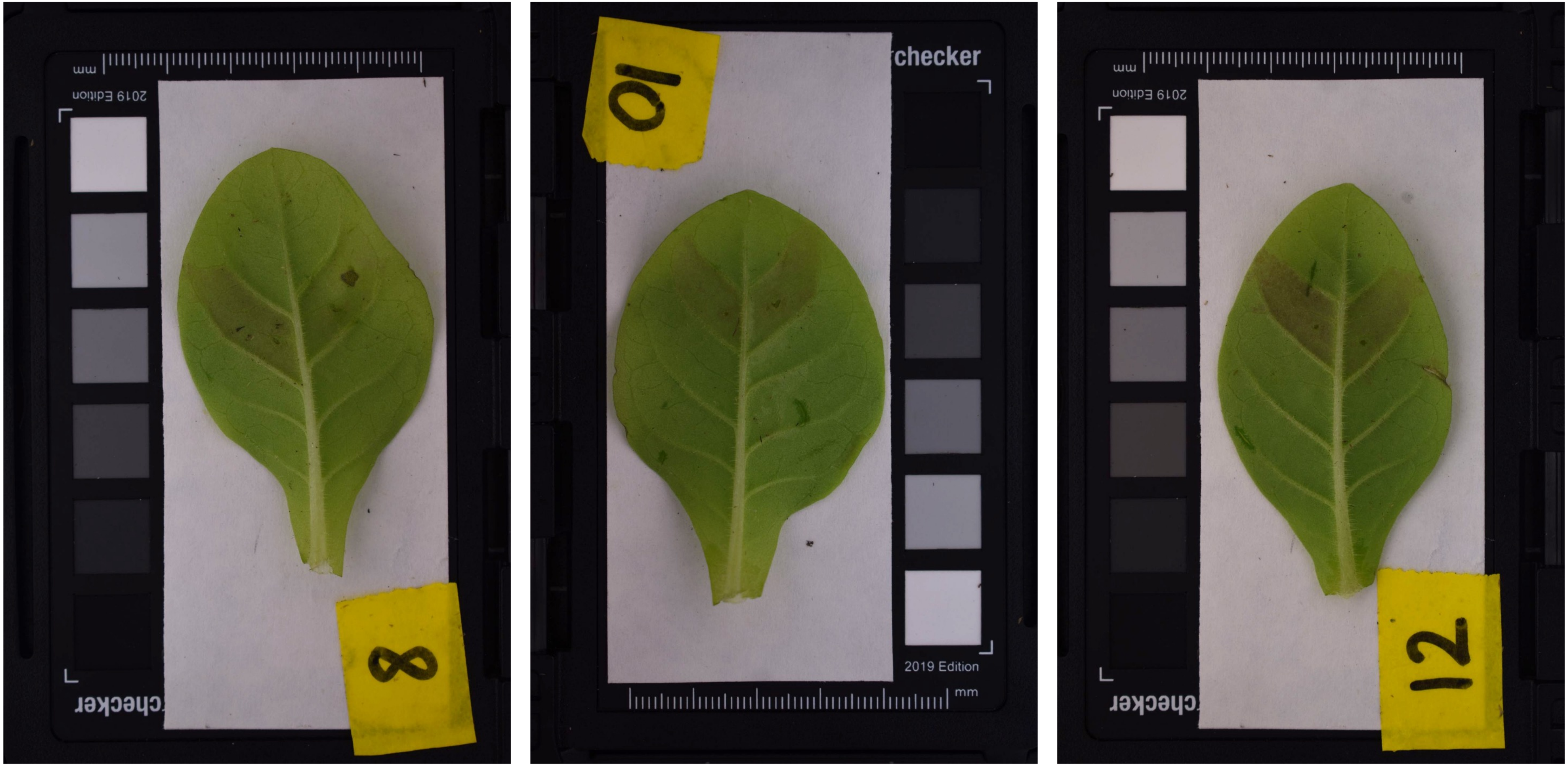
8
10
checker
2019 Edition
mm
12

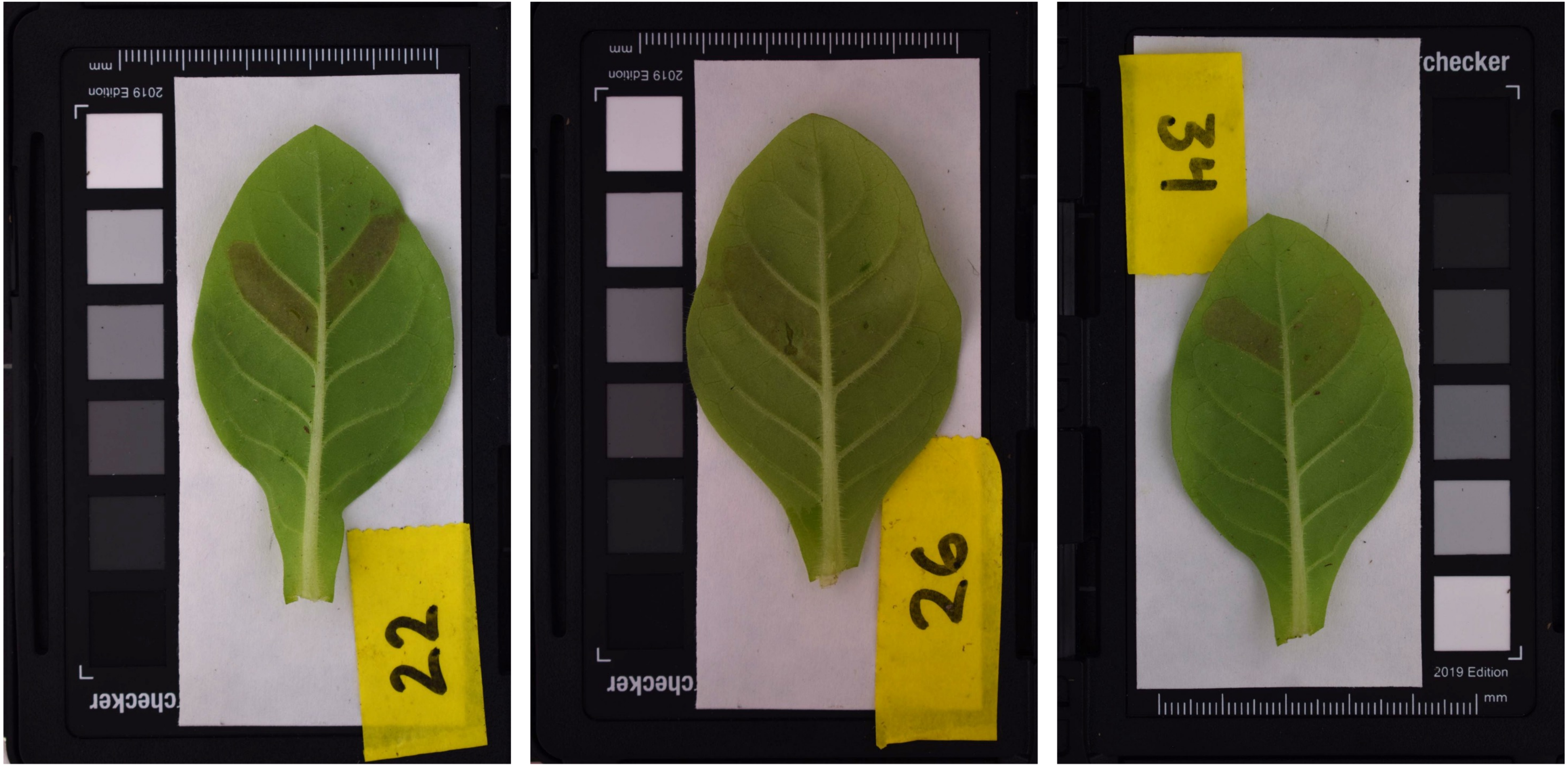
22
26
34
checker
2019 Edition
mm

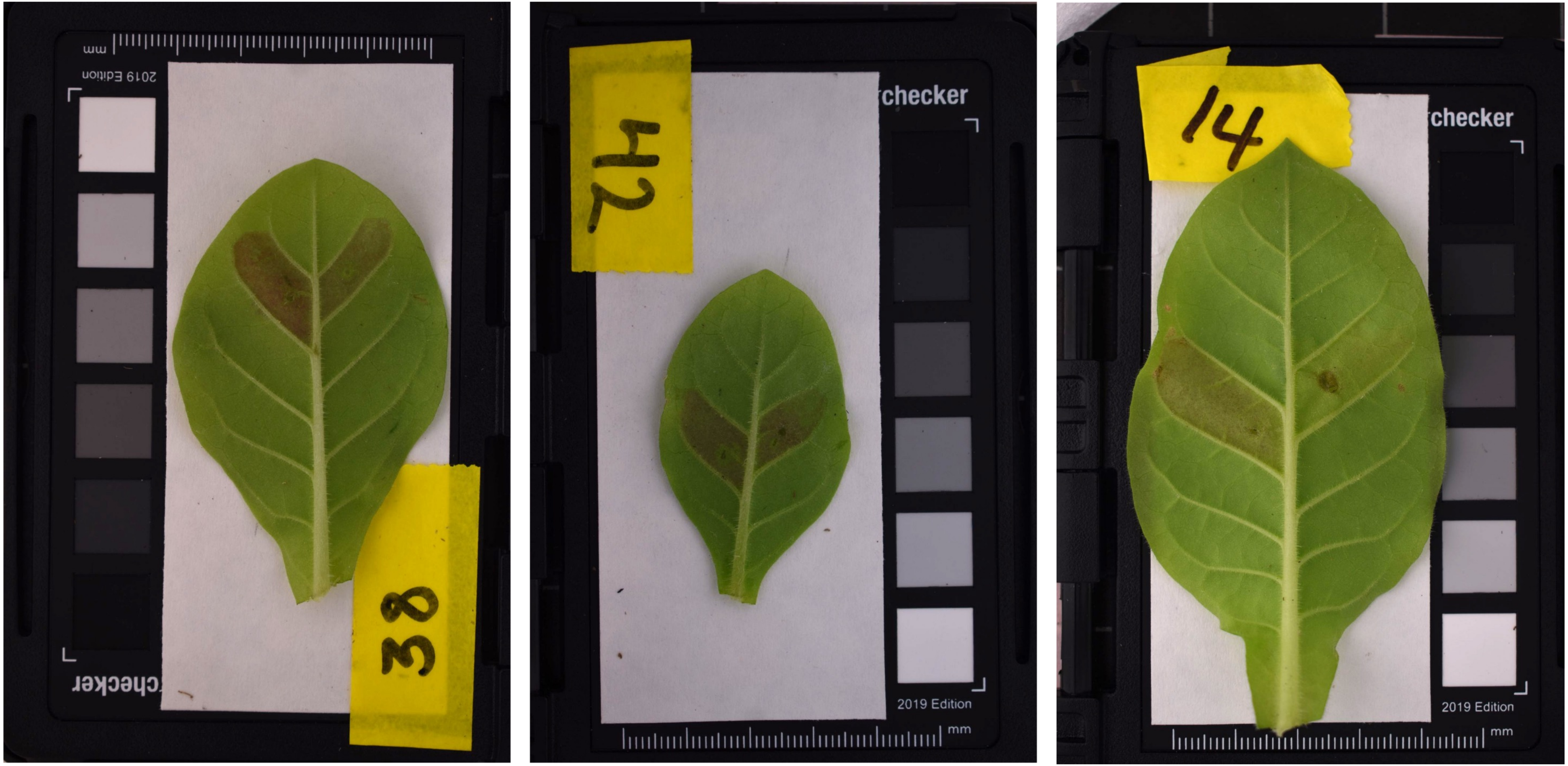
38
42
checker
2019 Edition
mm
14
checker
2019 Edition
mm

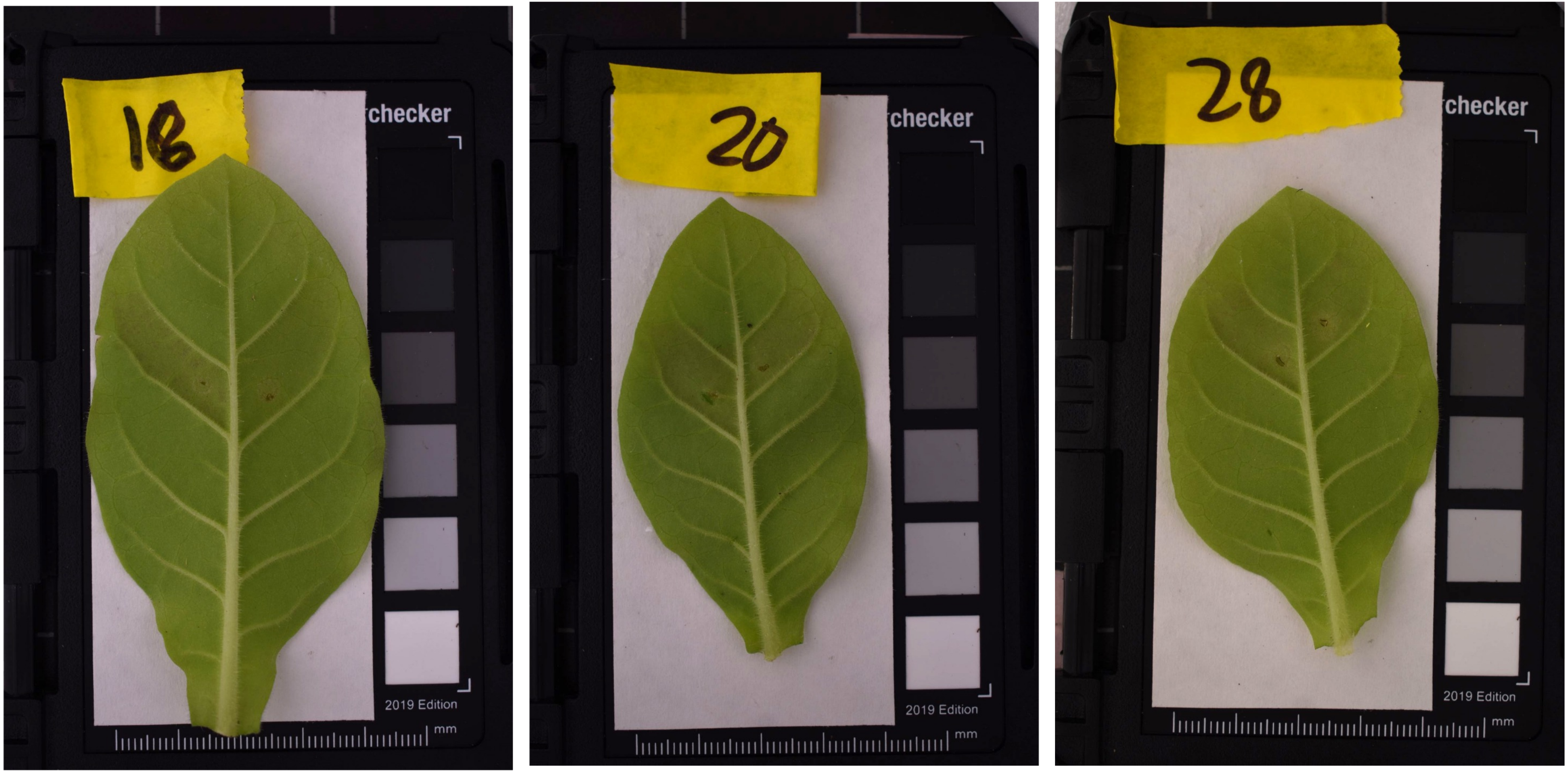
18
checker
2019 Edition
mm
20
checker
2019 Edition
mm
28
checker
2019 Edition
mm

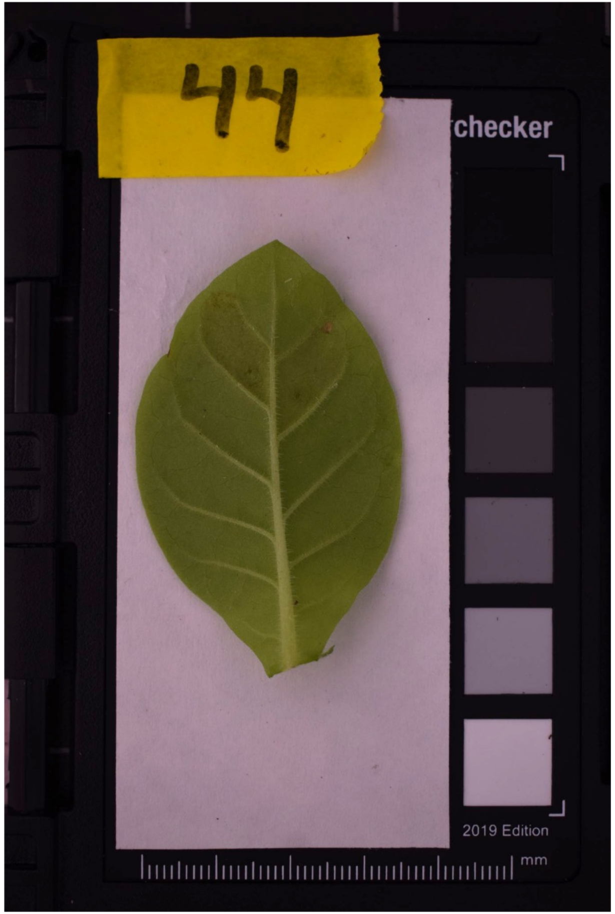
44
checker
2019 Edition
mm

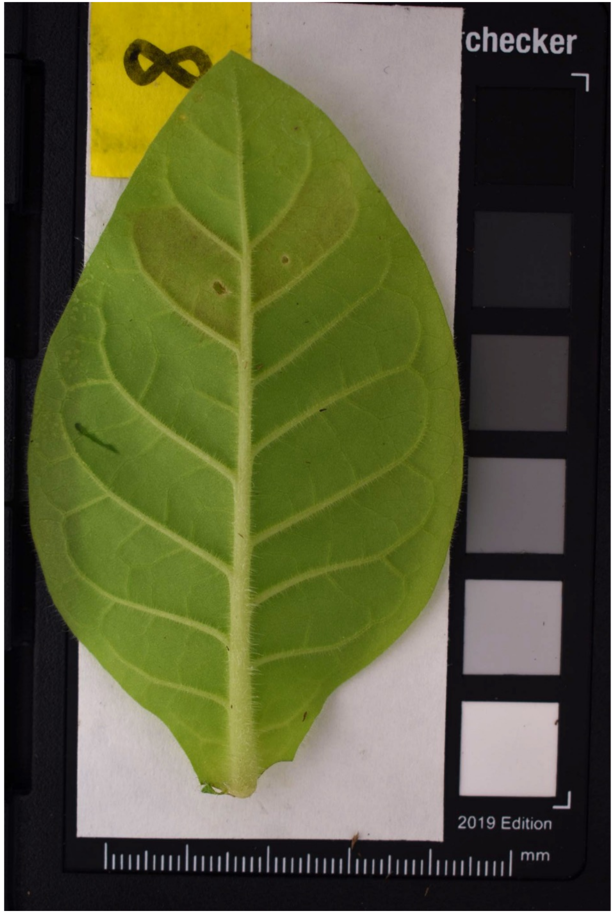
8
checker
2019 Edition
mm

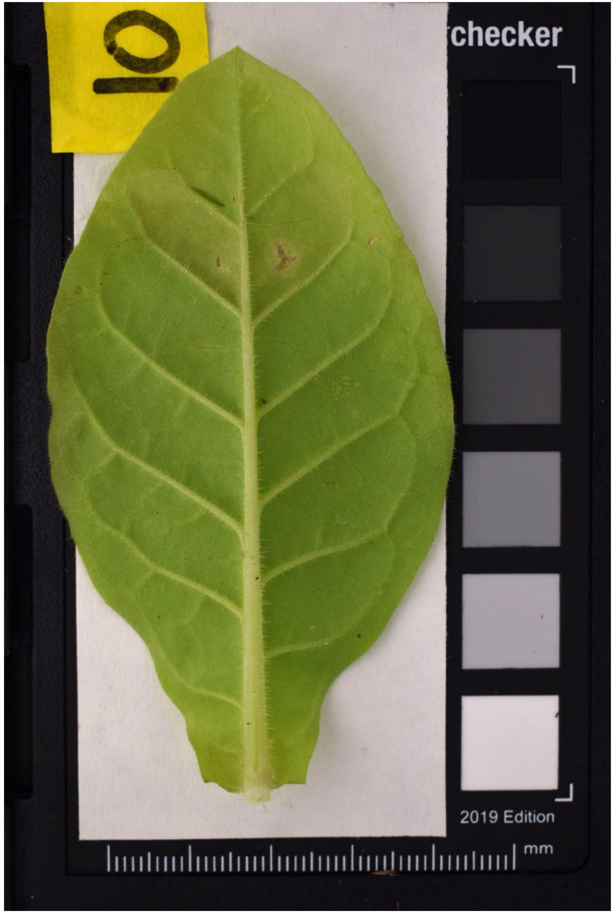
10
checker
2019 Edition
mm

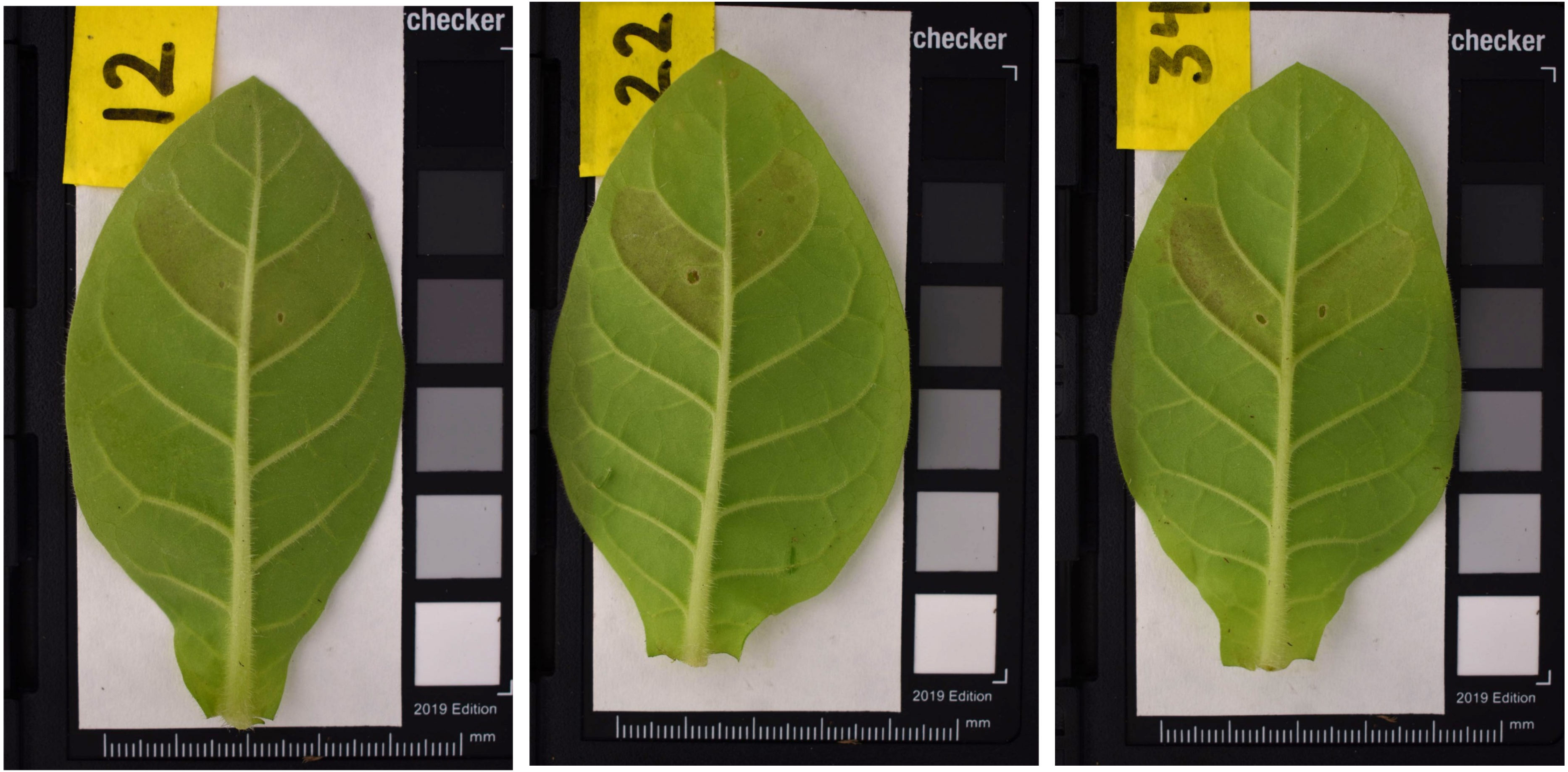
12
checker
2019 Edition
mm
22
checker
2019 Edition
mm
checker
2019 Edition
mm

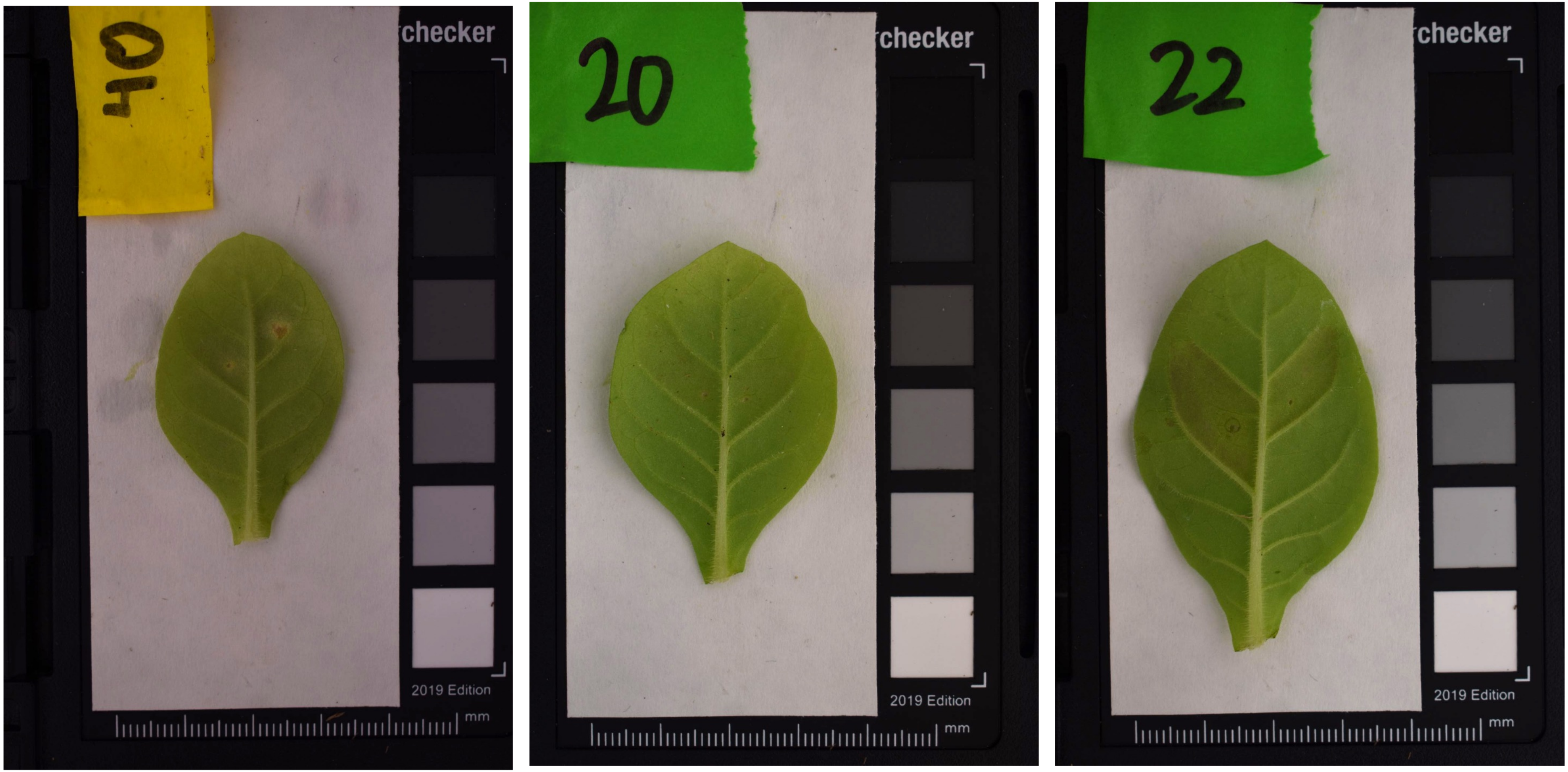
40
checker
2019 Edition
mm
20
checker
2019 Edition
mm
22
checker
2019 Edition
mm

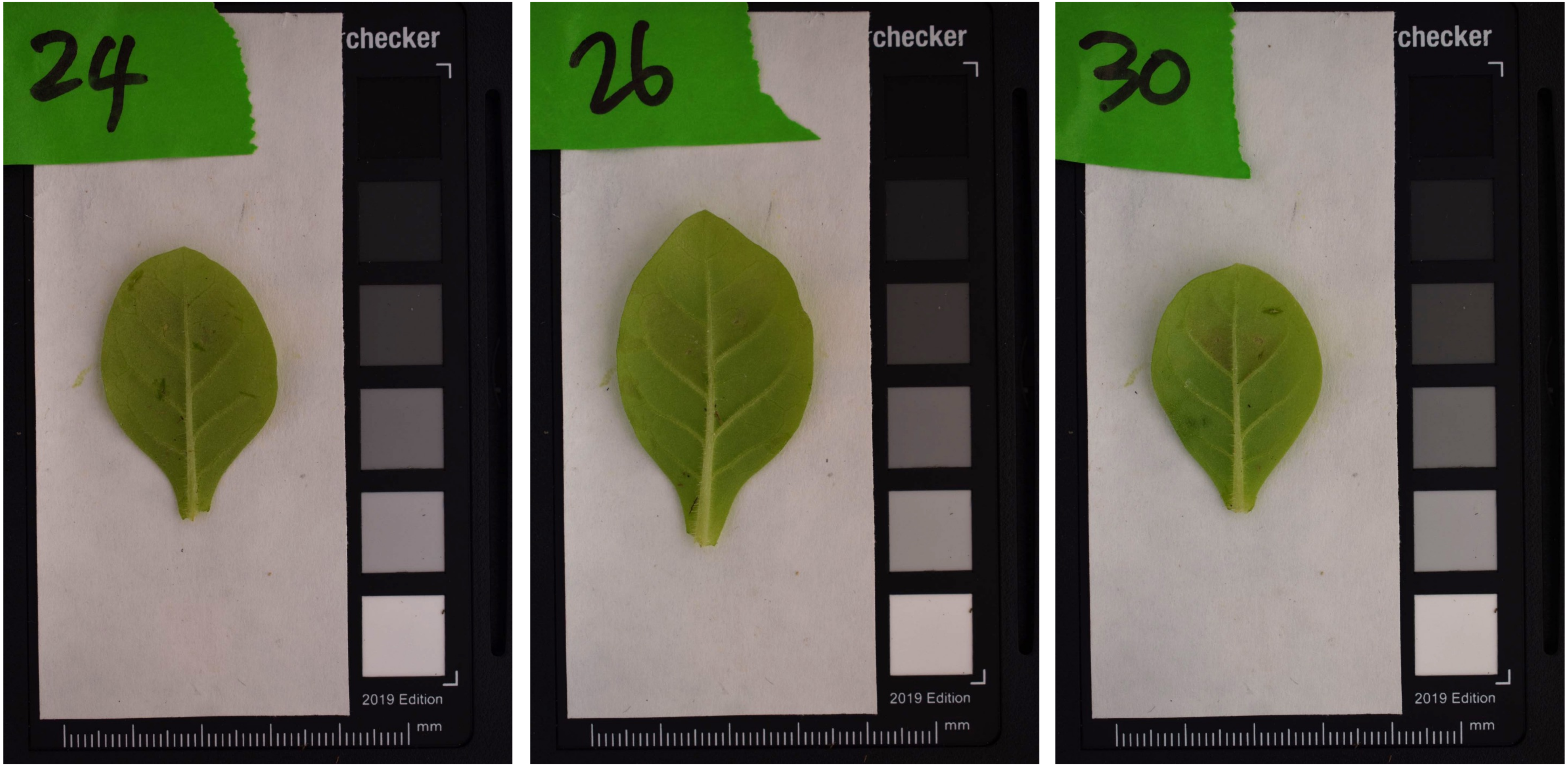
24
checker
2019 Edition
mm
26
checker
2019 Edition
mm
30
checker
2019 Edition
mm

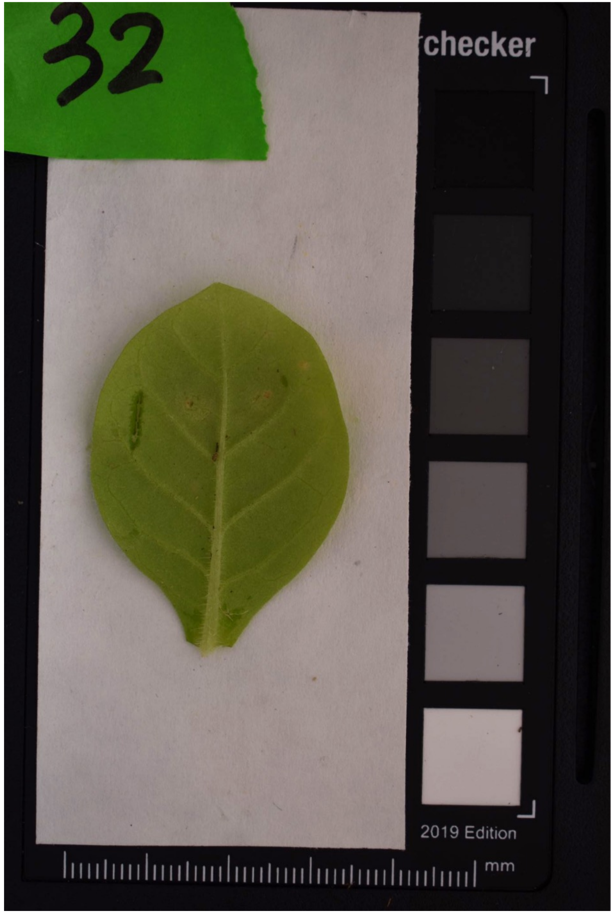
32
checker
2019 Edition
mm

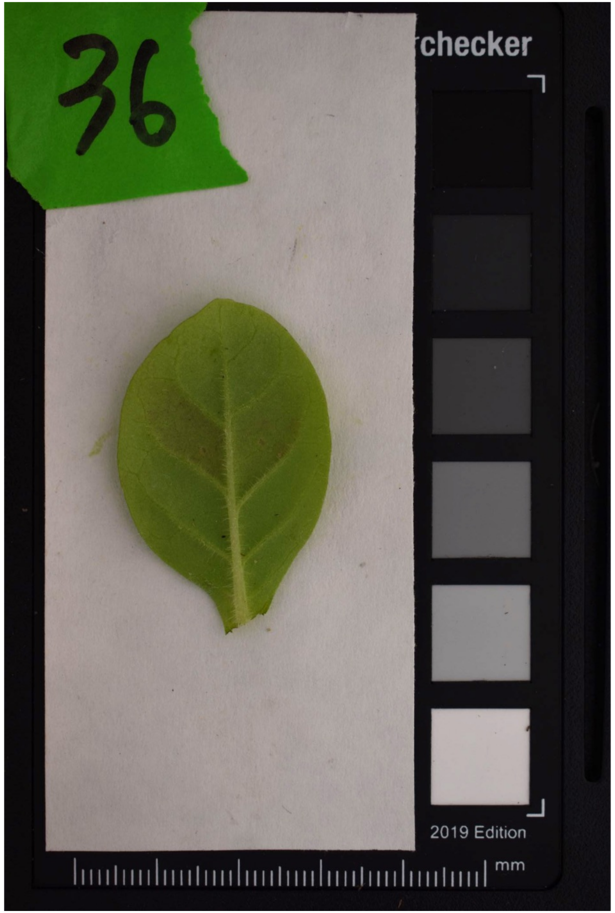
36
checker
2019 Edition
mm

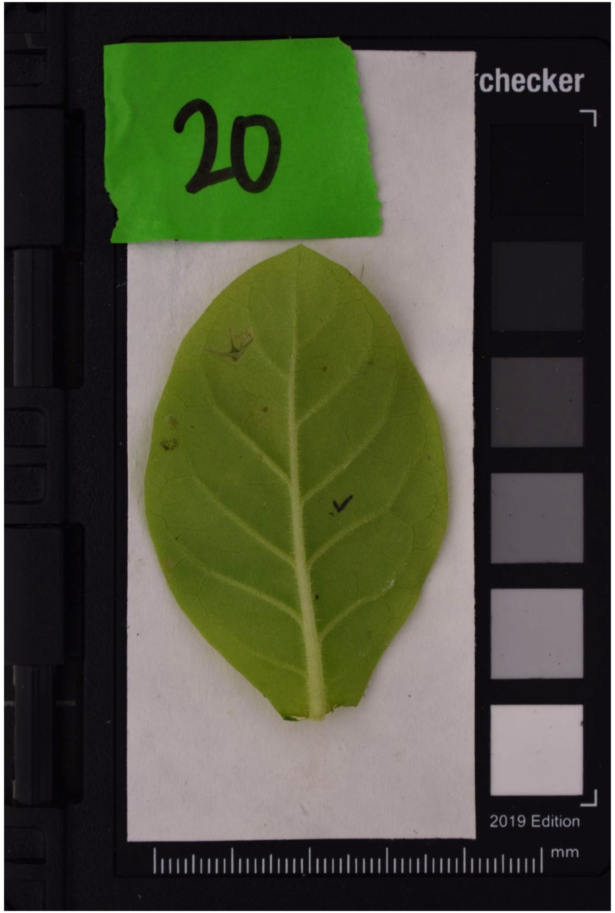
20
checker
2019 Edition
mm

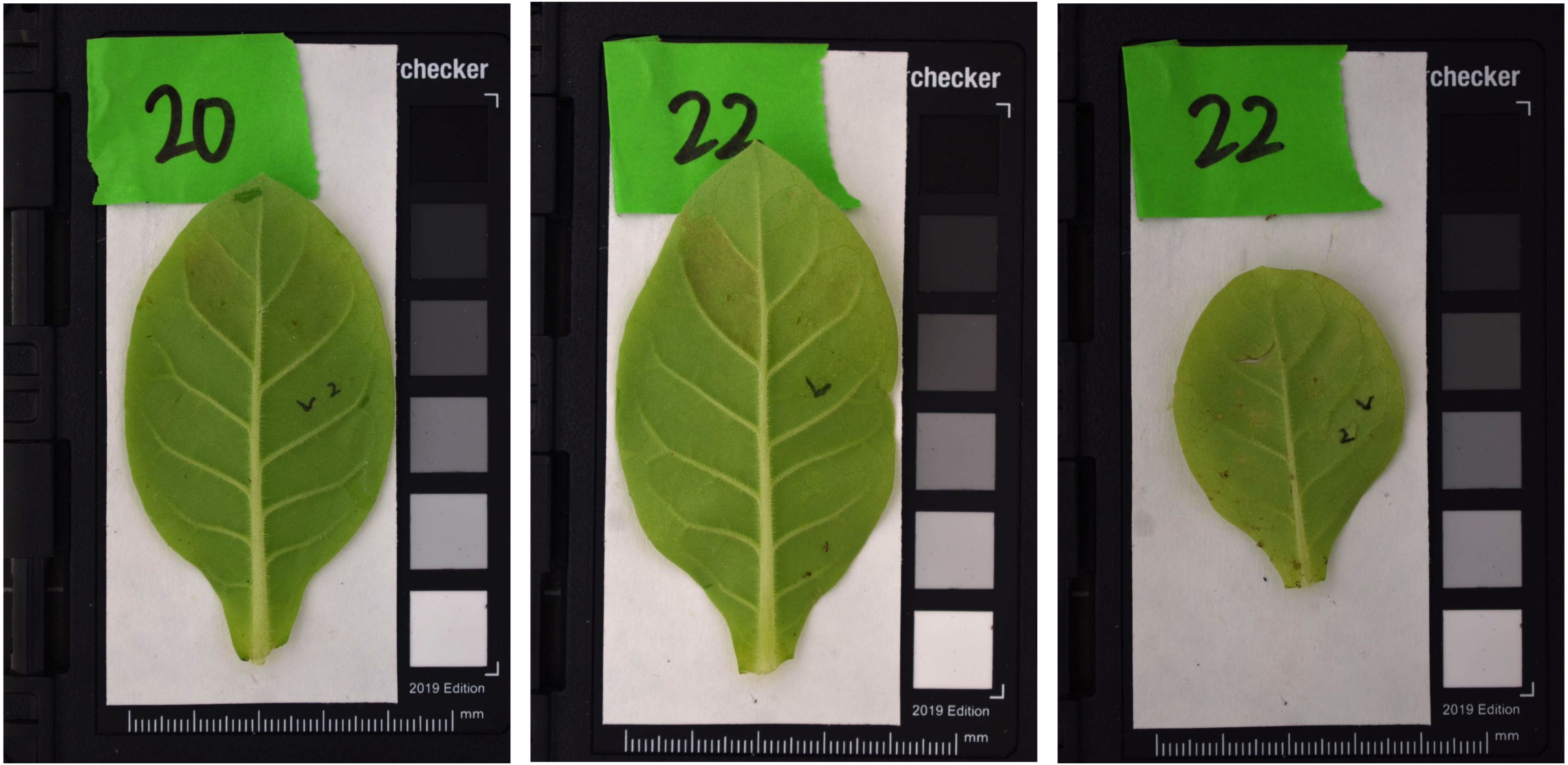
20
checker
2019 Edition
mm
22
checker
2019 Edition
mm
22
checker
2019 Edition
mm

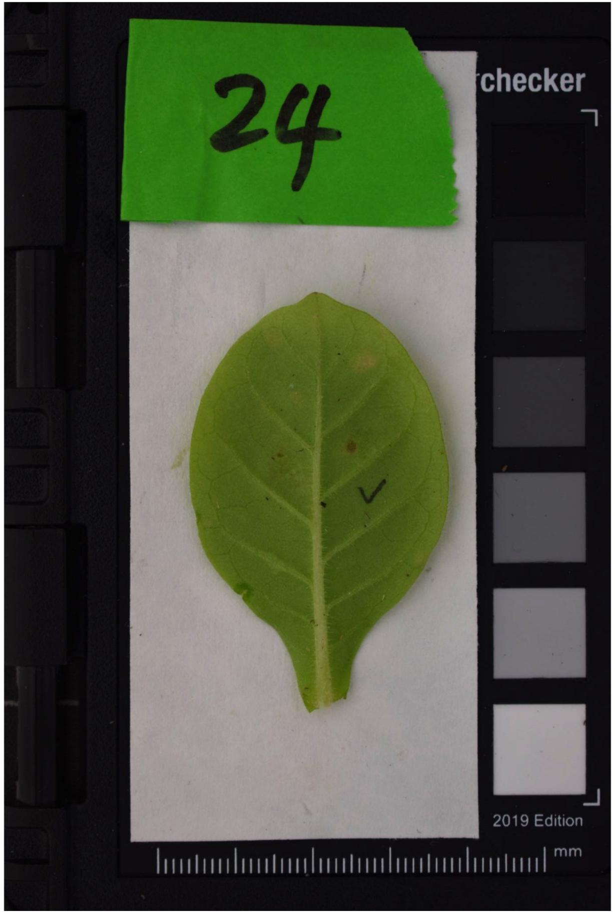
24
checker
2019 Edition
mm

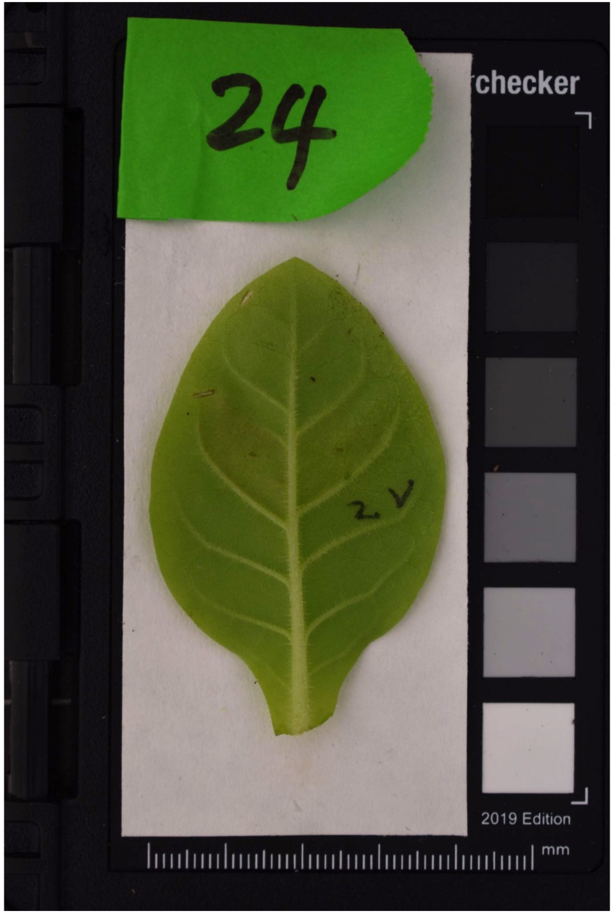
24
checker
2019 Edition
mm

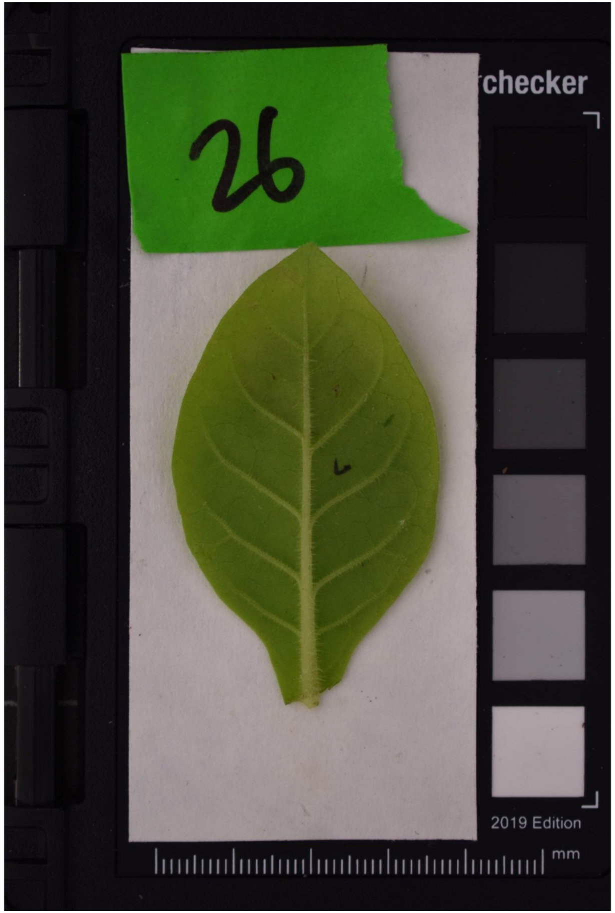
26
checker
2019 Edition
mm

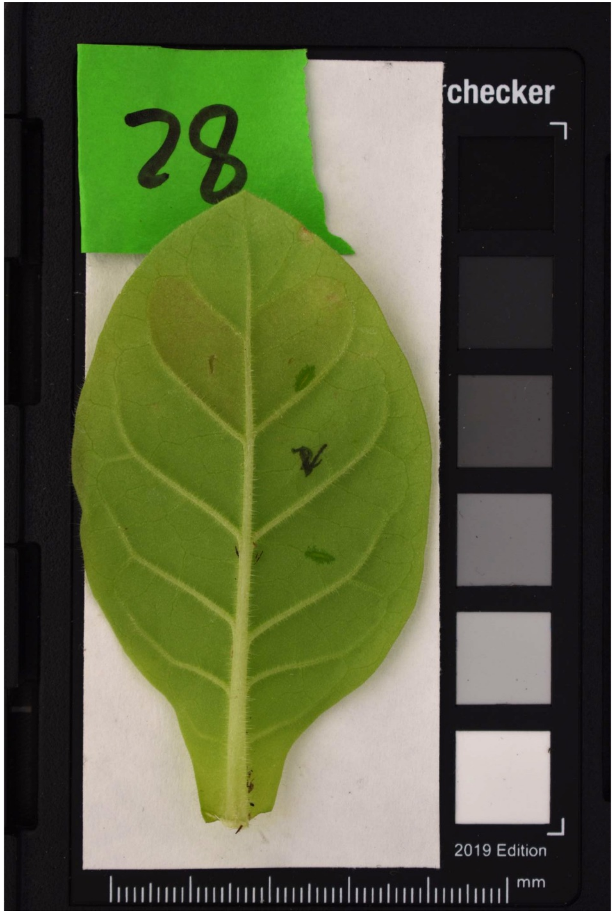
28
2019 Edition
mm

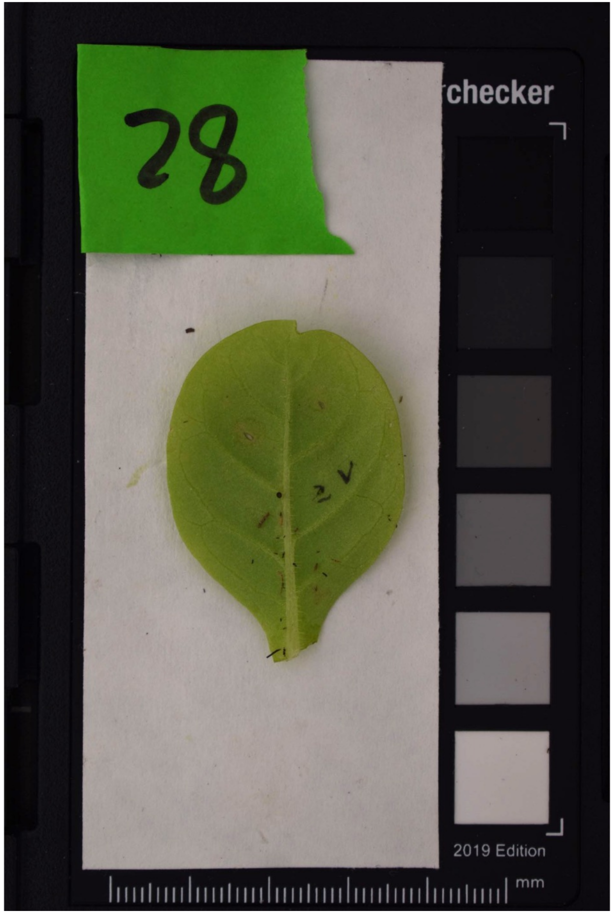
28
2019 Edition
mm

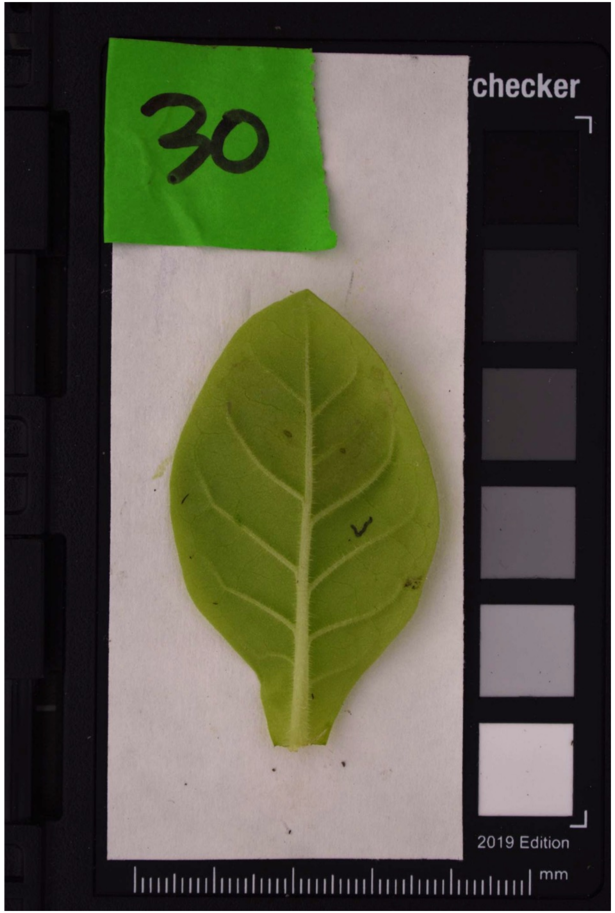
30
2019 Edition
mm

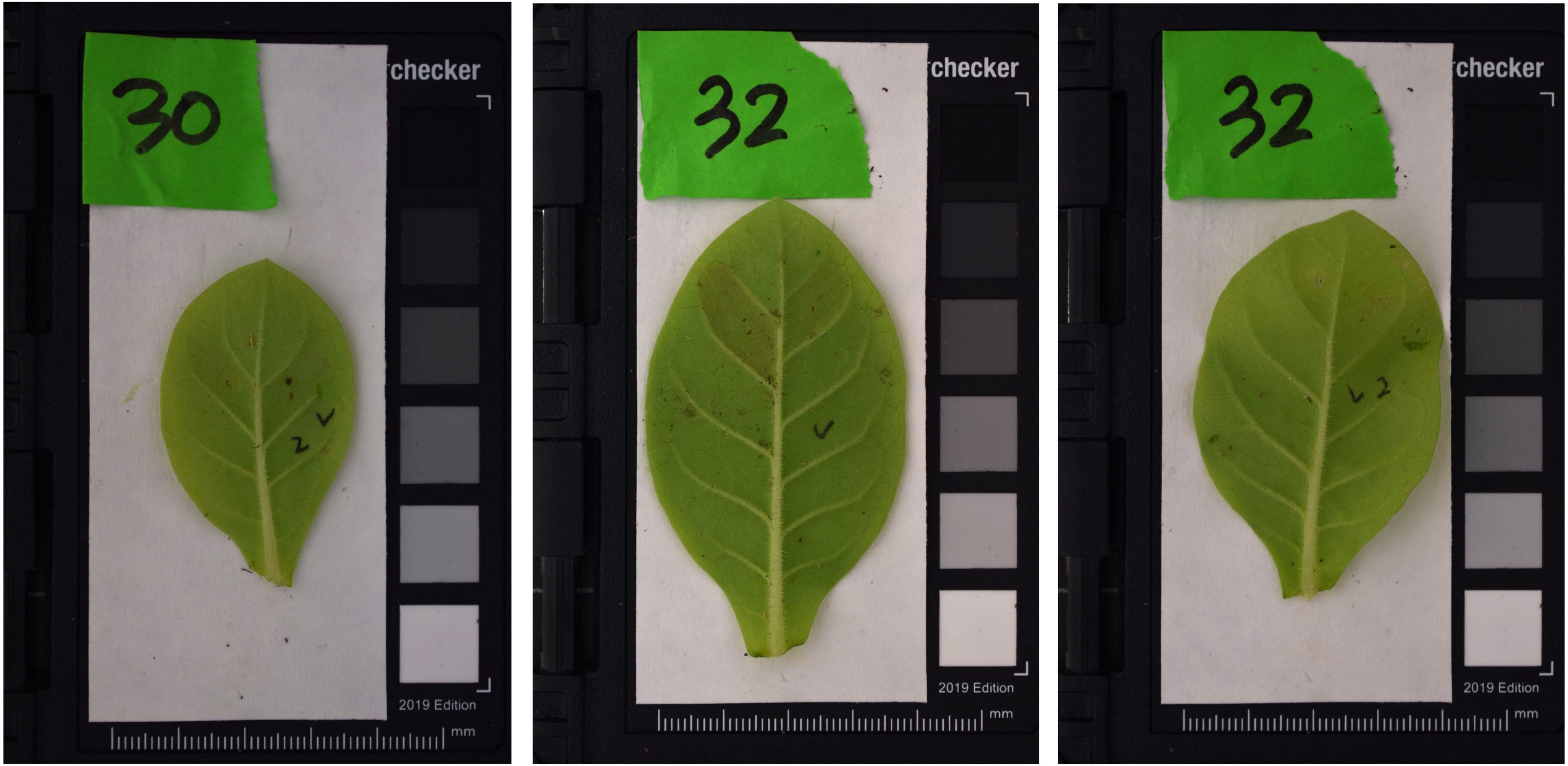
30
checker
2019 Edition
mm
32
checker
2019 Edition
mm
32
checker
2019 Edition
mm

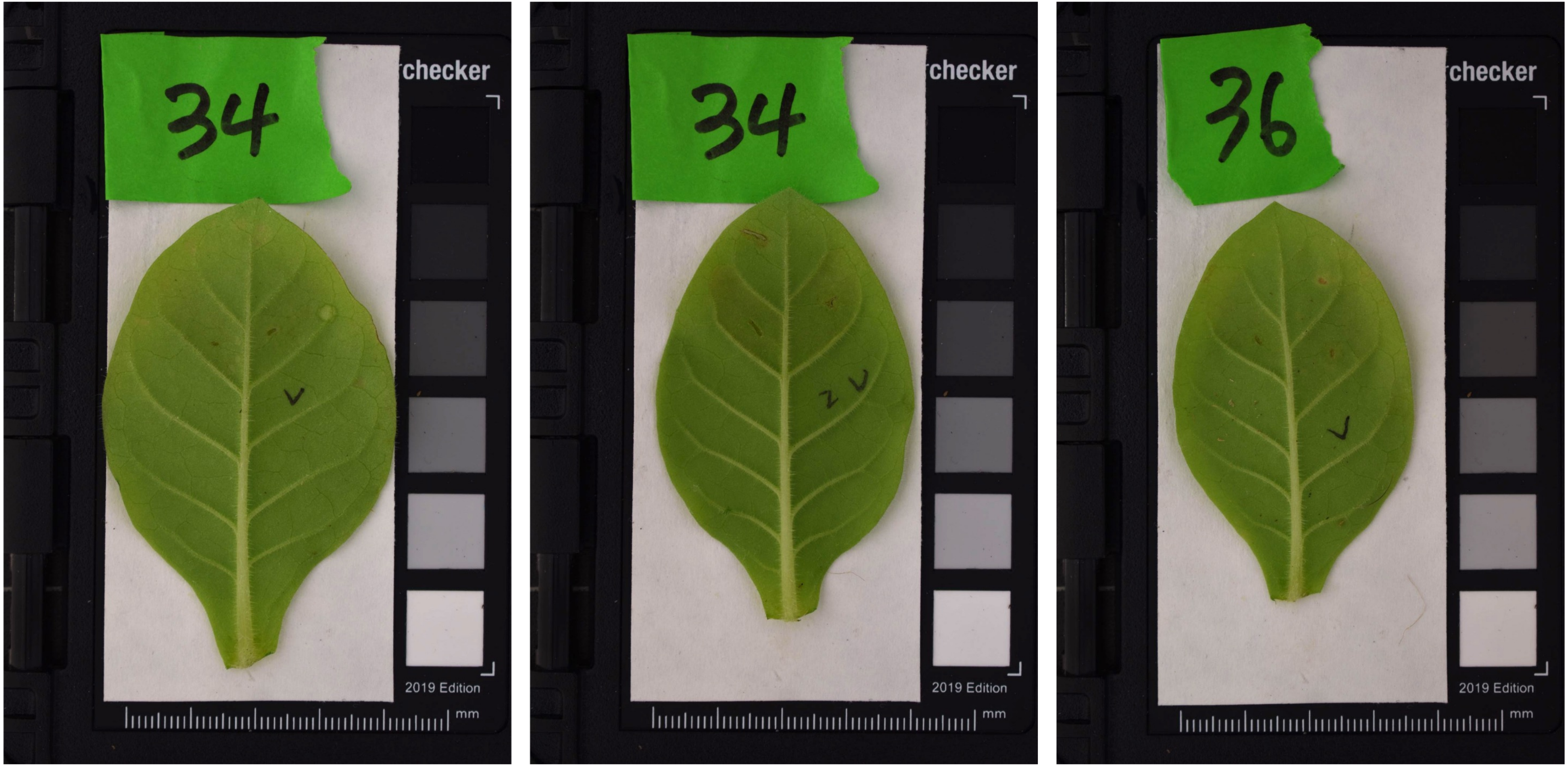
34
checker
2019 Edition
mm
34
checker
2019 Edition
mm
36
checker
2019 Edition
mm

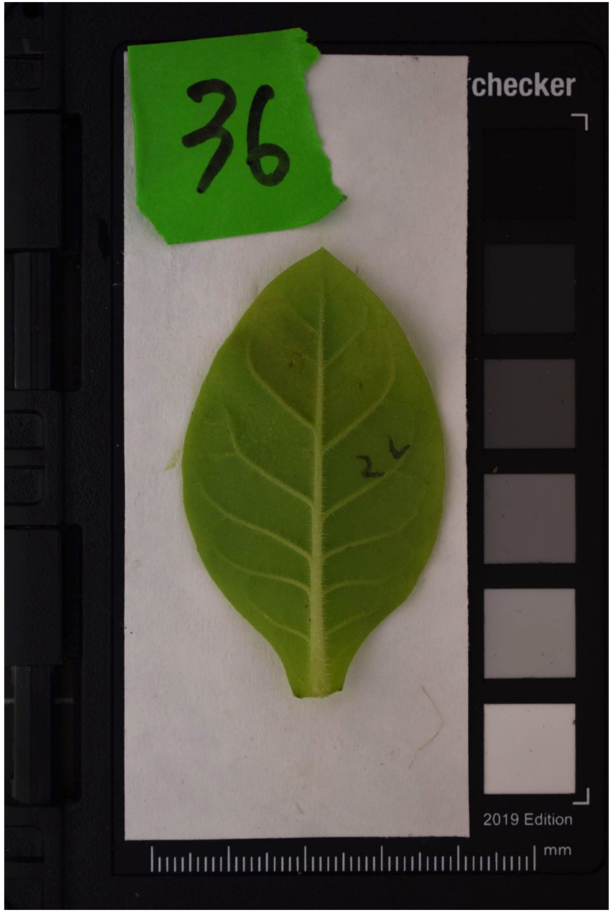
36
checker
2 v
2019 Edition
mm

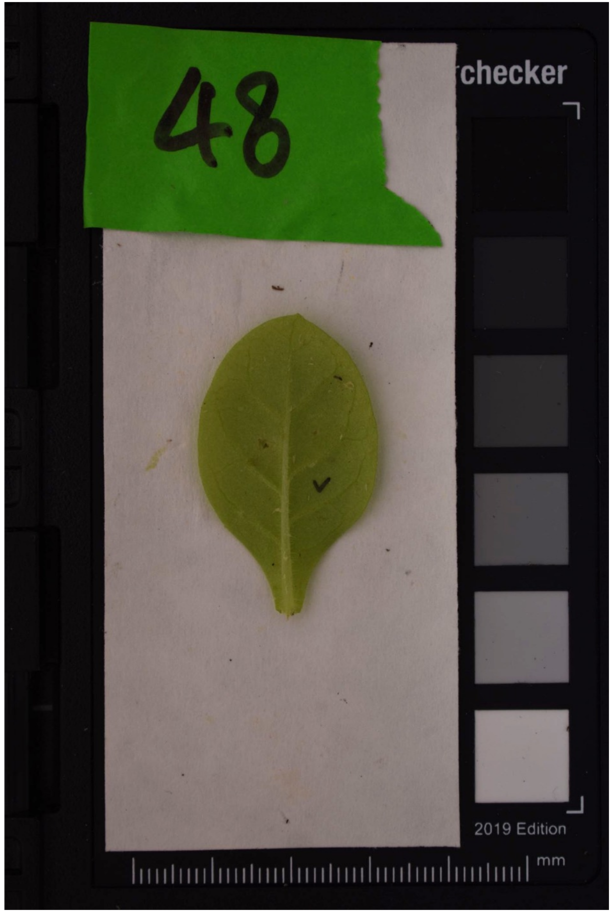
48
checker
v
2019 Edition
mm

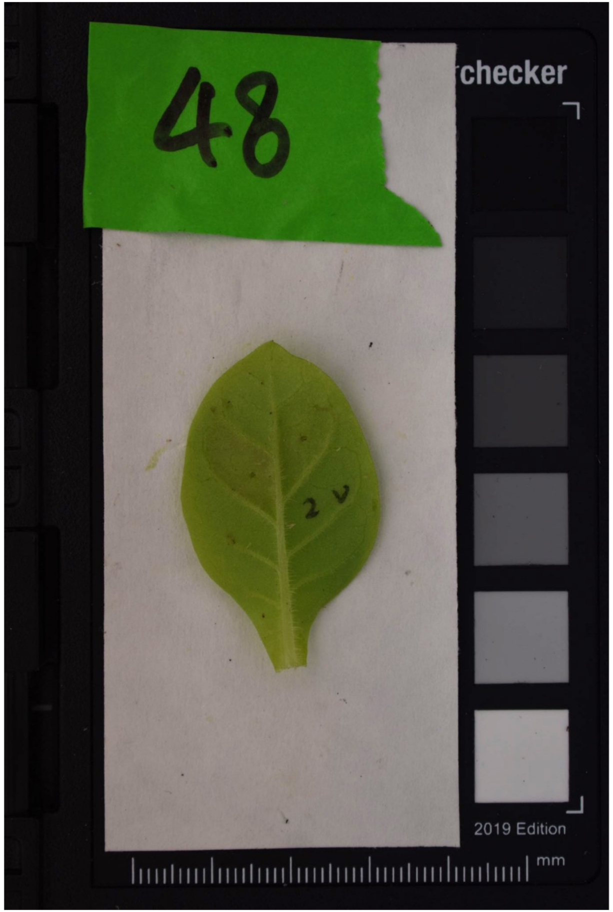
48
checker
2 v
2019 Edition
mm

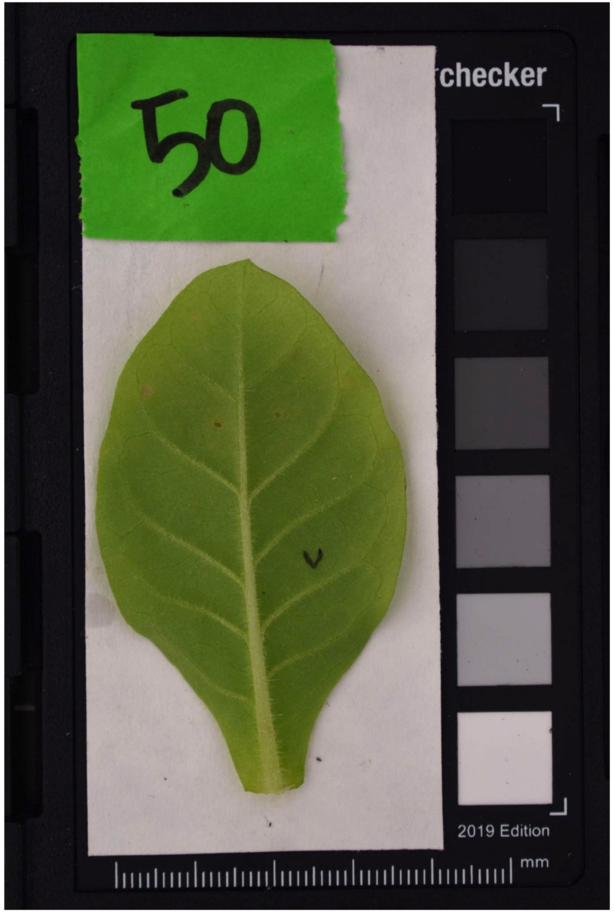
50
checker
2019 Edition
mm

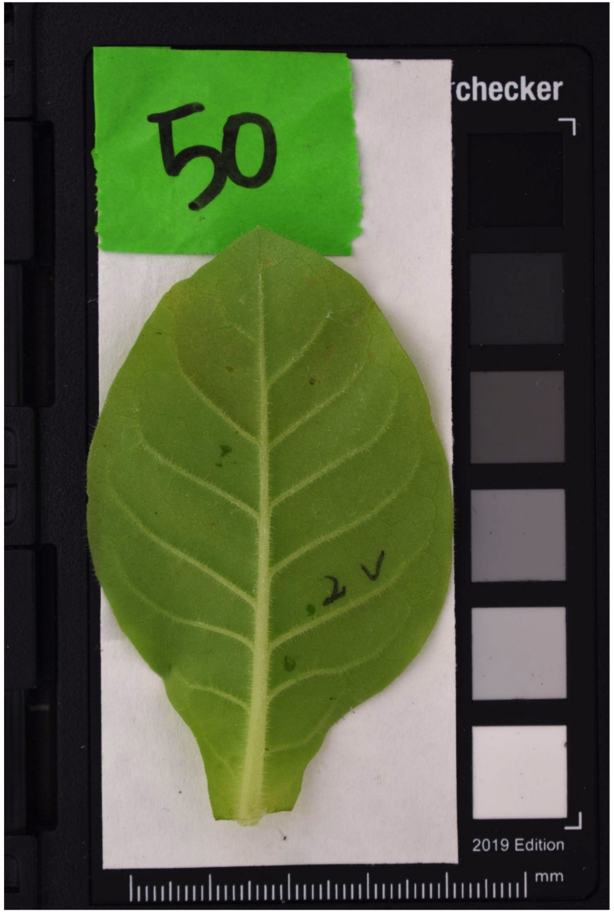
50
checker
2019 Edition
mm

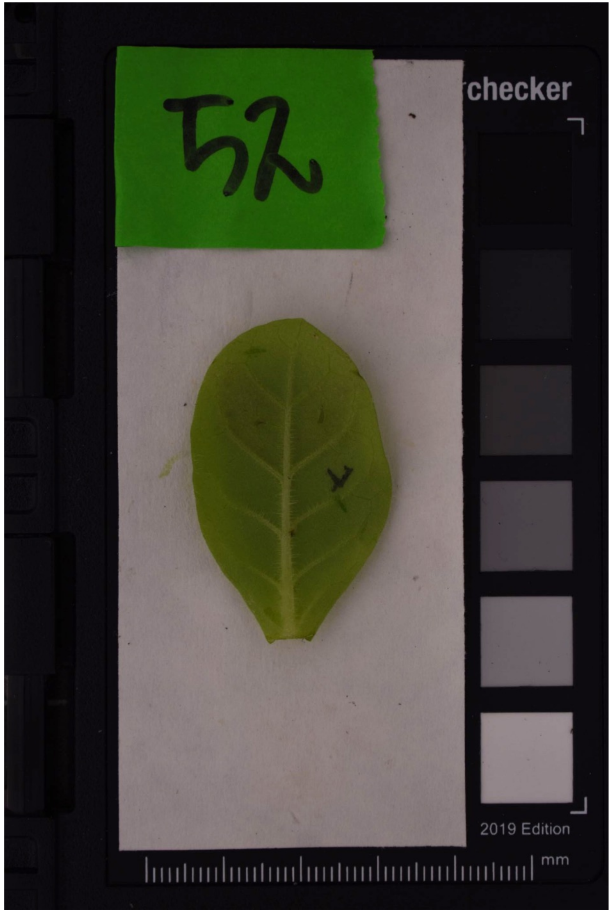
52
checker
2019 Edition
mm
